# Novel and improved *Caenorhabditis briggsae* gene models generated by community curation

**DOI:** 10.1101/2023.05.16.541014

**Authors:** Nicolas D. Moya, Lewis Stevens, Isabella R. Miller, Chloe E. Sokol, Joseph L. Galindo, Alexandra D. Bardas, Edward S. H. Koh, Justine Rozenich, Cassia Yeo, Maryanne Xu, Erik C. Andersen

## Abstract

**Background:** The nematode *Caenorhabditis briggsae* has been used as a model for genomics studies compared to *Caenorhabditis elegans* because of its striking morphological and behavioral similarities. These studies yielded numerous findings that have expanded our understanding of nematode development and evolution. However, the potential of *C. briggsae* to study nematode biology is limited by the quality of its genome resources. The reference genome and gene models for the *C. briggsae* laboratory strain AF16 have not been developed to the same extent as *C. elegans*. The recent publication of a new chromosome-level reference genome for QX1410, a *C. briggsae* wild strain closely related to AF16, has provided the first step to bridge the gap between *C. elegans* and *C. briggsae* genome resources. Currently, the QX1410 gene models consist of protein-coding gene predictions generated from short- and long-read transcriptomic data. Because of the limitations of gene prediction software, the existing gene models for QX1410 contain numerous errors in their structure and coding sequences. In this study, a team of researchers manually inspected over 21,000 software-derived gene models and underlying transcriptomic data to improve the protein-coding gene models of the *C. briggsae* QX1410 genome.

**Results:** We designed a detailed workflow to train a team of nine students to manually curate genes using RNA read alignments and predicted gene models. We manually inspected the gene models using the genome annotation editor, Apollo, and proposed corrections to the coding sequences of over 8,000 genes. Additionally, we modeled thousands of putative isoforms and untranslated regions. We exploited the conservation of protein sequence length between *C. briggsae* and *C. elegans* to quantify the improvement in protein-coding gene model quality before and after curation. Manual curation led to a substantial improvement in the protein sequence length accuracy of QX1410 genes. We also compared the curated QX1410 gene models against the existing AF16 gene models. The manual curation efforts yielded QX1410 gene models that are similar in quality to the extensively curated AF16 gene models in terms of protein-length accuracy and biological completeness scores. Collinear alignment analysis between the QX1410 and AF16 genomes revealed over 1,800 genes affected by spurious duplications and inversions in the AF16 genome that are now resolved in the QX1410 genome.

**Conclusions:** Community-based, manual curation using transcriptome data is an effective approach to improve the quality of software-derived protein-coding genes. Comparative genomic analysis using a related species with high-quality reference genome(s) and gene models can be used to quantify improvements in gene model quality in a newly sequenced genome. The detailed protocols provided in this work can be useful for future large-scale manual curation projects in other species. The chromosome-level reference genome for the *C. briggsae* strain QX1410 far surpasses the quality of the genome of the laboratory strain AF16, and our manual curation efforts have brought the QX1410 gene models to a comparable level of quality to the previous reference, AF16. The improved genome resources for *C. briggsae* provide reliable tools for the study of *Caenorhabditis* biology and other related nematodes.

## Background

The undisputed popularity of the free-living nematode *Caenorhabditis elegans* as a highly tractable model organism is facilitated by the vast collection of genetic and genomic resources that have been arduously maintained and improved by the research community. *C. elegans* possesses one of the highest quality metazoan genome assemblies with extensively validated models of its protein-coding genes. Similar efforts to generate and improve genomic resources for other species in the *Caenorhabditis* genus have enabled comparative studies that extended our understanding of genetics, development, and evolution [1–5]. The nematode *Caenorhabditis briggsae* has been a major focus of such comparative studies because of its striking morphological and behavioral similarities to *C. elegans*. Both species reproduce primarily by self-fertilization, have a nearly identical body plan, are globally distributed, and share similar ecology [6–8]. Conversely, genomic studies have shown that *C. briggsae* populations are stratified into distinct phylogeographic groups and maintain higher genetic diversity than *C. elegans* [9]. Despite the usefulness of *C. briggsae* to study *Caenorhabditis* biology, its genomic resources have not been developed to the same extent as *C. elegans*. The draft genome assembly of the *C. briggsae* laboratory strain, AF16, was first assembled in 2003 using a combination of whole-genome shotgun sequencing and a physical map based on fosmid and bacterial artificial chromosome libraries [10]. To date, the AF16 genome remains highly fragmented, with thousands of unresolved gaps and misoriented sequences [5, 11]. Moreover, unlike the gene models for the *C. elegans* laboratory strain N2, the AF16 gene models have not been extensively investigated and likely possess numerous structural and coding sequence errors. Although efforts to identify errors in the AF16 gene models have led to substantial improvements in accuracy and completeness, many loci remain inaccurate or cannot be corrected because of inconsistencies in the genome assembly [12].

Recently, advancements in long-read sequencing technologies offered by Oxford Nanopore and Pacific Biosciences have enabled new techniques to assemble highly contiguous genomes. Moreover, long-read sequencing has enabled the assembly of full-length mRNA transcripts used to accurately model protein-coding genes [13]. In a previous study, we introduced a new high-quality genome assembly for the *C. briggsae* strain QX1410, a new reference strain closely related to AF16 [5]. This newly generated genome assembly features chromosomally resolved contigs, defined chromosomal domains, and few unresolved gaps. Additionally, we generated preliminary protein-coding gene models using gene prediction tools that leverage short- and long-read RNA sequencing data. Here, we focused on the identification and repair of errors caused by automated gene predictions in the new *C. briggsae* reference strain, QX1410. To accomplish this large manual curation task, we assembled a team of nine high school, college, and graduate students. This team was trained to interpret transcriptomic data and interact with the Apollo genomic annotation editor [14], which enables easy viewing of primary transcriptomic data, along with interpretation and curation of gene models. We designed an extensive protocol with step-by-step workflows to identify and repair the most common errors found in automated gene predictions using RNA alignments. In the span of one year, we manually curated and revised over 21,000 loci to provide improved protein-coding gene models for the new *C. briggsae* reference genome. We exploited the high-quality gene models of the *C. elegans* laboratory strain N2 and the high level of protein sequence conservation between *C. elegans* and *C. briggsae* to assess the quality of the protein-gene models in the QX1410 and AF16 strains.

## Results

### Short- and long-read RNA alignments can be used to correct structural errors in protein-coding gene predictions

We aimed to identify and correct structural errors in every predicted gene in the QX1410 reference gene models by leveraging short- and long-read RNA sequence (RNA-seq) alignments. We extracted and pooled RNA from mixed-stage, stage-specific, male-enriched, or starved cultures to maximize transcript representation across all stages of development and both sexes. We sequenced the QX1410 transcriptome using both Pacific Biosciences (PacBio) Single-Molecule Real-Time (SMRT) and Illumina platforms and refined the PacBio long reads into 95,177 high-quality transcripts using the IsoSeq pipeline [15]. We generated a set of long-read based gene models by identifying common intron chains and predicting open reading frames (ORFs) in assembled PacBio high-quality transcripts using StringTie and TransDecoder [16, 17]. Additionally, we generated a set of gene models from short-read RNA-seq alignments using BRAKER (Figure 1) [18]. A merger of these long- and short-read based gene models was previously published along with the chromosomal genome assembly of QX1410 [5]. The manual curation process relied on the identification of differences between computationally generated gene models and underlying RNA read alignments. We loaded short- and long-read alignments and gene models as individual evidence tracks into the Apollo genome annotation editor. We complemented the high-depth Illumina RNA-seq data with the structural information from full-length PacBio transcripts to provide high-confidence gene models for every locus with RNA coverage (Figure 2). When appropriate, we edited the structure of gene predictions to match the intron chain and gene termini best supported by RNA evidence. We manually curated 22,189 genes, amounting to a total of 32,278 transcripts (1.45 transcripts/gene). Comparisons of protein sequences between QX1410 gene models indicates we modified the coding sequences of 8,064 genes during the curation process. From these modified genes, 390 (4.8%) were classified as a gene split, 919 (11.4%) were classified as gene fusion, 4,751 (58.9%) were classified as intron chain error (missing or additional exon, or exon-intron junction modification), and 2,004 (24.9%) were not classified because of missing manual entries in our curatorial records. Additionally, we modeled putative alternative splice isoforms when new splice sites were identified in the RNA alignments. We identified variation in spliced reads mapped to existing gene models and added 10,089 putative isoforms in 5,398 genes. We also annotated 50,131 preliminary UTRs, including 5’ UTRs for 58.9% of genes and 3’ UTRs for 64.8% of genes (Figure 3). Examples of each type of curation, including workflows to correct different gene model error types, are provided in the curation protocol (Supplemental Information 1).

**Figure 1:**
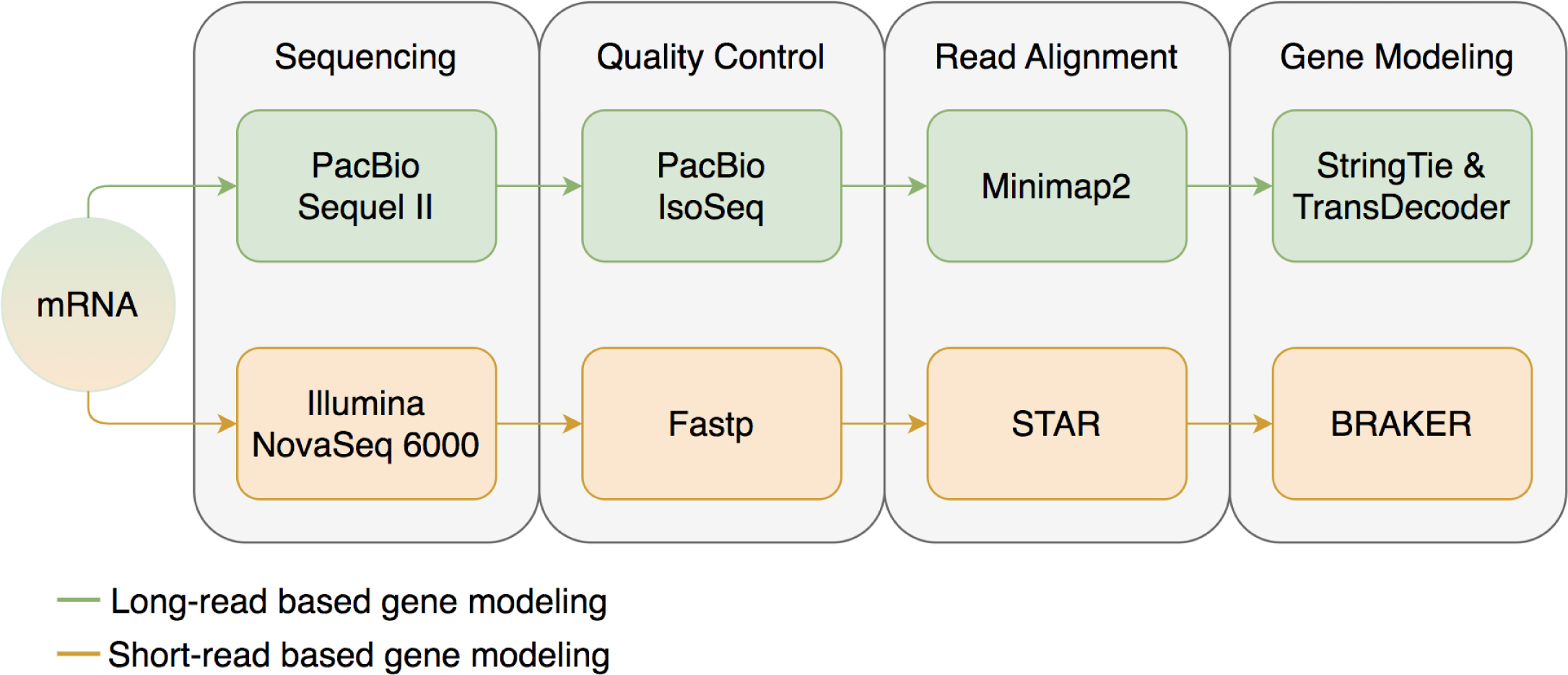
Data processing schematic. A flowchart describing the data processing steps required prior to the manual curation process. RNA is first sequenced using both PacBio and Illumina platforms. PacBio long reads are trimmed and refined using the IsoSeq pipeline, aligned to the reference genome using Minimap2, assembled into non-redundant transcripts using StringTie, and ORFs predicted using TransDecoder. Illumina short reads are trimmed using Fastp, aligned to the reference genome using STAR, and protein-coding genes are predicted using BRAKER.

**Figure 2:**
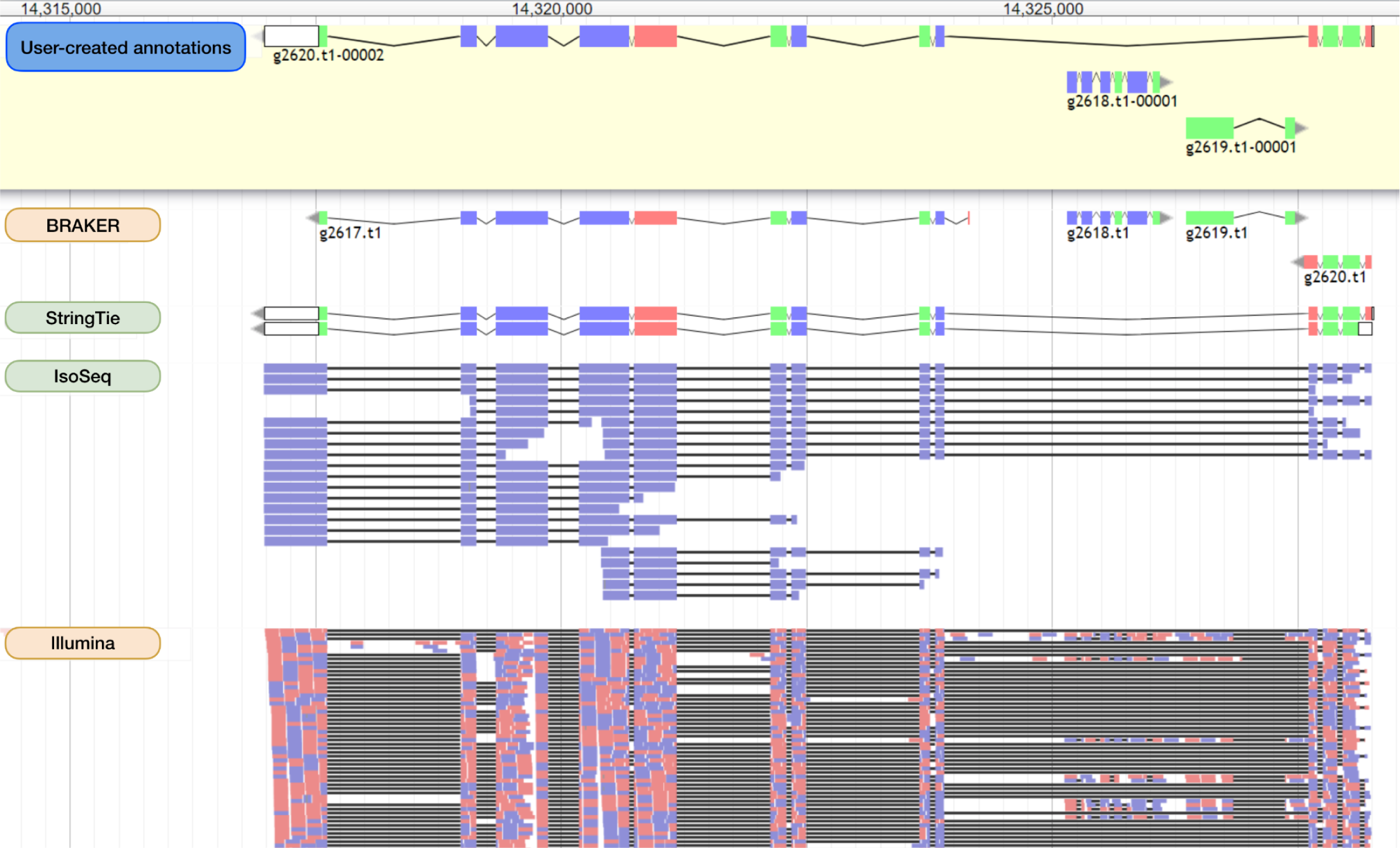
Identification of structural errors in protein-coding gene predictions. A screen capture of the Apollo genome annotation editor showing a set of manually curated genes (located on chromosome X from 14,314,000 to 14,325,000) and their underlying evidence. Four individual tracks are displayed from top to bottom: BRAKER gene models, StringTie gene models, PacBio Iso-Seq refined transcript alignments, and paired-end Illumina RNA-seq alignments. The final set of curated gene models is displayed in the top box shaded in yellow labeled ‘User-created Annotations’. Both Illumina and PacBio RNA data suggest that the two BRAKER genes at the ends of this region were incorrectly split. StringTie models resolve the incorrect split but lack the two internal genes on the opposite strand (g2618.t1 and g2619.t1), because they lack long-read RNA coverage. We kept the StringTie model that best matches the RNA evidence and added the two internal genes on the opposite strand predicted by BRAKER and supported by short-read RNA-seq. Curated and predicted gene models are colored by coding sequence phase. Illumina and IsoSeq alignments are colored by strand orientation.

**Figure 3:**
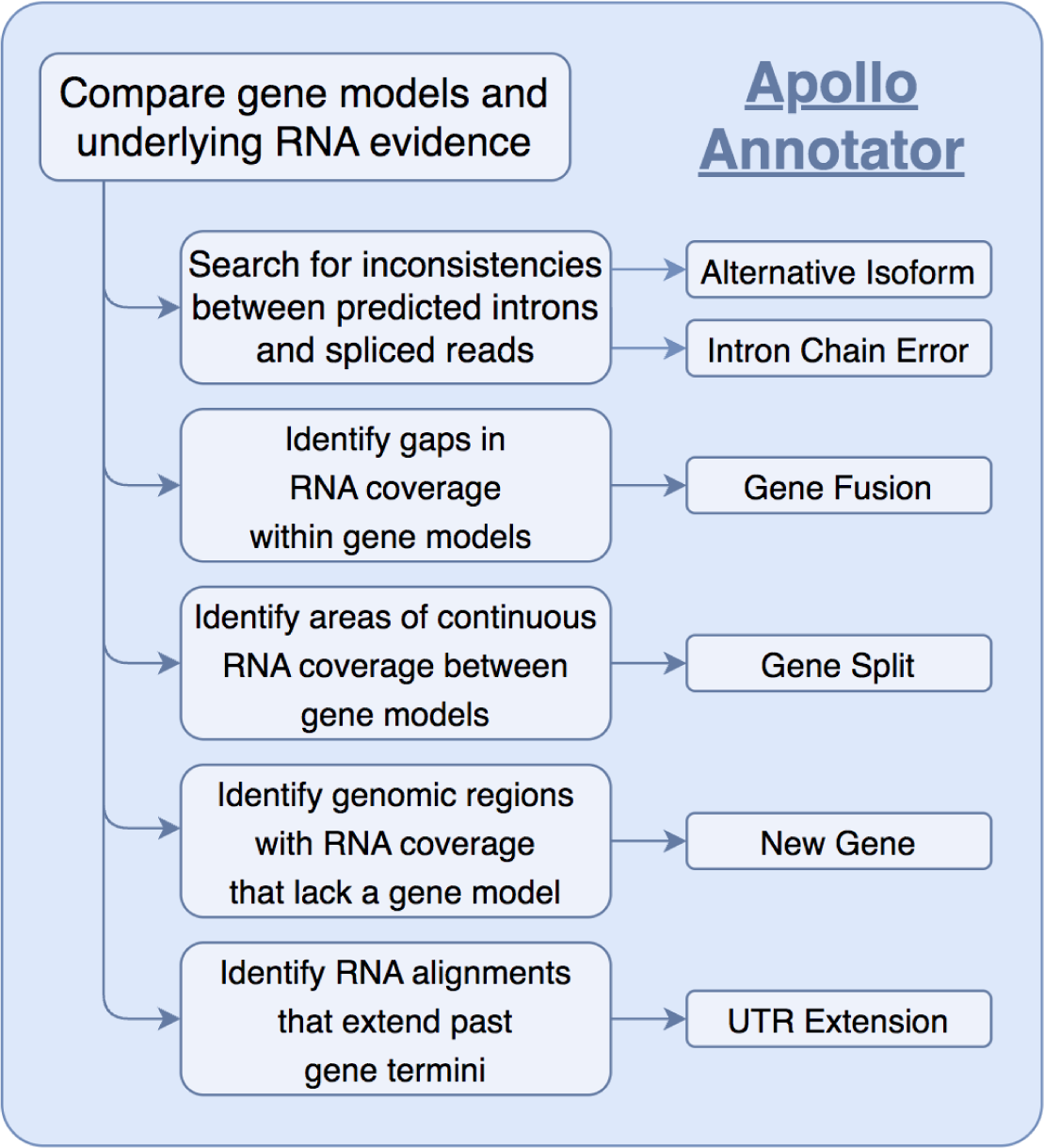
Schematic of manual curation process. Schematic describing the main logical steps to classify types of curation errors based on a comparison between RNA evidence and modeled genes from gene prediction tools. RNA evidence is sufficient to discern if a predicted gene model has major structural errors (*e.g*., gene fusion, gene split, intron chain error) or requires minor edits (*e.g*., additional isoform, UTR extension). Genomic regions with RNA coverage that lack a gene prediction were used to manually model genes *de novo*.

### Protein-length accuracy analysis reveals improvements after manual curation

To assess the quality of the manually curated *C. briggsae* gene models, we identified orthologous sequences between each *C. briggsae* gene set (QX1410 pre-curation, QX1410 post-curation, and AF16 WS280) and the reference *C. elegans* strain N2 using a reciprocal best BLAST hit approach. We identified the set of gene models in AF16 and QX1410 that share the same N2 ortholog, and compared the translated protein sequence length of each gene model against its respective N2 ortholog. Next, we estimated the ratio between a *C. briggsae* protein sequence and its best *C. elegans* reciprocal match, which we called ‘protein-length accuracy’. Finally, we counted the number of identical protein-length matches and matches that were within 5% of the protein length in N2 (5% off). Our manual curation efforts led to an overall improvement in protein-length accuracy (Figure 4). Specifically, we found a substantial increase in the number of identical and 5% off reciprocal BLAST matches (Table 1). Additionally, this improvement in protein-length accuracy was also accompanied by marginal improvements in BUSCO completeness. BUSCO completeness of the curated gene set was 99.7%, an improvement of 0.3% over the automated gene models, and 0.4% over AF16 (Table 1). The high BUSCO duplication values observed in both the curated and automated QX1410 gene models relative to AF16 gene models suggest an increase in alternative isoforms. Many isoforms were modeled from variation in spliced reads from short-read RNA-seq, and certain splice variant combinations might not exist as correct gene isoforms. Considering that manual curation was highly effective at reducing BUSCO duplication in the QX1410 genome (from 68.5% to 31.5%), the availability of full-length transcripts assembled from long-read RNA sequencing played a major role at identifying incorrect or redundant isoforms (Supplemental Information 2, Figure S1). Higher depth long-read RNA sequencing or other techniques specifically designed to maximize isoform discovery will be necessary to further identify and discard spurious isoforms that are still present in the current gene models.

**Figure 4:**
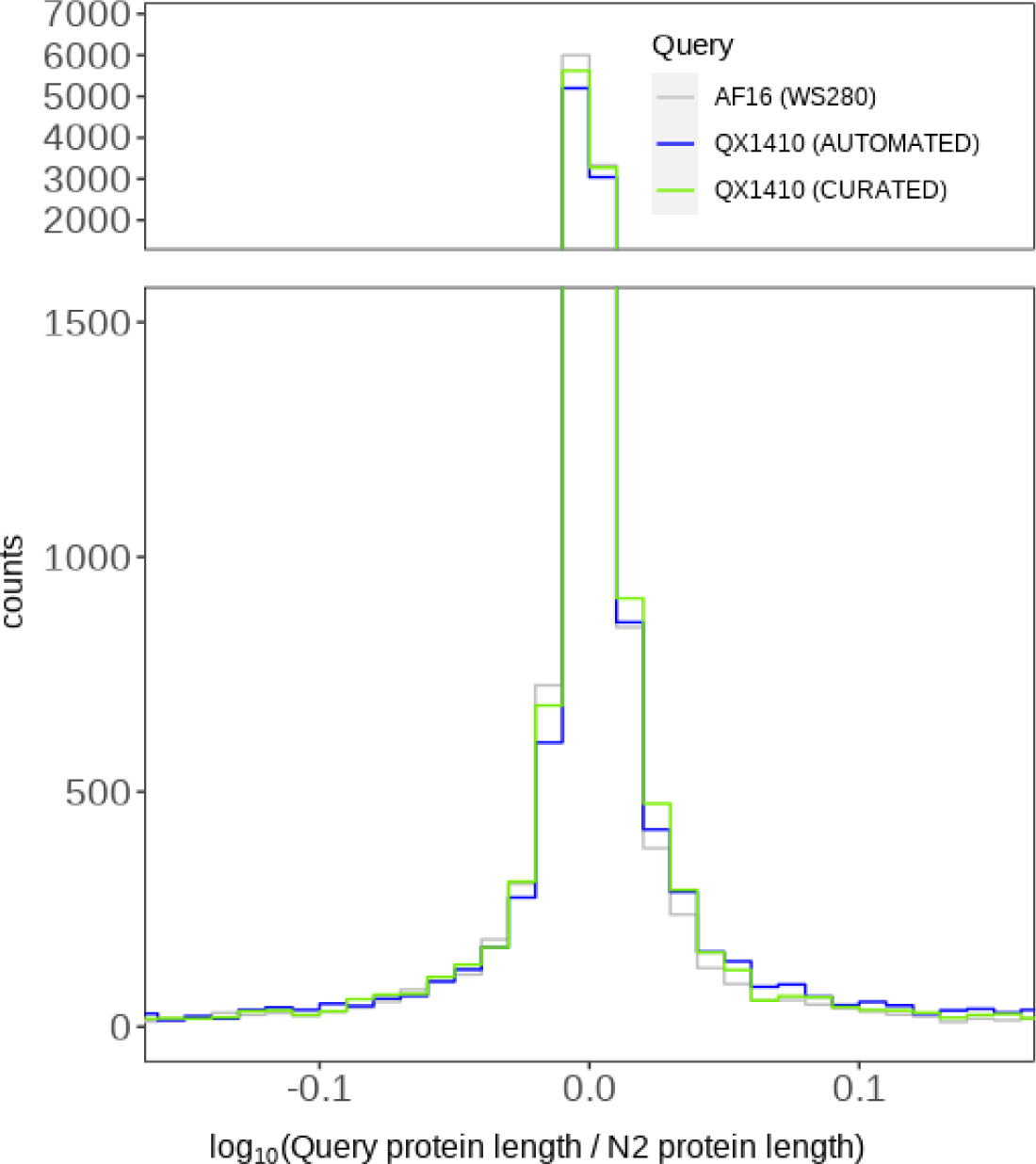
Protein-length accuracy of the *C. briggsae* gene models relative to reciprocal BLAST matches in *C. elegans* N2 proteins. The binned counts of log-transformed protein-length accuracy values for both sets of QX1410 gene models (automated and curated) and the AF16 gene models are shown. We calculated the protein-length accuracy of both automated and curated QX1410 gene models using the best reciprocal BLAST hit found in the N2 genome. We performed the same calculation for every gene in the AF16 (WS280) genome annotation. A value of zero represents a *C. briggsae* gene with a protein length that perfectly matches its best reciprocal hit in the N2 genome.

**Table 1.**
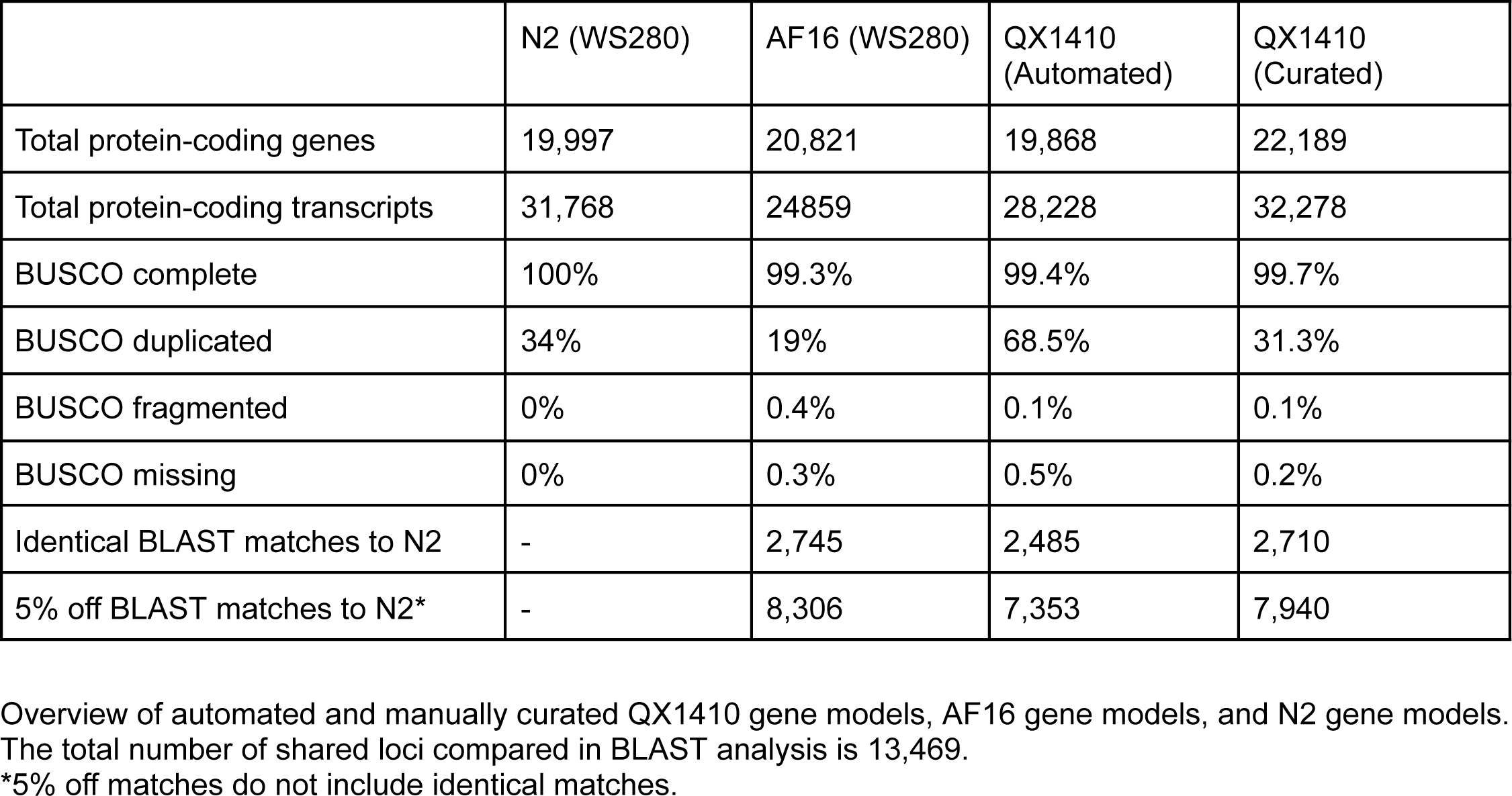
Annotation summary.

|  | N2 (WS280) | AF16 (WS280) | QX1410<br>(Automated) | QX1410<br>(Curated) |
| --- | --- | --- | --- | --- |
| Total protein-coding genes | 19,997 | 20,821 | 19,868 | 22,189 |
| Total protein-coding transcripts | 31,768 | 24859 | 28,228 | 32,278 |
| BUSCO complete | 100% | 99.3% | 99.4% | 99.7% |
| BUSCO duplicated | 34% | 19% | 68.5% | 31.3% |
| BUSCO fragmented | 0% | 0.4% | 0.1% | 0.1% |
| BUSCO missing | 0% | 0.3% | 0.5% | 0.2% |
| Identical BLAST matches to N2 | - | 2,745 | 2,485 | 2,710 |
| 5% off BLAST matches to N2* | - | 8,306 | 7,353 | 7,940 |

Overview of automated and manually curated QX1410 gene models, AF16 gene models, and N2 gene models. The total number of shared loci compared in BLAST analysis is 13,469.
\*5% off matches do not include identical matches.

### Protein-length analysis shows high concordance in protein-sequence length between the QX1410 and AF16 gene models

The vast majority of QX1410 genes are highly concordant in protein length with orthologous AF16 genes (Figure 5). Specifically, a total of 13,464 genes share the same N2 ortholog in the QX1410 and AF16 strains based on reciprocal BLAST analysis, and 12,690 (94%) are near-identical in protein length between the two *C. briggsae* strains. Considering the differences in genome assemblies and gene modeling methods between QX1410 and AF16, protein sequences that are distant from the N2 strain but highly concordant between both *C. briggsae* strains are likely to define true differences in protein sequence evolution between the two species. However, the curated QX1410 gene set lags behind the AF16 gene models in protein-length accuracy, with QX1410 showing slightly lower counts for identical and 5% off reciprocal BLAST matches to the N2 gene models (Figure 4, Table 1). To assess the predictive power and usefulness of standalone transcriptomic data in generating gene models for newly assembled genomes, our gene predictions and manual curations were performed agnostic of homology to *C. elegans* gene model data. The discrepancy in protein-length accuracy between the QX1410 and AF16 genomes is therefore partly explained by the use of *C. elegans* homology data in the curation process of the AF16 gene models. For example, we found that the selection of the translation initiation site when multiple in-frame start codons are present in the first exon can underlie small differences in protein length between the QX1410 and AF16 genomes (Supplemental Information 2, Figure S2). As a result, the use of defined *C. elegans* start codons during the development of AF16 gene models contributes to the difference in identical reciprocal BLAST matches when compared to QX1410 gene models with computationally predicted open-reading frames. To assess alternative explanations for the discrepancy in protein length accuracy between AF16 and QX1410, we revised 755 QX1410 genes that were shorter or longer in protein length than the respective N2 and AF16 orthologs (Figure 5). Based on RNA alignments alone, we made corrections to the coding sequences of 128 genes, leading to minor improvements in protein-length accuracy (Supplemental Information 2, Table S1). The 627 genes that remained unchanged were in agreement with underlying RNA alignments or had insufficient RNA coverage to manually curate. Considering that 78.6% of QX1410 genes that are at least 10% shorter or longer in protein length relative to N2 protein lengths are expressed below 20 transcripts per million (TPM), the differences in protein-length accuracy between the AF16 and QX1410 gene models can be attributed to insufficient RNA sequencing depth necessary to resolve the correct structure of lowly expressed genes (Supplemental Information 2, Figure S3).

**Figure 5:**
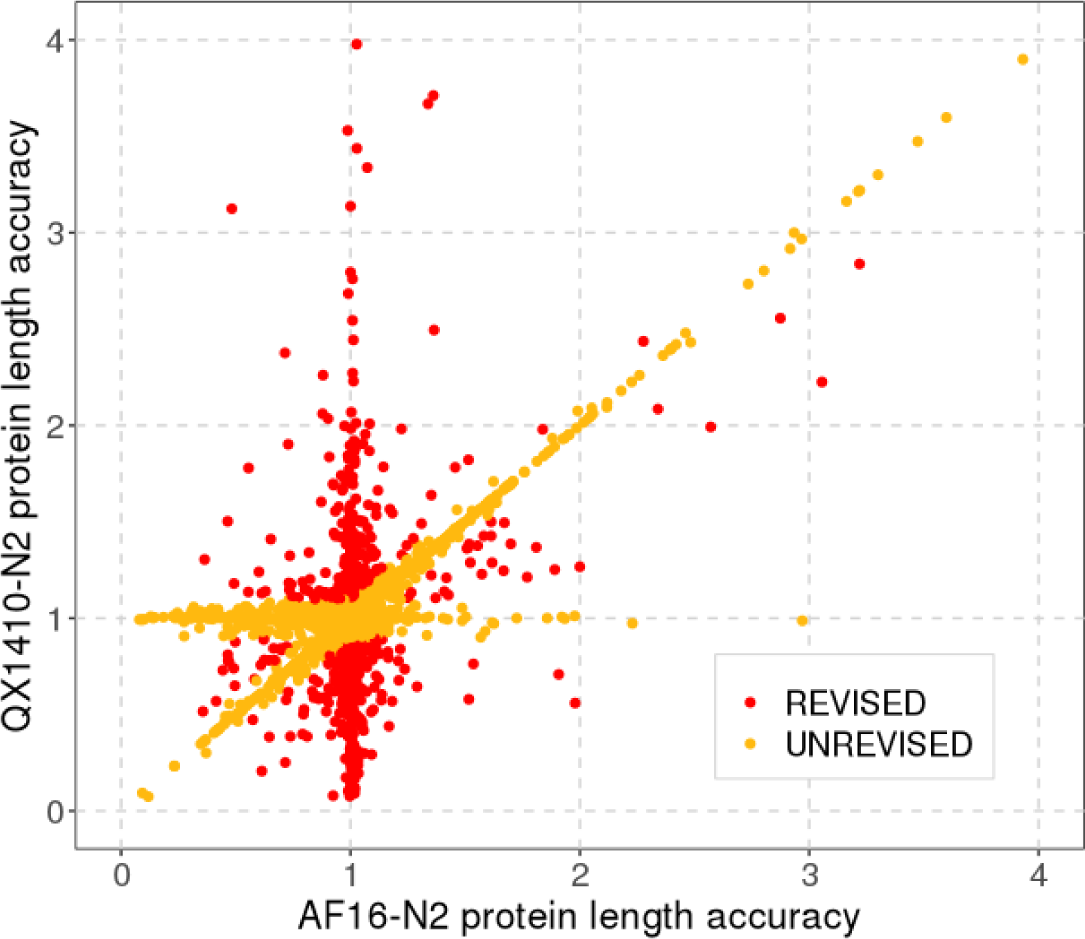
Protein-length differences from N2 between orthologous QX1410 and AF16 gene models. Each dot represents a gene in QX1410 that shares an N2 ortholog with an AF16 gene based on reciprocal BLAST analysis. We compared the protein length of each QX1410 and AF16 gene relative to its shared N2 ortholog and identified every QX1410 gene that is at least 10% shorter or longer in protein length (highlighted in red). We revised the RNA evidence used to model each gene that deviated in protein length relative to AF16 and made corrections when appropriate. Genes with protein-length accuracy values near to one (near-identical to the N2 gene) or genes that were near-identical in protein length between AF16 and QX1410 were not revised (highlighted in orange).

### Orthology analysis between *C. briggsae* and *C. elegans* shows incorrect gene duplications resolved in the QX1410 genome

In parallel to the protein-length accuracy analysis based on orthologs identified through sequence similarity estimates, we used OrthoFinder to study more complex orthologous relationships between *C. briggsae* and *C. elegans* genes [19]. This approach accounted for variable rates of sequence evolution between genes and facilitated the distinction of single-copy and multiple-copy orthologs between the two species. This distinction enabled the exclusion of multiple-copy orthologs, which prevented incorrect comparisons of protein length between genes of the same gene family. This approach also enabled the identification of orthologous relationships with lower identity that are missed using sequence similarity estimates used by BLAST search. We clustered 88,905 protein sequences translated from the gene models of the *C. elegans* N2 strain and both *C. briggsae* strains, QX1410 and AF16, into 17,790 putative orthologous groups. We identified 10,984 single-copy orthologs among all three strains, from which 86.2% were in agreement with reciprocal matches from the previous BLAST analysis, 0.3% were in disagreement, and 13.5% were not previously identified. Analysis of protein-length accuracy using only single-copy orthologs shows that the AF16 gene models still have higher counts of protein sequences with identical length to their N2 orthologs, but QX1410 gene models have higher counts for sequences that are within 5% of the protein length of their N2 orthologs (Supplemental Information 2, Figure S4 and Table S2). The discrepancy in number of identical BLAST matches is explained by the same factors identified in the sequence similarity analysis, including the use of *C. elegans* homology data or limitations in sequencing depth and transcript coverage. Additionally, the exclusion of multiple-copy orthologs and the inclusion of previously missed orthologs influenced the change in pattern for sequences that are within 5% of the protein length of their N2 orthologs. Both QX1410 and AF16 gene models appear to have small subsets of genes with higher protein-length accuracy, which is reflected in the differences in protein-length accuracy counts when using different ortholog selection criteria (Figure 5). Nonetheless, the orthology comparisons using either BLAST or OrthoFinder are largely in agreement and suggest that this manual curation of the QX1410 gene models using a single instance of transcriptome sampling provides gene models that are similar to the quality of the extensively curated AF16 gene models.

Interestingly, we identified a set of 317 single-copy orthologs between the QX1410 and N2 strains that are found as two or more copies in the AF16 strain. Manual inspection of orthologs that are found as two copies in AF16, but as single-copy in QX1410 and N2 (2:1:1 orthologs), suggests that additional gene copies have originated from misassembly of genomic contigs in the AF16 genome leading to spurious duplications. Approximately 30% of 2:1:1 orthologs are found in two separate scaffolds in the AF16 genome, often in both chromosomal and non-chromosomal (unplaced) scaffolds. The remaining 70% of 2:1:1 orthologs are found in the same chromosomal scaffold and provide clear evidence of spurious duplications and inversions that occurred during genome assembly. For example, we found two orthologs of the N2 *ubh-1* gene on chromosome II of the AF16 genome: *Cbr-ubh-1* and *CBG18955*. We performed a collinearity analysis of the QX1410 and AF16 genomes and revealed that both AF16 *ubh-1* orthologs and adjacent sequences align uniquely with a single segment of the QX1410 genome. Additionally, the *CBG18955* locus appears to be inverted relative to the *Cbr-ubh-1* locus (Figure 6A). The same duplication pattern has been observed in many other identified 2:1:1 orthologs, with or without inversion (Figure 6B-D). By using the alignment patterns observed in the manually inspected QX1410 and AF16 2:1:1 orthologs, we identified 5,316 spurious intra-chromosomal duplication events in the AF16 genome affecting a total of 1,896 genes (Supplemental Information 2, Table S3). Therefore, alignments between the AF16 and QX1410 genomes suggest that additional gene copies in the AF16 genome arise from artifactual duplications and/or inverted regions both within the chromosomal scaffolds and in unplaced scaffolds. Considering that the described genome assembly issues in the AF16 genome also affect multiple-copy orthologs, non-orthologous genes, and duplication events between chromosomes that were not included in this analysis, the QX1410 genome and gene models have not only resolved the duplication of numerous single-copy orthologs but also artifactual expansions of gene families present in the AF16 genome.

**Figure 6:**
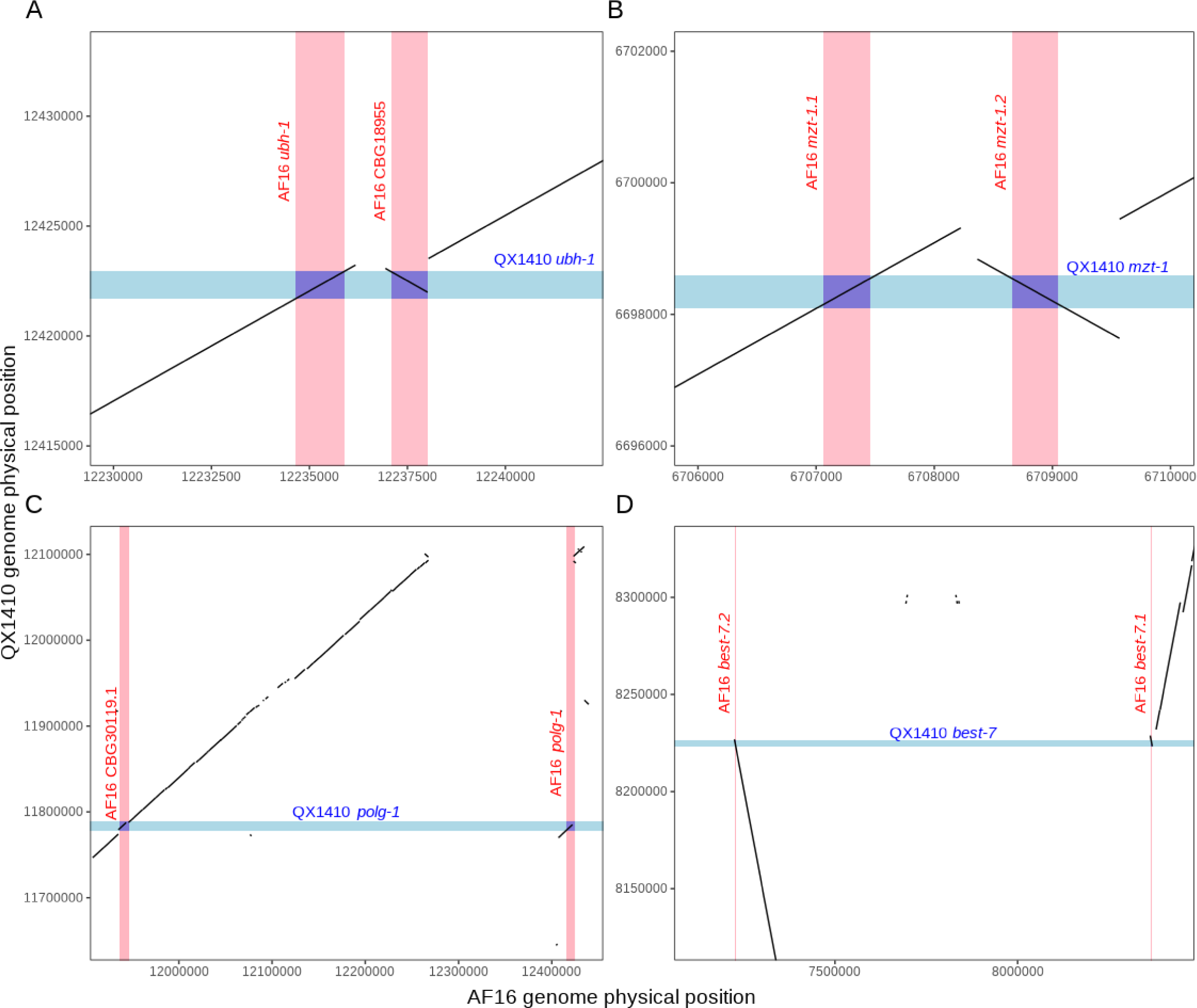
Examples of duplicated, inverted, and misplaced sequences in the AF16 genome assembly. Four plots showing the aligned QX1410 and AF16 genomes restricted to the physical positions of the *ubh-1* locus on chromosome V (A), *mzt-1* locus on chromosome I (B), *polg-1* locus on chromosome II (C), and *best-7* locus on chromosome IV (D). The gene boundaries are shaded in blue for the QX1410 genome, and shaded in red for the AF16 genome. The genomic sequences that contain each ortholog copy in the AF16 genome align uniquely to a single sequence in the QX1410 genome, demonstrating that the AF16 genome has an incorrect duplication. We selected two regions where the sequences are duplicated and in close proximity (A and B), and two regions where the sequences are duplicated and distant (C and D).

## Discussion

Although the development of high-quality genomic resources has traditionally been an arduous and time-consuming process, recent advancements in long-read sequencing technologies and associated computational tools have enabled the development of fast and effective methods to generate highly contiguous and complete genome assemblies for newly sequenced organisms [18, 20]. Using these techniques, improved reference genomes have been generated for numerous nematode species [3, 4, 21, 22]. In a previous study, we presented a new reference genome for the *Caenorhabditis briggsae* strain QX1410 [5]. This genome resolved thousands of gaps and numerous missasemblies that were present in the existing reference genome for the laboratory strain AF16. Additionally, we modeled protein-coding genes using a combination of short- and long-read transcriptome sequencing with modern gene prediction tools [5]. However, protein-coding gene prediction accuracy remains a challenge. When compared to the *C. elegans* reference gene models, errors such as the retention of non-coding sequences, incorrect intron-exon junctions, and gene fusions or splits are prevalent across thousands of loci in the new QX1410 reference genome. Although new computational tools could resolve these prediction errors in the near future, community-based manual curation has proven an effective method to improve the quality of protein-coding gene models in nematodes [21, 23, 24]. Here, we manually curated every *C. briggsae* QX1410 gene, attesting to the feasibility of large-scale community curation projects. We exploited the high level of protein length conservation between *C. briggsae* and *C. elegans* to quantify the changes in protein-length accuracy of manually curated QX1410 gene models relative to orthologous *C. elegans* N2 gene models. We performed the same comparison in the existing AF16 reference gene models. This analysis revealed substantial improvements in protein-length accuracy after manual curation, accompanied by a slight improvement in BUSCO completeness. Additionally, according to the protein-length metric, fewer QX1410 gene models have identical or near-identical protein-length matches to N2 relative to the AF16 gene models. Manual revisions of loci that underperform in protein-length accuracy in QX1410 revealed that most genes with shorter or longer protein sequences than their N2 orthologs have insufficient RNA coverage to manually curate. The RNA extraction protocols that we used maximize transcript representation across all stages of development, but it is possible that transcripts from certain stages are underrepresented in the pool of RNA extracted from the different developmental stages and sexes. Additional transcriptome sequencing of stage-specific samples will be required to further maximize transcript discovery and improve gene models that currently lack sufficient RNA coverage. With single-cell RNA-seq becoming more affordable, future efforts to further improve the QX1410 protein-coding gene models will benefit from RNA sequencing of specific tissues that may be underrepresented in the transcriptome sequencing effort. Despite the limitations in sequencing coverage, this single instance of manual curation led to QX1410 gene models that are comparable to AF16 gene models in accuracy and completeness. Moreover, we identified over 1,800 loci affected by spurious duplications stemming from assembly errors in the AF16 genome that are now resolved in the QX1410 genome. The small differences in protein-length accuracy among the QX1410 and AF16 gene models are overshadowed by the numerous errors that persist in the AF16 genome. As a result, the high level of sequence contiguity in the QX1410 genome provides a reliable platform to continue to improve and expand the *C. briggsae* genome resources. The availability of high-quality *C. briggsae* reference gene models in the correct genomic context will also help establish complete and accurate gene family relationships within the species, improve measurements of genetic relatedness among *C. briggsae* wild isolates and phylogeographic groups, and enable candidate gene discovery in genome-wide association studies.

A major motivation to improve the quality of *C. briggsae* genome resources is to enable new avenues to investigate *Caenorhabditis* genome evolution. Specifically, recent studies in *C. elegans* wild isolates have led to the discovery of punctuated genomic regions with abnormally high genetic diversity relative to expectations of diversity in self-fertilizing organisms. These hyper-divergent regions are thought to retain ancestral alleles maintained by balancing selection during the evolutionary history of *C. elegans*. These regions appear to be implicated in local adaptation because they harbor unique sets of environmental response genes among different *C. elegans* wild isolates [25]. The characterization of the gene content of hyper-divergent regions in *C. elegans* was made possible because of the high-quality genome resources available for this species. Interestingly, evidence points to the presence of hyper-divergent regions in a small subset of *C. briggsae* wild strains [25]. The availability of complete and accurate genome resources for *C. briggsae* will be essential to characterize and test hypotheses surrounding the origin and function of these regions.

## Conclusion

In this study, we presented a detailed workflow to manually curate gene models using short- and long-read RNA sequencing data. Our manual curation efforts led to improved gene models for the new chromosome-level reference genome for the *C. briggsae* strain QX1410. According to our protein-length metrics and BUSCO completeness scores, the first-pass manual curation of the QX1410 gene models are similar in quality to the extensively curated gene models of the laboratory strain AF16. Considering the pervasive errors in the AF16 genome assembly, the use of QX1410 genome resources provides a reliable and improved platform to perform genomic studies in this species and make comparisons against other species in the *Caenorhabditis* genus.

## Methods

### Nematode culture

Animals were reared at 20°C on standard nematode growth medium (NGMA) plates, and *Escherichia coli*

OP50 was used as a food source.

### RNA library preparation

We extracted RNA from a mixed-stage and mixed-sex population composed of samples from all larval stages, adults, dauers, and males. Plates containing adults and larvae from every stage were prepared by chunking every two days for several generations. Plates containing dauer and arrested L1 and L2 larvae were prepared by allowing the plate to starve. Male-enriched plates were prepared by setting up crosses between male and hermaphrodite animals and expanding the population for up to three generations. We collected a sample from one 10 cm plate for each population (mixed-stage, starved, male-enriched) into 100ul S Basal. Samples were flash frozen in liquid nitrogen and stored at −80°C. RNA was extracted from each sample using 1 mL TRIzol reagent (Invitrogen, catalog no. 15596026) with addition of 100 uL of acid-washed sand to aid sample homogenization, and resuspended in nuclease-free water. We used a Nanodrop spectrophotometer (ThermoFisher) to quantify the purity of the RNA samples, and a Bioanalyzer and a Qubit (ThermoFisher) were used to determine RNA concentration. The Qubit RNA HS Assay kit (ThermoFisher, catalog no. Q32852) was used. We pooled 1.5 mg RNA from each sample, and purified and concentrated the pooled RNA using the RNeasy MinElute Cleanup kit (Qiagen, catalog no. 74024). We eluted the purified RNA into nuclease-free water and repeated the quality control steps (purity and concentration determination) as described previously.

### Short-read RNA sequencing

The Illumina RNA-seq library was prepared in a 96-well plate. We purified and enriched mRNA from 1 ug of total RNA using the NEBNext Poly(A) Magnetic Isolation Module (New England Biolabs, catalog no. E7490L). RNA fragmentation, first and second strand cDNA synthesis, and end-repair processing was performed using the NEBNext Ultra II RNA Library Prep with Sample Purification Beads (New England Biolabs, catalog no. E7775L). We ligated adapters in the cDNA library using adapters and unique dual indexes from the NEBNext Multiplex Oligos for Illumina (New England Biolabs, catalog no. E6440L). All procedures were performed following the manufacturer protocols. We used Qubit dsDNA BR Assay Kit (Invitrogen, catalog no. Q32853) to determine the concentration of the RNA library. The library was then pooled and qualified using the 2100 Bioanalyzer (Agilent) at Novogene, CA, USA. We sequenced the pooled library with the Illumina NovaSeq 6000 platform (150-bp paired-end reads).

### Long-read RNA sequencing

The PacBio Iso-Seq full-length sequencing library was prepared using 300 ng of total RNA using NEBNext Single Cell/Low Input cDNA Synthesis and Amplification Module (NEB, catalog no. E6421) and SMRTbell Express Template Prep Kit 2.0 (Pacific Biosciences, catalog no. 100-938-900). This library was prepared in the Duke Center for Genomic and Computational Biology’s Sequencing Technologies Core facility, and was sequenced using three SMRT cells.

### Repeat Masking

Prior to gene prediction, we masked repetitive sequences in the QX1410 genome to avoid spurious predictions. We generated a custom repeat library using a previously described approach [3, 26]. In summary, we used RepeatModeler from RepeatMasker v2.0.1 [27] for *de novo* repetitive sequence identification. Additionally, we identified transposable elements using TransposonPSI [28], and long terminal repeat (LTR) retrotransposons using LTRharvest from GenomeTools v1.6.1 [29, 30]. Identified LTR retrotransposons were annotated using LTRdigest from GenomeTools with HMM domain profiles from Gypsy Database 2.0 [31] and select Pfam domains [32] (listed in tables SB1 and SB2 of [33]). We removed repeat candidate sequences without conserved protein domains using *gt-select* from GenomeTools. Lastly, we retrieved Rhabditida-specific repeats Dfam [34] and *C. elegans* ancestral repeats from RepBase [35]. We merged newly generated and retrieved repeat libraries into a single redundant library. We clustered and classified repeats in the redundant library using VSEARCH v2.14.2 [36] and RepeatClassifier from RepeatMasker, respectively. Unclassified repeats with significant BLAST hits to the *C. elegans* proteome (WS279) were removed. We soft-masked the QX1410 genome assembly using RepeatMasker.

### Automated protein-coding gene prediction

The QX1410 genome was retrieved from NCBI under the accession PRJNA784955 [5]. We aligned short-read RNA sequences to the soft-masked QX1410 genome using STAR v2.7.3a [37] with a maximum intron size of 10 kilobases, and generated protein-coding gene predictions using BRAKER v2.1.6 [18]. Additionally, we generated high-quality transcripts from PacBio long RNA reads using isoseq3 v3.4.0 [15] and aligned them to the QX1410 genome using minimap2 [38]. We performed transcriptome assembly using StringTie v2.1.2 [16] from PacBio high-quality transcript alignments. We predicted coding sequences in the assembled transcripts using TransDecoder [39]. We assessed the biological completeness of generated gene models using BUSCO v5.0 [40]

### Manual curation of gene prediction errors

Short- and long-read sequence alignments and gene models were uploaded as individual tracks to the Apollo platform v2.6.4 [14] for manual curation. We repaired three classes of gene prediction errors: gene splits, gene fusions, and intron chain errors (additional, missing, or modified exons). We identified potential gene splits by inspecting RNA alignments in intergenic regions. When two adjacent gene models have uninterrupted long- and short-read RNA coverage across their intergenic region, we merged both models and re-predicted the coding sequence. If the gene merger led to a premature stop codon, we reverted the merger. We identified gene fusions by searching for gaps in RNA coverage within a single gene accompanied by a predicted intron that bypasses a stop codon with RNA coverage. Additional, missing, or incorrect exons of a gene were repaired using the consensus exon-intron structure between long-read transcripts and collapsed short-read alignments. If long-read RNA transcripts were not available to confirm a potential split, fusion, incorrect exon identified solely by short-read RNA reads, or were in disagreement with short-read RNA alignments, we kept both alternative models (original and manually curated).

### Manual curation of coding sequences and untranslated regions

We used the built-in open-reading frame (ORF) prediction tool from Apollo to select the optimal coding sequence for each transcript. This tool weights longer ORFs over coding sequence continuity across all exons. We manually set start and stop codons that led into a shorter ORF that spanned a higher number of exons compared to the predicted ORF with longer coding sequence. We exploited full-length, long-read transcripts to extend untranslated regions of every gene. When long-read transcripts were unavailable, we used the longest set of short read alignments that overlapped with the coding sequence of the gene. We matched the UTR boundaries of gene isoforms with identical terminal exons. We did not annotate UTRs of genes in close proximity and in the same strand with continuous short-read coverage throughout their intergenic region.

### Manual curation of additional isoforms and new genes

In many cases, multiple isoforms were predicted by BRAKER and StringTie. We removed any isoforms that were unsupported by both short- and long-read alignments. We added any potential isoforms that were present as structurally unique long-read transcripts. Isoforms modeled from long-read transcripts that had substantially shorter coding sequences (one or more non-coding exons) were kept. Differences in structure observed between the predicted gene models and a subset of the short-read alignments were also included as additional isoforms. In cases where multiple structural differences were observed in short-read alignments, we modeled all possible structural permutations that had a continuous ORF from the first to the last exon. When a multi-exon structure was observed in the short-read alignments and no genes were predicted at that locus, we modeled a gene *de novo*. We discarded any *de novo* gene models that did not have an ORF.

### Protein sequence length analysis

We generated BLAST v5 libraries for each set of *C. briggsae* (QX1410 and AF16) protein sequences and for *C. elegans* (N2). We performed forward and backward BLASTp v2.12 searches between N2 and each *C. briggsae* strain [41]. We selected the best hit (ranked using expectation value and bitscore) for every protein sequence in the forward search that had a reciprocal best hit in the backward search. We calculated the ratio of protein sequence length between each *C. briggsae* protein sequence and its reciprocal *C. elegans* protein sequence. We kept only one reciprocal hit per gene. We counted the number of protein sequences with identical protein length matches and matches that were within 5% of the length of its respective reciprocal hit.

### Orthology analysis

We extracted protein sequences from gene model files in GFF format using Gffread v0.12.1 [42]. The GFF files were filtered to only keep the longest isoform per gene using *agat_sp_keep_longest_isoform.pl* from AGAT v0.8.1 [43]. We clustered protein sequences extracted from *C. briggsae* QX1410, *C. briggsae* AF16, and *C. elegans* N2 gene models into orthologous groups using OrthoFinder v2.1.4. We classified single-copy and multiple copy orthologs across all three strains in R. We performed the same protein sequence length analysis described previously on single-copy orthologs identified across all three strains.

### Collinearity analysis

The AF16 (WS280) and QX1410 genomes were aligned using NUCleotide MUMmer (NUCmer) v3.1, allowing a maximum gap of 500 bp [44]. Duplicated sequences of the AF16 genomes were identified using R by selecting alignments that had a single set of coordinates in the QX1410 genome, but distinct sets of coordinates in the AF16 genome.

## Data Availability

Raw sequencing data, genome assembly, and gene model files have been archived under the study accession PRJNA784955. Code used to produce analyses and figures can be found at: https://github.com/AndersenLab/briggsae_gene_models_MS

## Acknowledgements

We would like to thank members of the Andersen laboratory that reviewed this manuscript and provided comments and suggestions. We would also like to thank Gaotian Zhang, Rojin Citrakar, and Ryan L. Baugh for the sequencing efforts that led to the initial work to develop the new *C. briggsae* genome. Lastly, we are deeply thankful for the *Caenorhabditis* research platform WormBase, because their contributions continue to support the nematode research community.

## Funding

This work was funded by grants to E.C.A. from NIH (R01 ES029930) and the Human Frontiers Science Program (RGP0001/2019). E.C.A was also funded by a subcontract (R21 OD30067) from Helen Chamberlin at Ohio State University. N.D.M. was supported in part by the National Institutes of Health Training Grant (T32 GM008449) through Northwestern University’s Biotechnology Training Program

## Author Information

### Authors and Affiliations

Department of Molecular Biosciences, Northwestern University, Evanston, IL 60208, USA Nicolas D. Moya, Isabella R. Miller, Chloe E. Sokol, Joseph L. Galindo, Alexandra D. Bardas, Edward S. H. Koh, Justine Rozenich, Cassia Yeo, Maryanne Xu, and Erik C. Andersen1,‡

Interdisciplinary Biological Sciences Program, Northwestern University, Evanston, IL 60208, USA Nicolas D. Moya

Tree of Life, Wellcome Sanger Institute, Cambridge, UK Lewis Stevens

### Contributions

Conceptualization, E.C.A.; Investigation, E.C.A. and N.D.M., and L.S.; Data curation: N.D.M., I.R.M., C.E.S., J.L.G., A.D.B., E.S.H.K., J.R., C.Y., and M.X.; Visualization, E.C.A. and N.D.M.; Writing original draft, E.C.A. and N.D.M.; Writing - review & editing, E.C.A. and N.D.M.; Project Administration, E.C.A.; Supervision, E.C.A.; Funding acquisition, E.C.A. The author(s) read and approved the final manuscript.

### Corresponding Author

Correspondence to Erik C. Andersen

## Notes

https://github.com/AndersenLab/briggsae_gene_models_MS

### Competing Interest Statement

The authors have declared no competing interest.

