## Supplemental Information 1 for "Novel and improved *Caenorhabditis briggsae* gene models generated by community curation"

***Caenorhabditis briggsae* gene annotations improved by community curation using short- and long-read RNA sequencing**

**Supplemental Information 1**

### Curation Protocol

#### Version 1.3

**Table of contents:**


1. Introduction
   1.1. Genomes and genes
   1.2. Short-read sequencing
      1.2.1. Illumina RNA-seq
      1.2.2. BRAKER: Short-read based gene models
   1.3. Long-read sequencing
      1.3.1. Single Molecule, Real-Time (SMRT) RNA sequencing
      1.3.2. StringTie: long-read transcriptome assembly
   1.4. Curation project aims

2. Apollo
   2.1. Navigating the genome browser
   2.2. Tracks
      2.2.1. BRAKER models
      2.2.2. StringTie models
      2.2.3. IsoSeq transcript alignments
      2.2.4. Illumina short-read alignments
      2.2.5. Track Visualization
   2.3. User-created annotations
      2.3.1. Adding annotations
      2.3.2. Removing annotations
      2.3.3. Renaming annotations
   2.4. Gene curation
      2.4.1. Only BRAKER
      2.4.2. BRAKER and limited Illumina reads
      2.4.3. UTR Extension
      2.4.4. Gene split
      2.4.5. Gene fusion
      2.4.6. Multiple Isoforms
         2.4.6.1. Simple multiple isoforms case
         2.4.6.2. Complex multiple isoforms case
      2.4.7. Terminal Exon Repair
         2.4.7.1. Missing Exon
         2.4.7.2. Additional Exon
      2.4.8. Incomplete coding sequence (CDS)
      2.4.9. New gene
   2.5. Curation metadata
   2.6. New cases

#### 1. Introduction

##### 1.1. Genomes and genes

An organism's genetic information is stored in double-stranded DNA molecules, packaged in compact units called chromosomes. DNA molecules are composed of a long, alternating sequence of four different nucleotides: adenine (A), thymine (T), cytosine (C), and guanine (G). A genome is the complete set of genetic information in an organism.

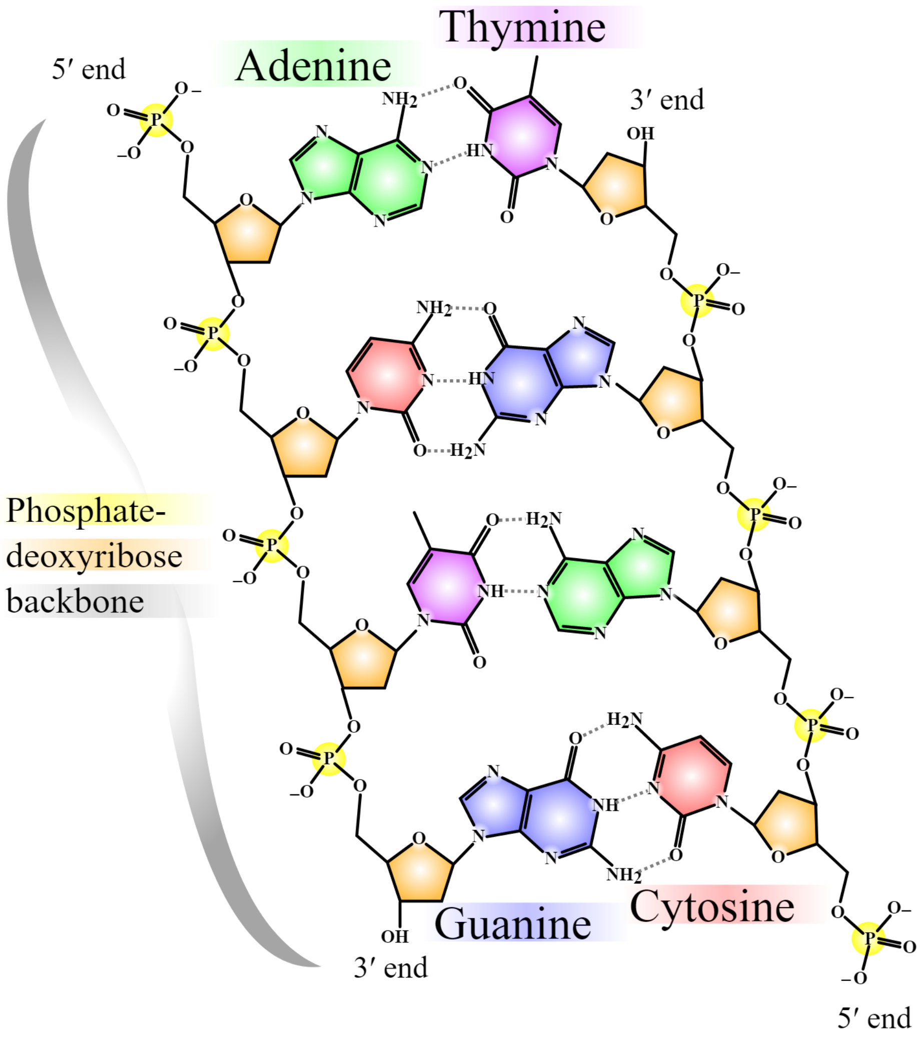


The central dogma of molecular biology describes the flow of genetic information from a DNA sequence to a functional protein. The process by which single-stranded RNA is synthesized from double-stranded DNA is called transcription. The process by which amino acid chains are translated from an RNA template is called translation.

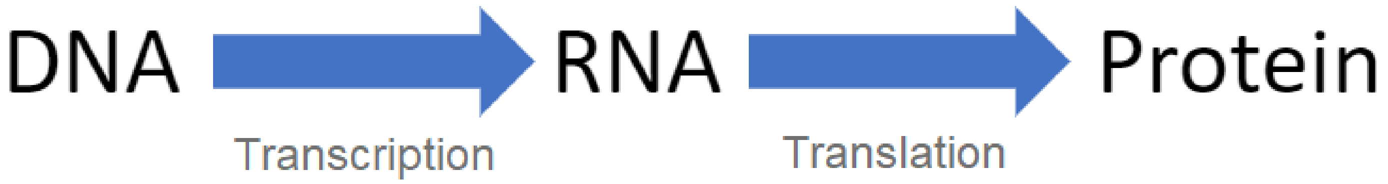


Within a genome, genes are discrete units. A gene is a specific sequence of DNA that encodes a functional product (RNA or protein). Genes are present in both DNA strands (shown below). Genes are composed of three distinct elements: untranslated regions (UTRs), exons, and introns. UTRs are non-coding sequences found at the 5’ and 3’ ends of a gene. As the name implies, these regions are not translated into amino acids but they are transcribed into RNA. Exons are the sequences of the gene that code for amino acids. Exons are interrupted by non-coding sequences called introns.

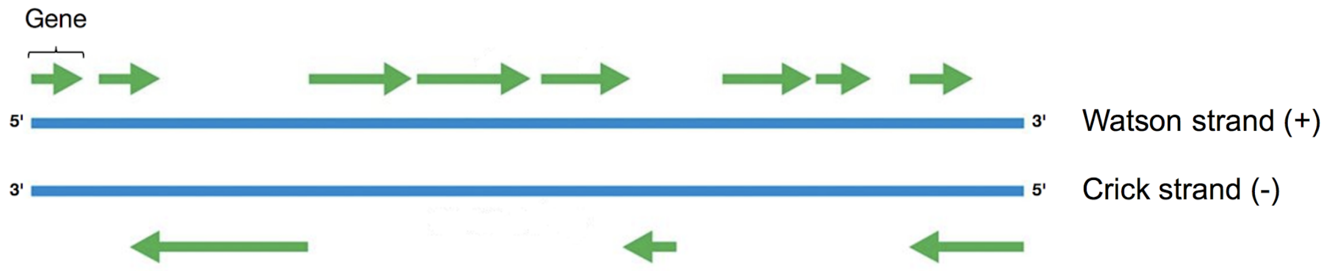


Protein-coding genes are transcribed into messenger RNA (mRNA). First, the transcription machinery binds to a promoter region upstream of the gene. This machinery ‘reads’ through the DNA sequence downstream of the promoter. Once the transcription machinery encounters a transcription start site, it will synthesize a mRNA molecule (transcript) using the DNA sequence as a template. Transcription continues until a transcription stop site is encountered. The nascent mRNA molecule that is produced during transcription is called precursor mRNA (pre-mRNA). A 5’ cap and a 3’ polyA tail are added to the pre-mRNA to prevent its degradation. Introns are then excised, and exons are joined together (splicing). UTRs are retained after pre-mRNA processing. Once introns are removed, the now mature mRNA is ready to be translated. Some mRNAs can be alternatively spliced, where multiple mature mRNAs can be formed by changes in the succession of exons joined during splicing. Different mature mRNA transcripts that are generated from a single pre-mRNA are called isoforms. Isoforms can also originate from alternative transcription start and stop sites within a single gene.

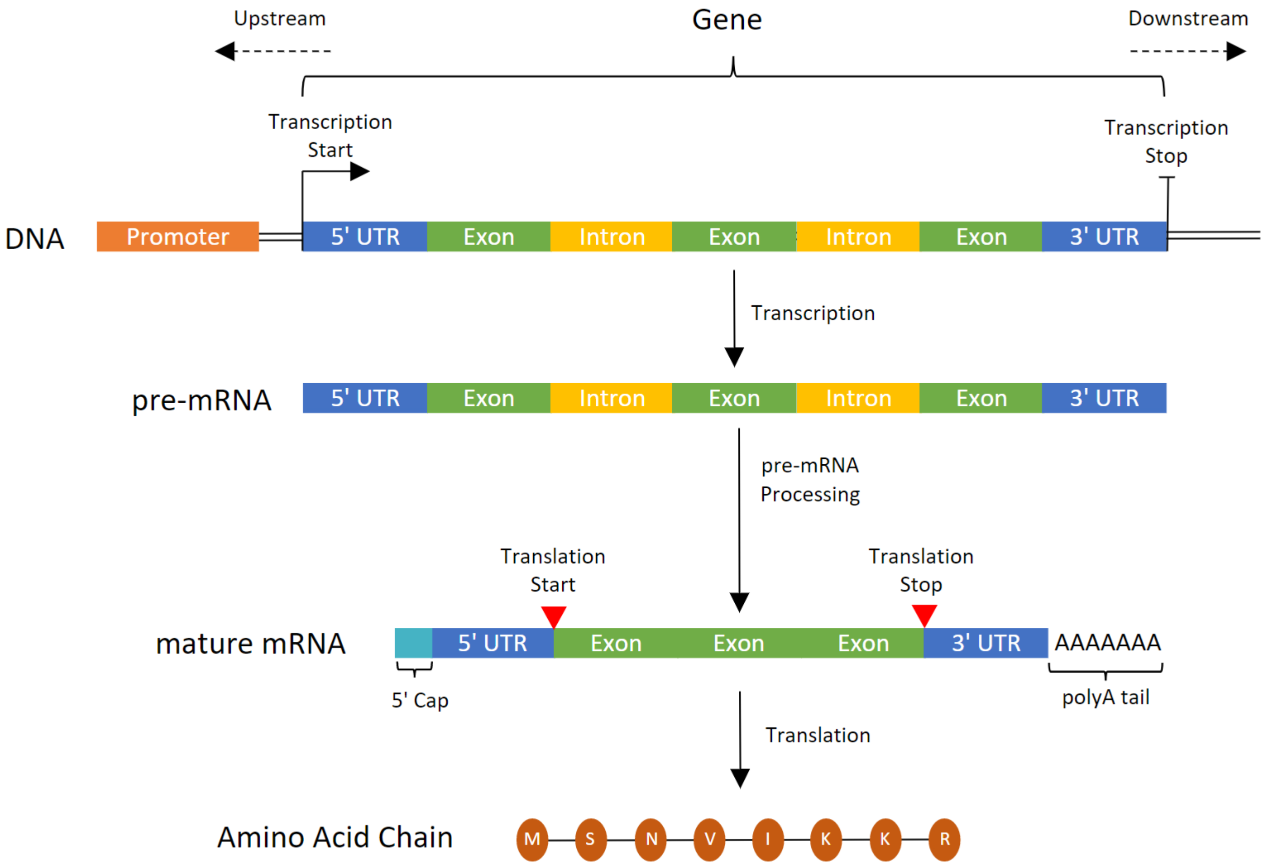


After splicing, the mature mRNA can be translated into an amino acid sequence. In translation, the instructions to build an amino acid chain from an mRNA template are in the form of codons. Codons are nucleotide triplets that signal for a specific amino acid. Translation is carried out by ribosomes. A ribosome (and associated factors) binds to the ribosomal binding site (RBS) within the 5’ UTR of the mature mRNA. The ribosome will ‘read’ through the mRNA transcript and will begin translation after it encounters a start codon (AUG). Translation will continue until a stop codon (UAA, UAG, UGA) is encountered. Refer to this translation table to see all possible codon combinations and their encoded amino acid: Translation Table

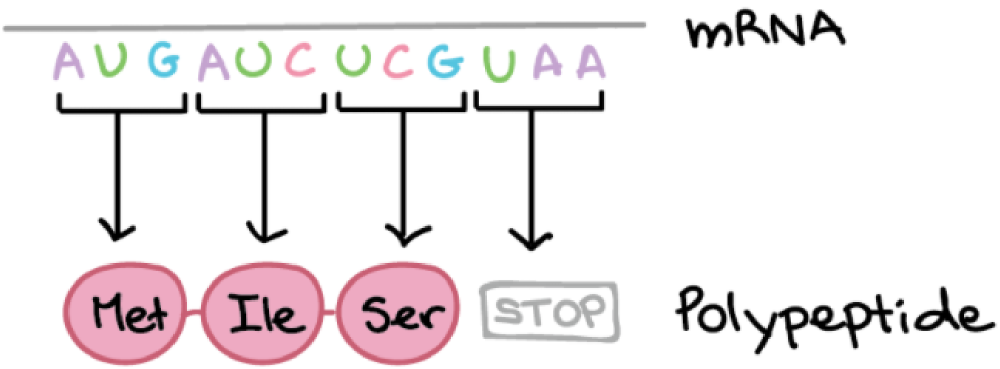

##### 1.2. Short-read sequencing

The field of genomics focuses on the large-scale characterization of genomes and genes. In the past few decades, advances in short-read sequencing technologies have allowed researchers to generate complete genomes for many organisms. Additionally, the development of RNA sequencing (RNA-seq) enabled researchers to predict the structure and location of genes in a genome. These gene predictions are referred to as gene models. In this section, we will review the molecular basis of RNA-seq and its application in generating gene models.

###### 1.2.1. Illumina RNA-Seq

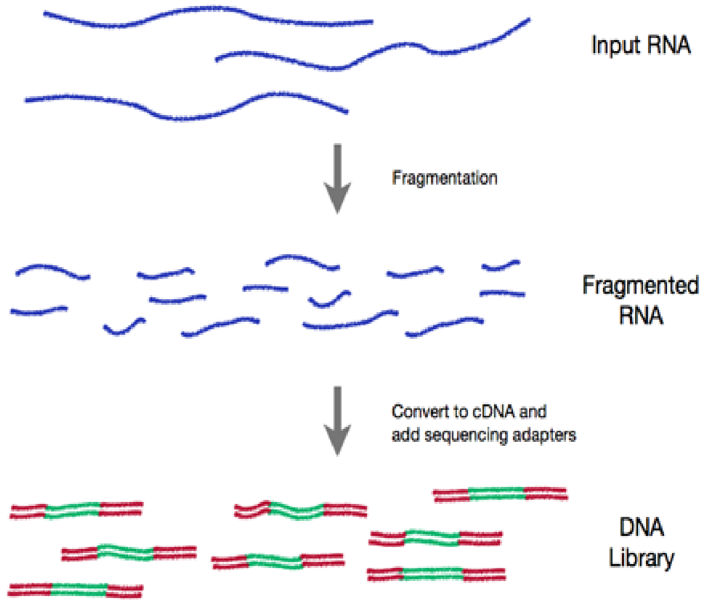


The first step in RNA-seq involves breaking mRNA into small fragments to create a library of complementary DNA (cDNA) fragments. cDNA synthesis stems from the process of reverse transcription, where double-stranded DNA is synthesized from a single-stranded RNA template. cDNA fragments are more stable than RNA and can be readily amplified and sequenced. The ends of the cDNA fragments are then ligated to specific oligonucleotide adapters that permit sequencing. The adapter-ligated cDNA library is amplified and sequenced using an Illumina sequencer machine. The sequencer determines the nucleotide sequence of each cDNA fragment. The sequence is called a 'read'. Billions of reads can be generated in a single RNA-seq run. The length of these reads ranges from 50 to 150 base pairs (bp), depending on the cDNA library preparation approach.

RNA reads can be aligned back to the genome to identify which genomic regions are actively transcribed. In the example shown on the right, 150 base-pair reads (grey bars) are aligned to a 1.3 kilobase (kb) region of a genome. As mentioned previously, RNA-seq reads are obtained from fragmented mRNA. Mature mRNA does not possess introns. As a result, reads that cover the intersection between two exons possess large gaps in their alignment (black lines). Additionally, short RNA reads are error prone, subject to modifications such as insertions (purple marks) and deletions (blue marks). The distribution of read alignments in this region alludes to the structure of a gene (exons interrupted by introns). We will next learn how these short-read alignments are used to generate gene models.

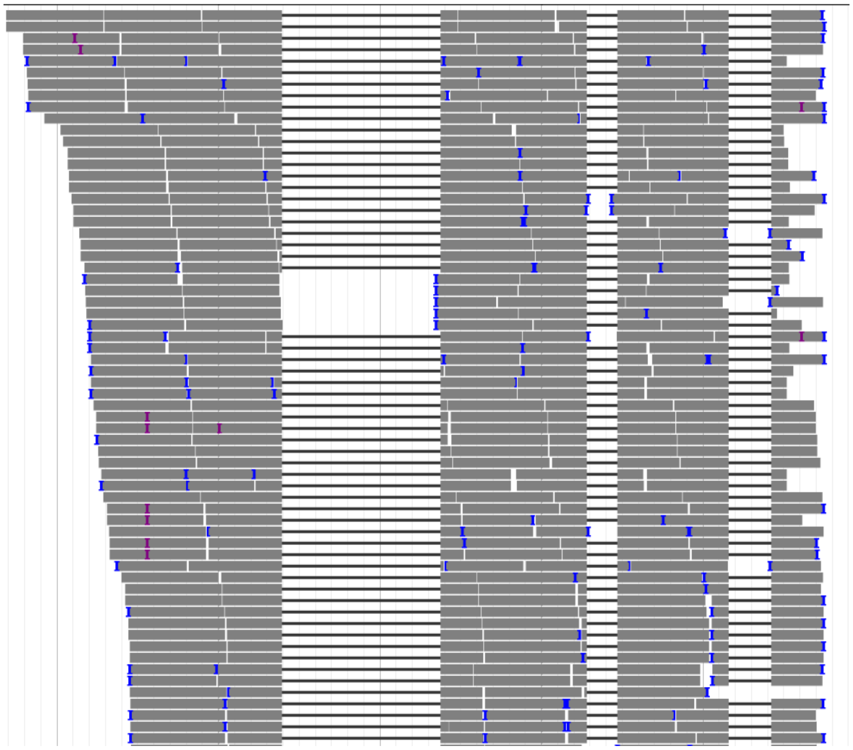

##### 1.2.2. BRAKER: Short-read based gene models

Prior to the expansion of computational biology, the identification of genes in a newly sequenced genome was an arduous and time-consuming process. In the early 2000s, the implementation of computer science into biological data analysis enabled the development of novel tools for automated gene prediction. To date, the BRAKER pipeline is the most successful gene prediction tool available. This tool leverages short-read RNA-seq data to train a probabilistic model that identifies protein-coding genes in a genomic sequence. The BRAKER model generated from the RNA reads in the previous figure is shown on the right. Although BRAKER is the best gene prediction tool currently available, it has several limitations. BRAKER is unable to predict the UTRs of a gene (red dot boxes). Additionally, BRAKER models often contain errors such as missing exons, fused genes, and split genes, among others. In the next section, we will learn how long-read sequencing technologies can be exploited to improve the quality of gene models.

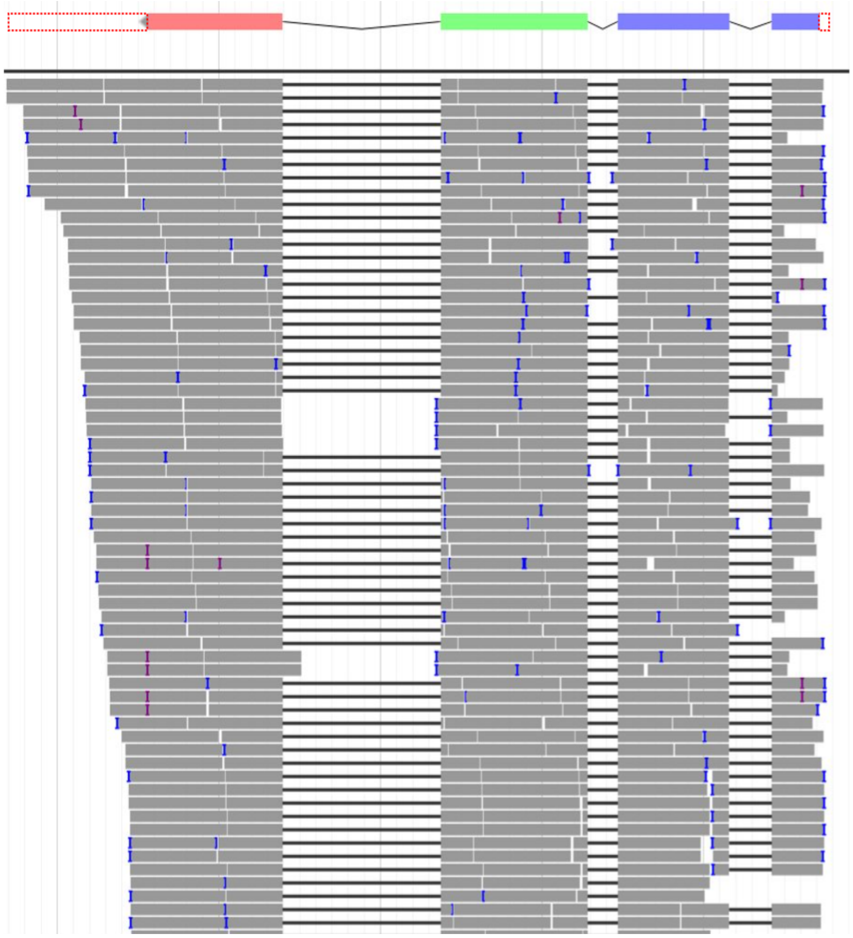

##### 1.3. Long-read sequencing

The development of long-read DNA sequencing technologies has allowed researchers to resolve complex genomic structures, such as repetitive regions and large rearrangements. As a result, new genomes with improved quality and completeness were generated for many organisms. Additionally, long-read RNA sequencing enabled the capture of full-length RNA transcripts with high accuracy. These technologies have the potential to improve over current short-read approaches for generating gene models.

###### 1.3.1. Single Molecule, Real-Time (SMRT) RNA Sequencing

In recent years, Pacific Biosciences (PacBio) released the Single Molecule, Real-Time (SMRT) sequencing technology. Similar to short-read RNA-Seq, the first step in SMRT RNA sequencing involves the preparation of a cDNA library from mRNA. However, unlike short-read RNA sequencing, SMRT sequencing supports average read lengths of 30,000 bases! We can sequence large cDNA fragments, which may encompass full-length mRNA transcripts. SMRTbell adapters are ligated to the end of each cDNA fragment. The SMRTbell adapters contain a small sequence that allows for primer annealing. A DNA polymerase then binds to the primer site on each adapter and begins the elongation of the cDNA fragment. As a result, the cDNA fragment is circularized, allowing for repeated sequencing passes. Each sequencing pass is called a subread. The consensus read obtained by collapsing all subreads obtained from a single cDNA fragment is called a circular consensus (CCS) read. CCS reads are then processed through the IsoSeq pipeline, which yields high-quality transcripts. An illustration of the SMRT sequencing method is shown below.

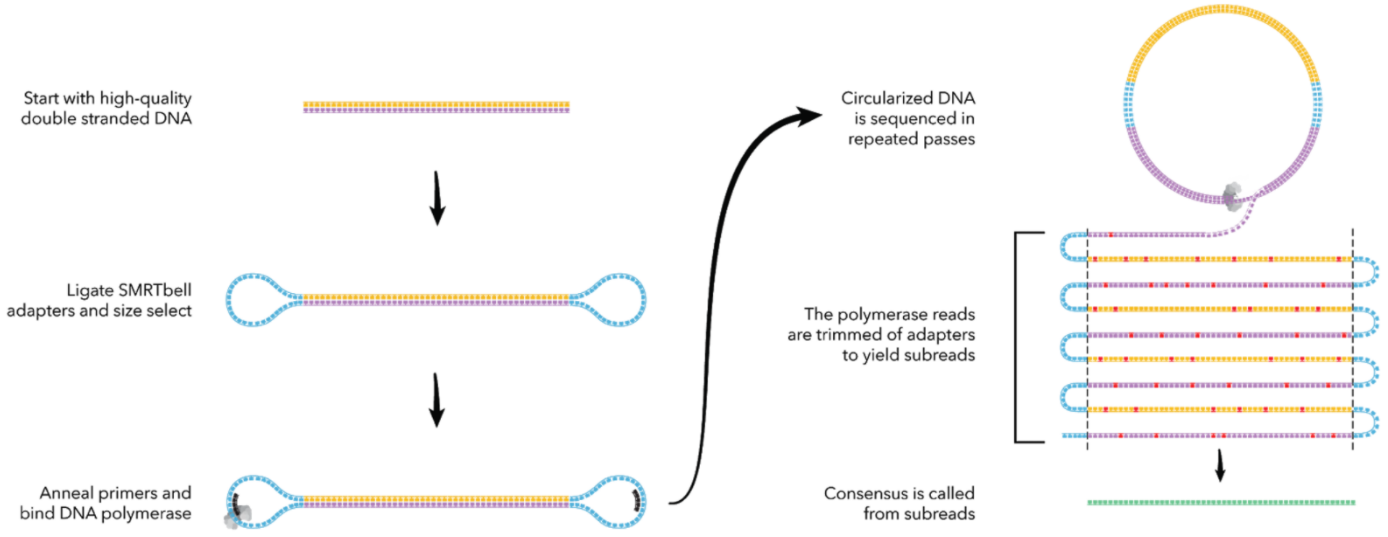

##### 1.3.2. StringTie: long-read transcriptome assembly

Processing CCS reads with the IsoSeq pipeline often yields a cluster of high quality transcripts, which may vary in length and exon composition (shown below). These clusters often represent incomplete transcripts (degraded mRNA) or alternative splice variants (isoforms) of a single gene.

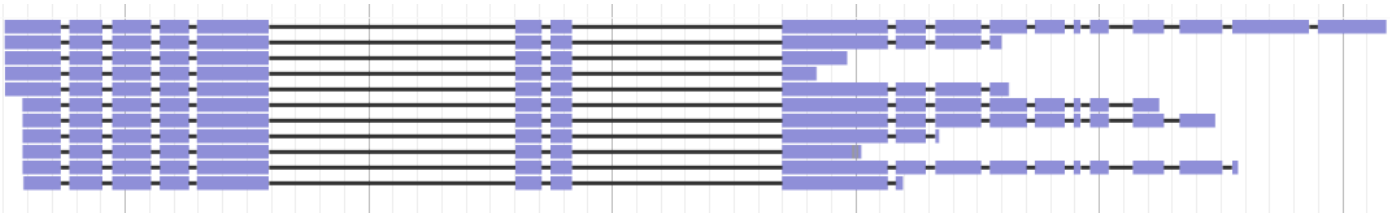

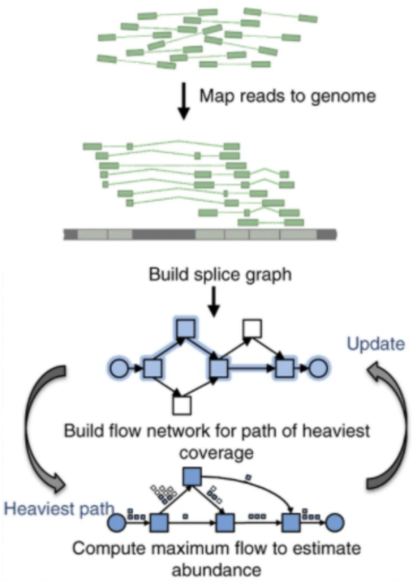


In order to identify an accurate gene model from the IsoSeq transcript cluster, we use StringTie. StringTie is a tool that allows for the assembly of a gene model from long-read alignments. StringTie incorporates a single alignment and generates a splice graph that includes all exon combinations in that alignment. The abundance of each possible path in the splice graph is then estimated using all read alignments mapping to the region of interest. The StringTie model undergoes several rounds of self-training, incorporating a new read alignment and updating the splice graph on each iteration. As a result, StringTie will identify and assemble the optimal transcript(s) from all the provided sequence alignments.

##### 1.4. Curation project aim

In this project, we will curate the complete set of genes of the *Caenorhabditis briggsae* strain QX1410. A highly curated gene set for this strain is necessary to further our studies of the evolutionary trajectory of *C. briggsae* and *Caenorhabditis* nematodes in general. The Andersen laboratory has generated a novel, high-quality genome assembly for QX1410. Additionally, the Andersen laboratory has obtained high-coverage Illumina RNA reads and IsoSeq reads for this strain. These sequencing efforts have allowed us to generate BRAKER and StringTie gene models for this strain. We will use these reads and gene models to curate each individual gene identified in the genome of this strain using the annotation software Apollo.

#### 2. Apollo

##### 2.1. Genome Browser Navigation Features

The Genome Browser allows you to travel through the coordinates of a genome in a fast and efficient way. The menu of the genome browser is shown below.

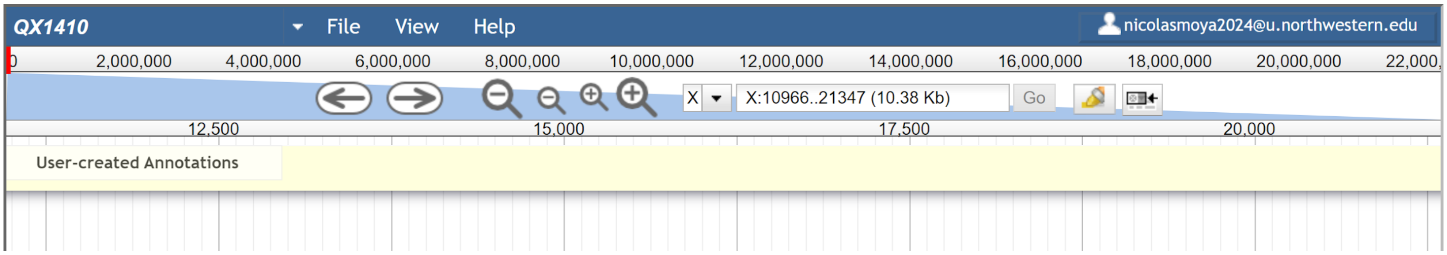


This menu possesses several features:

1. Chromosome Selection Box
    - This menu displays the current chromosome and allows you to move between chromosomes.

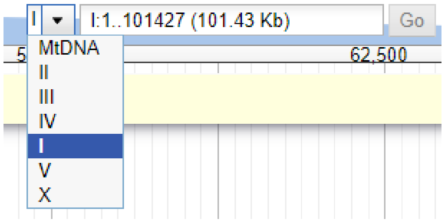


2. Chromosomal Coordinates Bar
    - This bar shows the coordinates of the entire chromosome you have selected, as well as a red box that highlights the specific region of the chromosome.

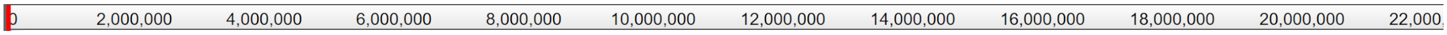


3. Coordinate Search Box
    - Here, you can see the start and end coordinates of the currently selected region. You can enter new coordinates, and click ‘Go’ to move rapidly throughout the genome.

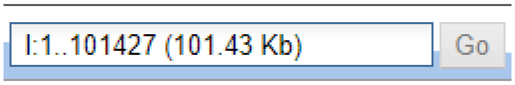


4. Region Coordinates Bar
    - This bar displays the coordinates of the selected region, and the reference coordinates in the bar are aligned with the lines in the browser window.

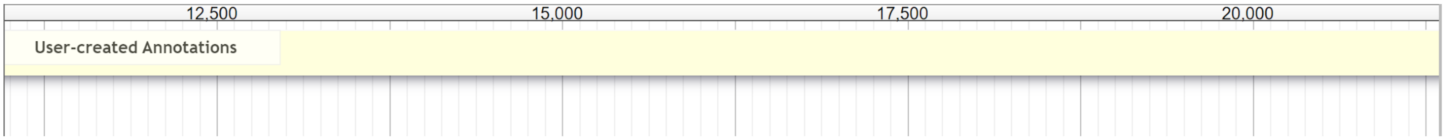

5. Lateral Movement Buttons
    - These buttons allow you to move right and left in the browser window in large steps.

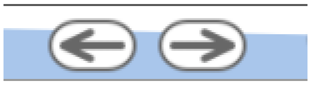

    - Alternatively, you can click-and-drag in any empty space of the browser window to move in either direction.

6. Zoom Buttons
    - These buttons allow you to zoom in and out of the current region.

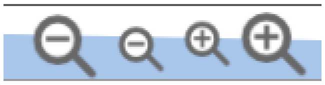

    - Additionally, you can click-and-drag in either the chromosomal coordinates bar or the region coordinates bar to zoom into a specific region.

Chromosomal Coordinates Bar

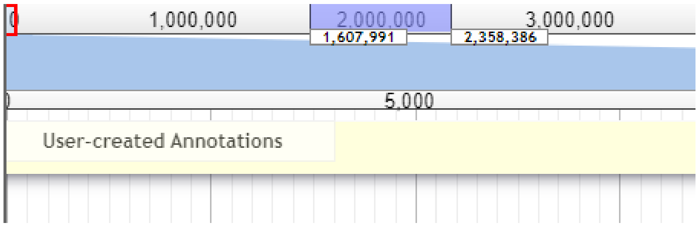


Region Coordinates Bar

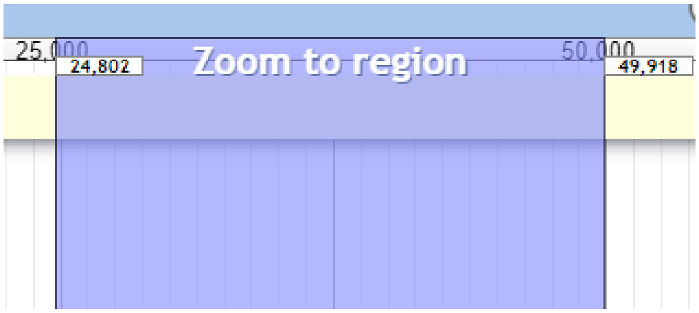


7. Reference Sequence Bar
    - When you reach the maximum zoom in a genomic window, a new bar will appear:

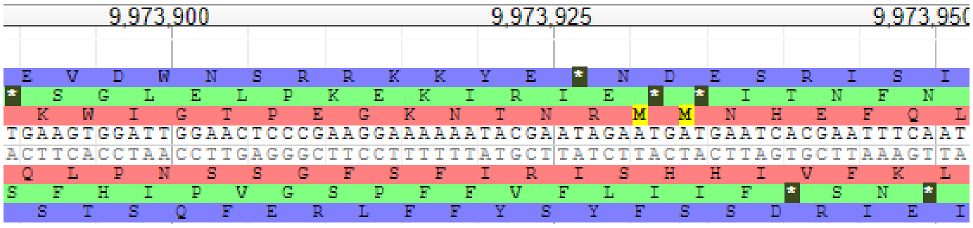


    - This bar displays the nucleotide sequence of the current region, as well as the three possible reading frames into which the nucleotide sequence can be translated.
    - The yellow boxes highlight methionine, the first amino-acid in any protein-coding sequence.
    - The asterisks represent stop codons.

8. Highlight Tool
    - By clicking on the highlight tool icon, you can highlight a specific region within the browser window. The highlighted region will appear in both chromosomal coordinates and region coordinates bars. You can only highlight one region at a time, and the marker will remain until removed. You can remove the highlighted region by clicking on the tool icon again.

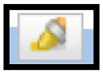

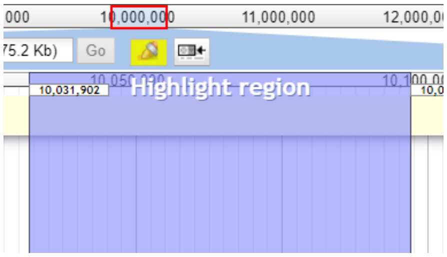


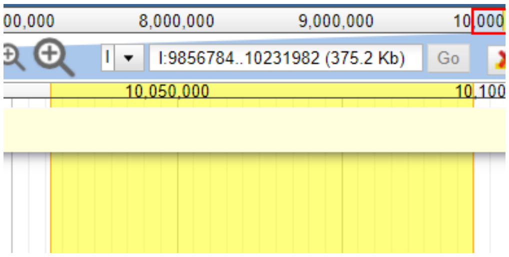


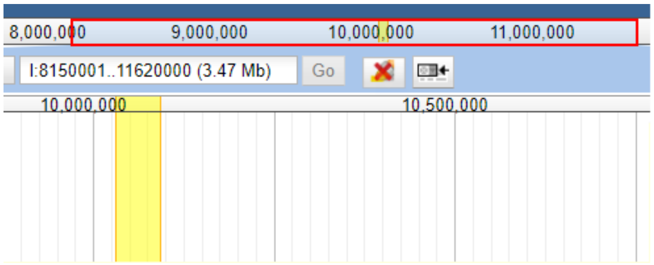

##### 2.2. Tracks

Tracks allow you to visualize sources of evidence in the Genome Browser. In the example below, four tracks were added to display BRAKER gene models, StringTie gene models, Illumina short-read sequence alignments, and IsoSeq high-quality transcript alignments. In this section, we will break down the features of each individual track.

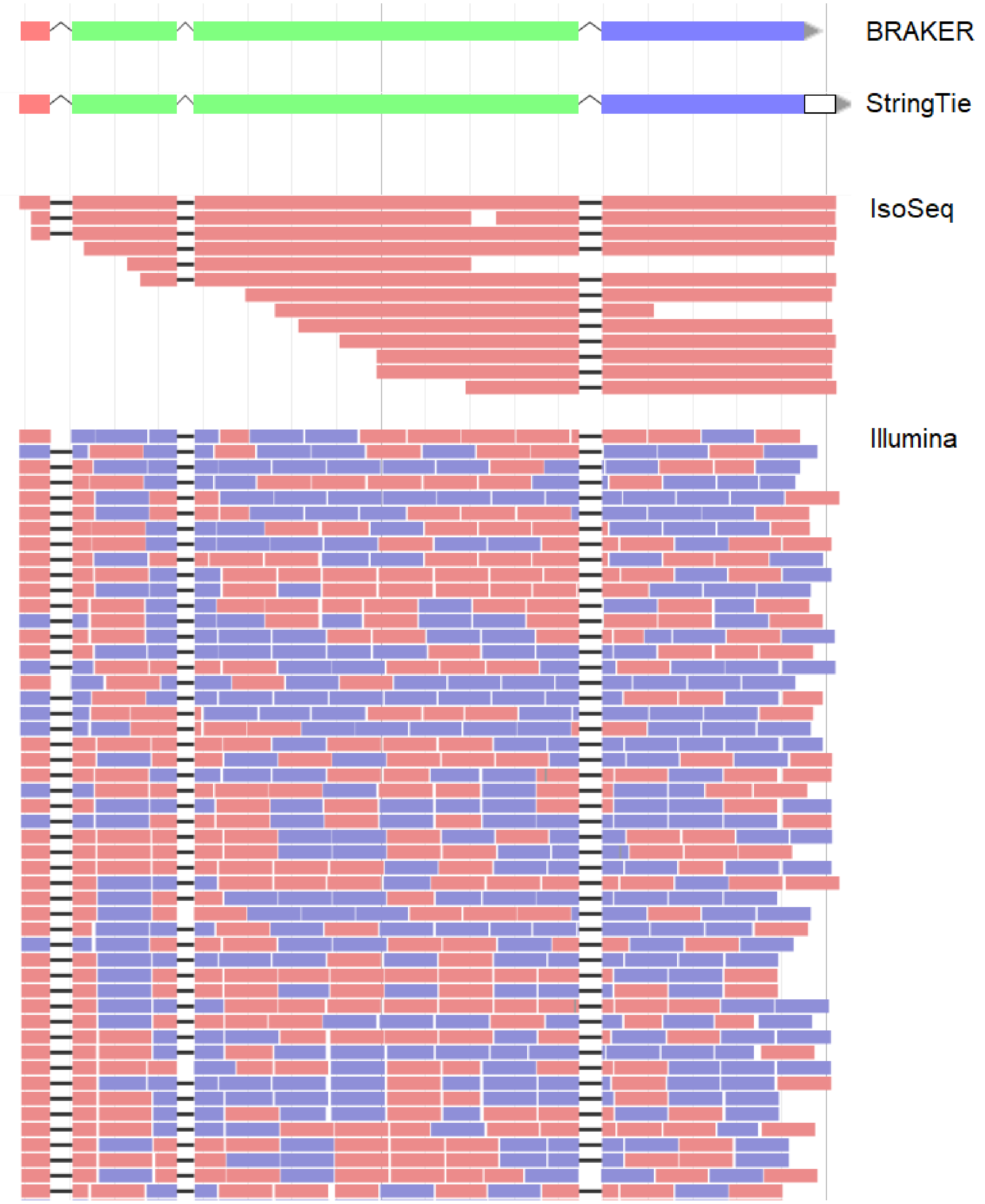

##### 2.2.1. BRAKER models

BRAKER models are contained within a single track. This track is draggable (can be directly added to the User-created Annotations space). BRAKER models do not possess UTRs. The orientation of the BRAKER model is observed in the orientation arrowhead in either terminal exon. Exons within the BRAKER model are colored by their protein reading frame, in reference to the Reference Sequence Bar. In the example below, the orientation arrowhead indicates the model is in the ‘Watson’ orientation (plus stranded). Additionally, the first exon is colored in red, which indicates the reference reading frame that is encoded by this exon.

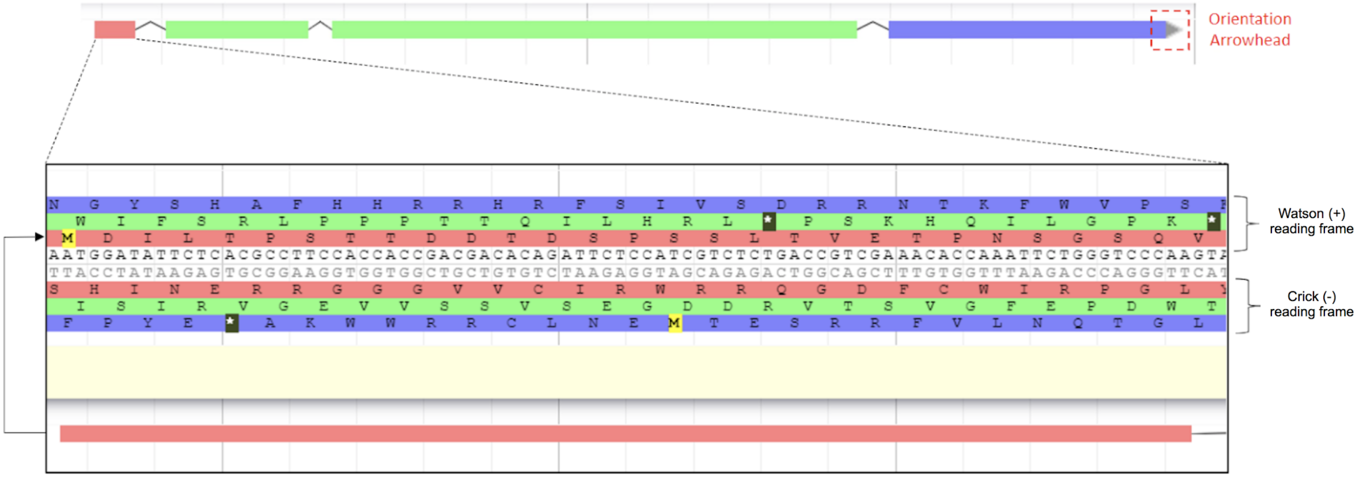


##### 2.2.2. StringTie models

The StringTie track is almost identical to the BRAKER track. The only difference is that StringTie models can possess UTRs.

##### 2.2.3. IsoSeq transcript alignments

IsoSeq transcript alignments are contained in two separate tracks. Both tracks are shown below. The first track track (IsoSeq HQ) is draggable, which allows IsoSeq transcripts to be moved directly into the User-created Annotation space. The first track also displays the orientation of each alignment (black arrows at the end). The second track (IsoSeq HQ - Spliced Display) is not draggable, but serves as a visual aid to easily identify the spliced regions of the transcript alignments in the genome browser. The second track also has marks that show deletions (blue) and insertions (purple). Both tracks mirror each other, as transcripts are found in the exact same order across both tracks.

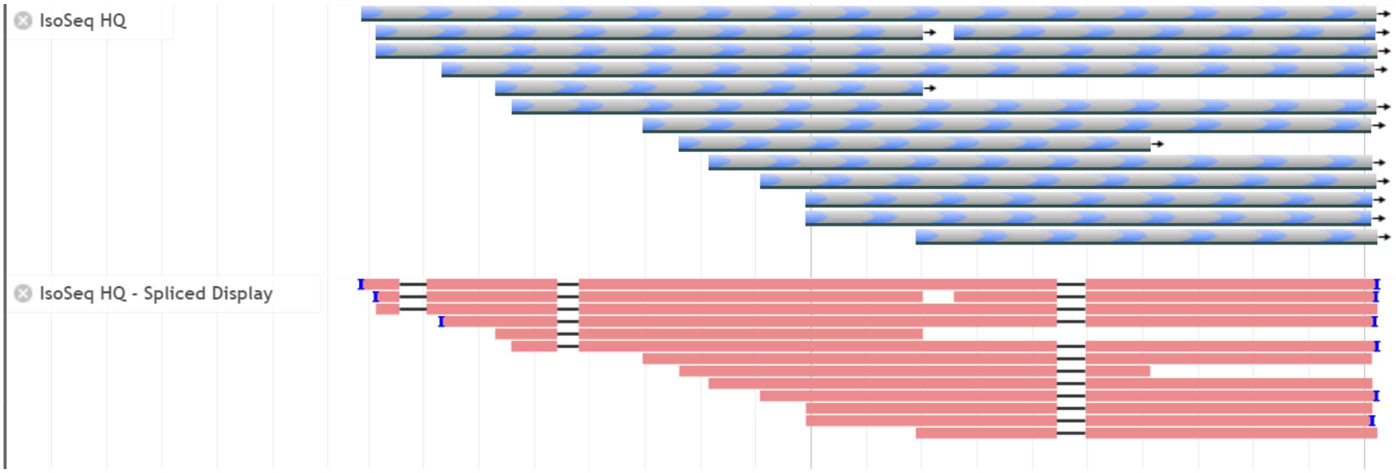

##### 2.2.4. Illumina short-read alignments

Illumina short-read alignments are also contained in two separate tracks. Both tracks are shown below. The features of these tracks are identical to IsoSeq transcript alignments.

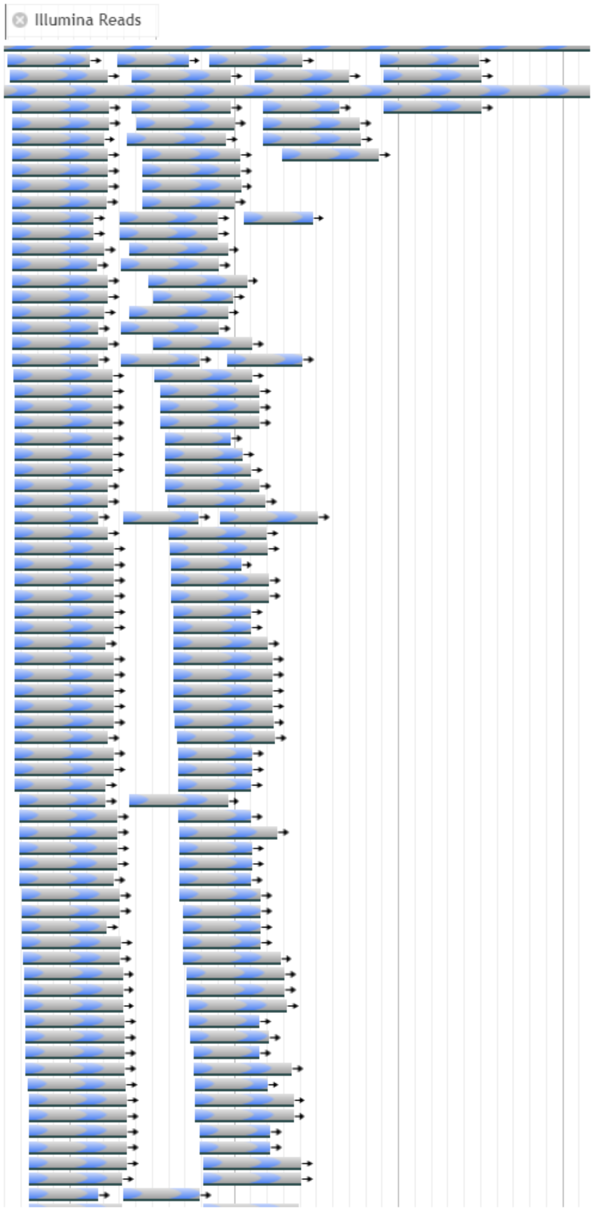


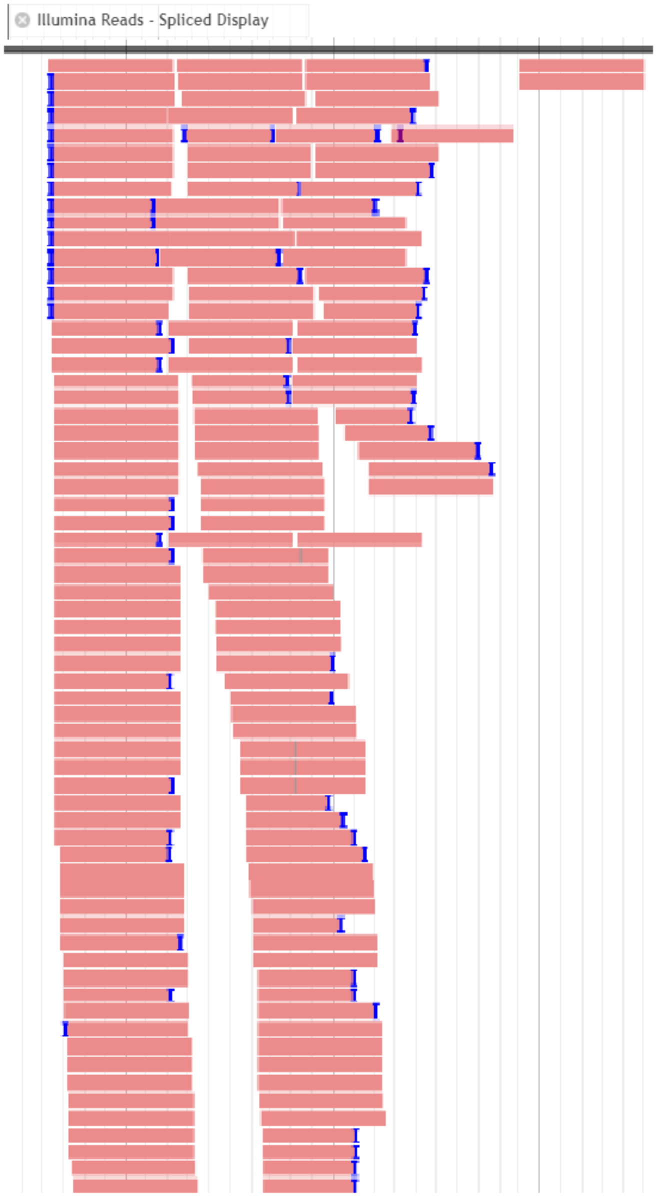

##### 2.3.4. Improving track visualization

There are a few handy features to improve the visualization of large data sets in Apollo. These features will be particularly useful to visualize lots of Illumina reads.

<u>Compact View</u>

You can compress the size of bars representing mapped reads in a BAM track (IsoSeq and Illumina). To do this, you can click on the drop-down menu on the track tag > Display mode > compact. This will allow you to fit more reads into the same window space.

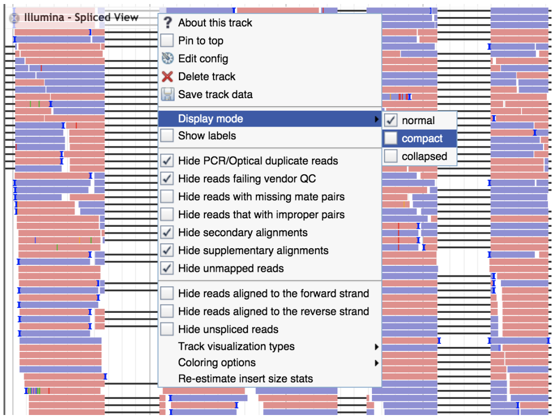


>>>>

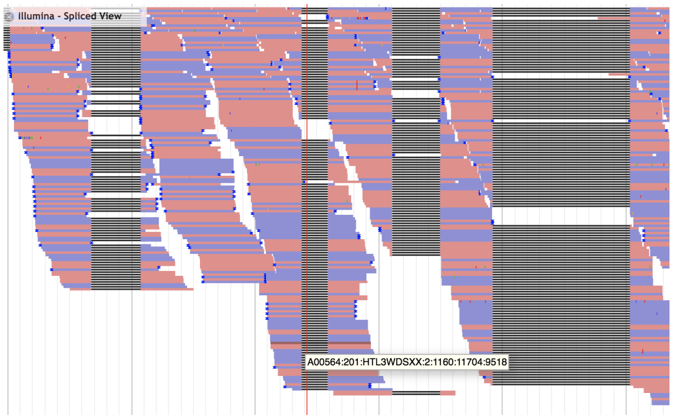


<u>Hide Unspliced Reads</u>

In some cases, collapsed view is not sufficient to visualize all reads that are mapping to a portion of the gene. For example, if you are looking for Illumina support for a rare isoform and you are only concerned about reads mapping to a specific intron, you can hide unspliced reads. This will only show reads that are spliced, allowing you to visualize reads that span the boundaries of introns. To do this, you can click on the drop down menu of the track tag > Hide unspliced reads.

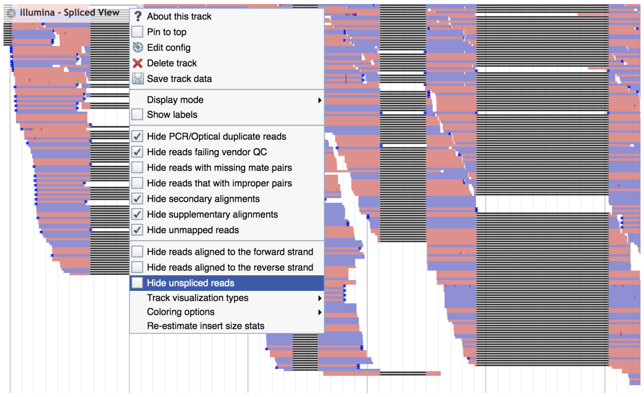


>>>>

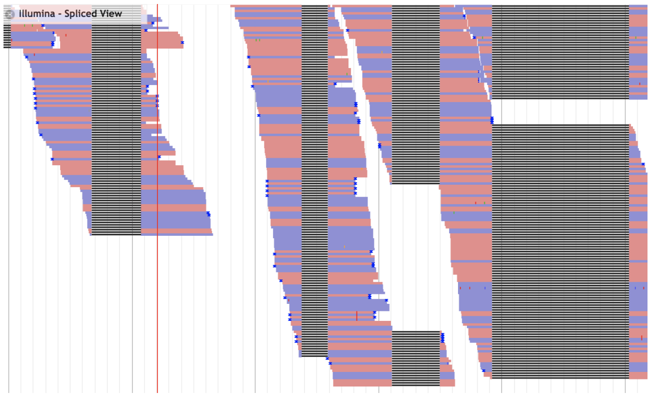


**NOTE:** Please ensure you **turn off** this option after curating the gene of interest. Hiding unspliced reads may make it seem like there is no support for an exon, and it may lead to mistakenly splitting a gene. It is crucial that you move onto the next gene **only after turning off** this feature.

##### 2.3. User-created Annotations

To start the curation process, we will need to add evidence to the ‘User-created Annotations’ space from each individual track.

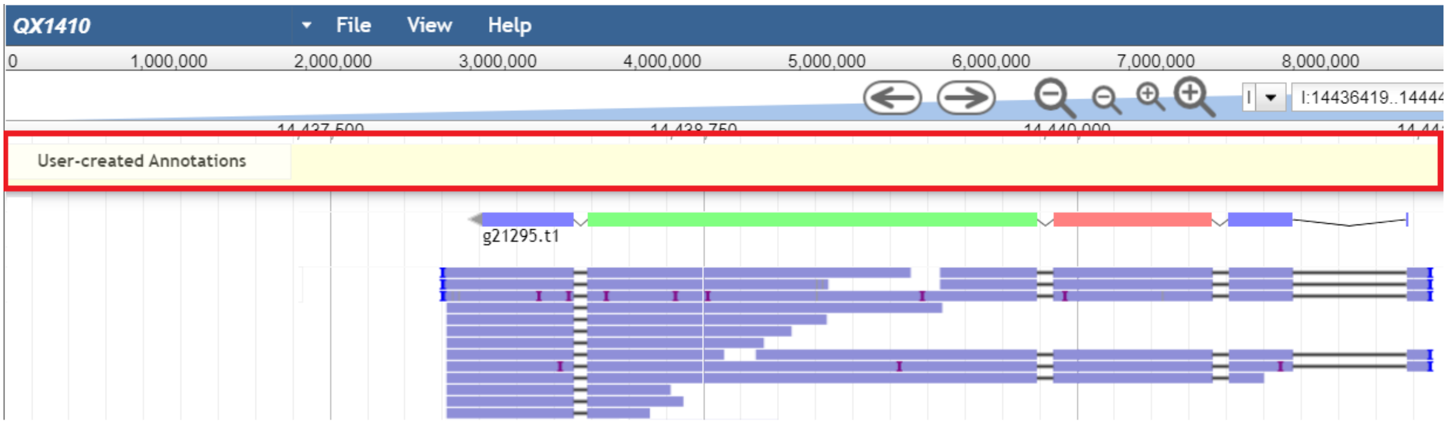


###### 2.3.1. Adding Annotations

To add features to ‘User-created Annotations’, double-click on a feature from a track (when selected, a red outline will appear around the entire feature):

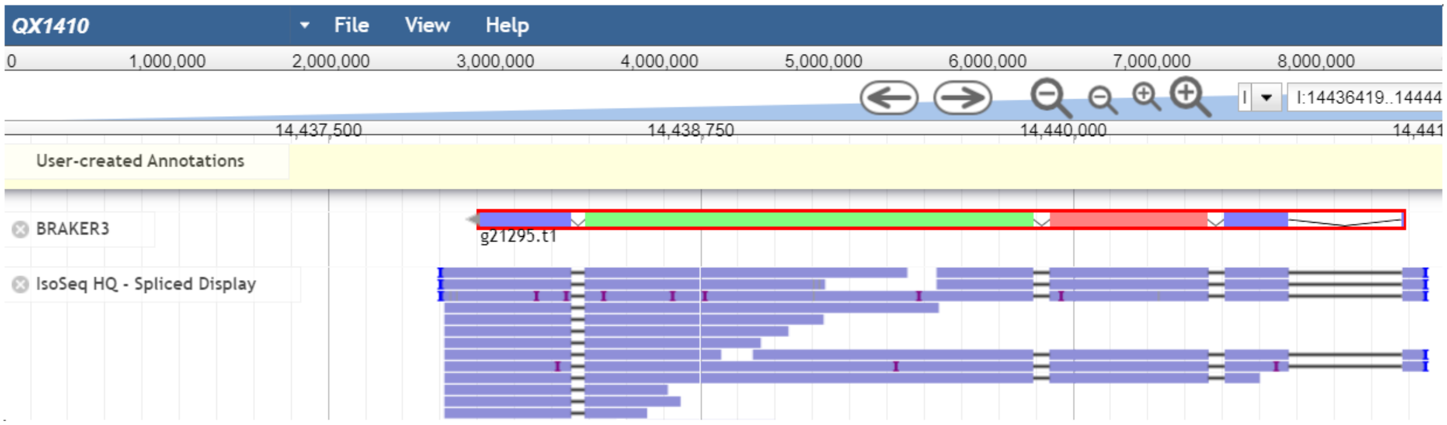


Then, drag-and-drop the feature in the ‘User-created Annotation’ space:

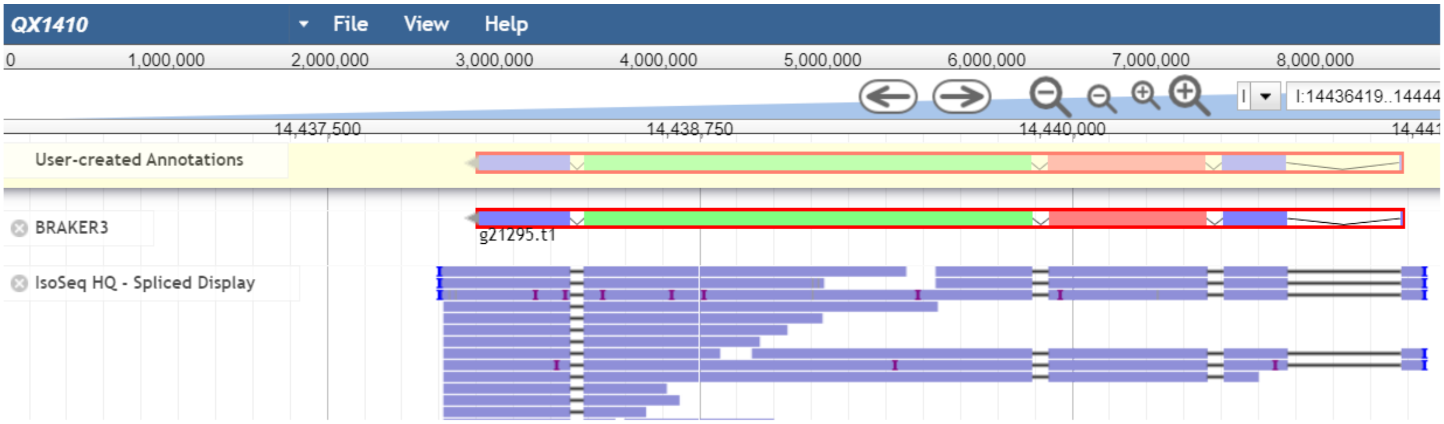

Once dropped, the feature will appear in the 'User-created Annotations' space. The first feature is the primary feature.

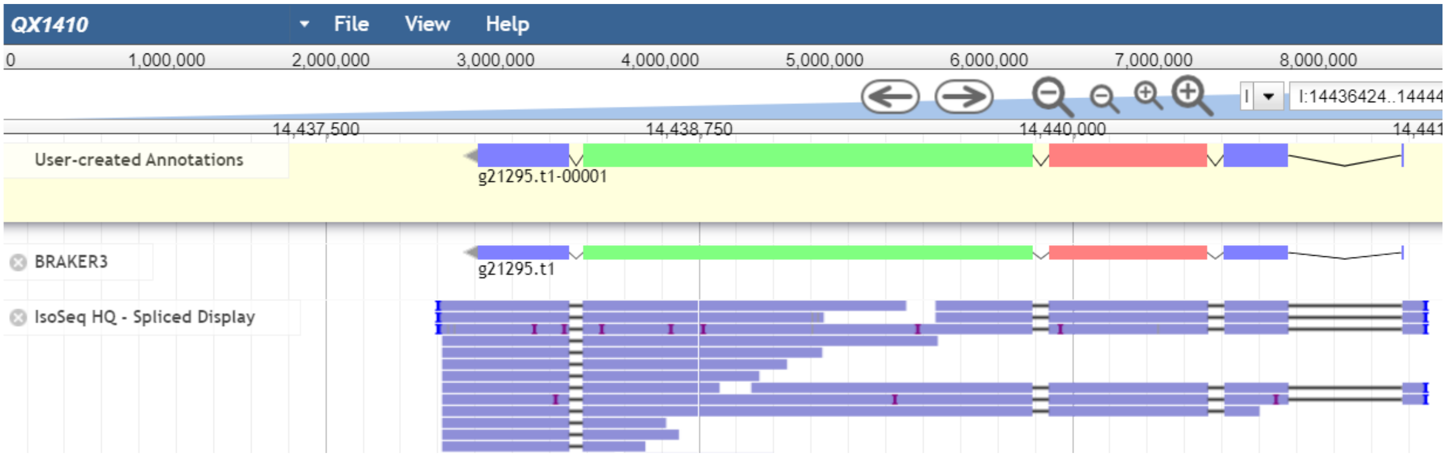


You can add multiple features from any draggable track into the 'User-created Annotations' space. Any additional features added after the primary feature are referred to as secondary features.

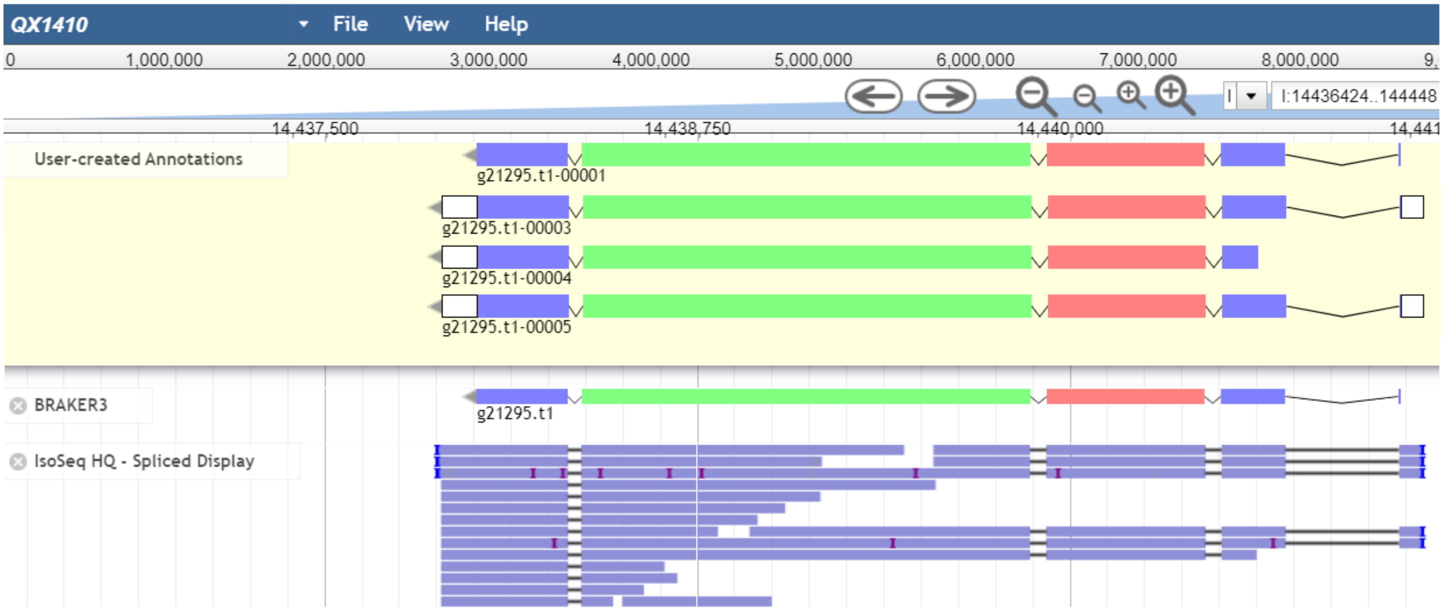


##### 2.3.2. Removing annotations

To remove an annotation from the User-created Annotations space, double-click on the annotation to select it (a red box will appear around the annotation).

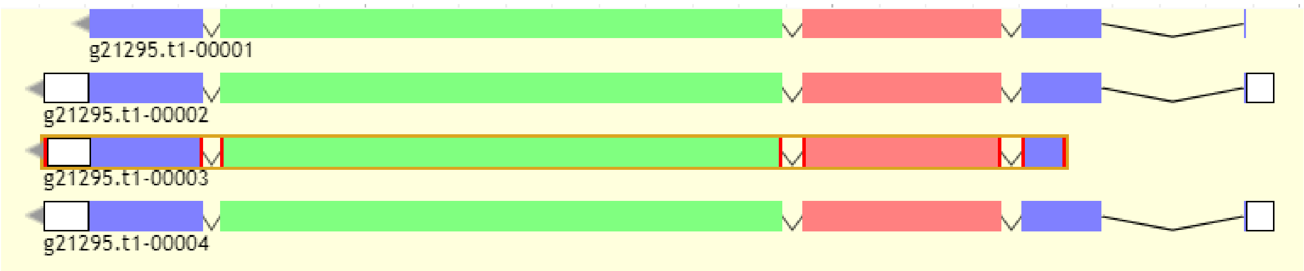

Then, right-click on any section of the annotation, and select the ‘Delete’ option.

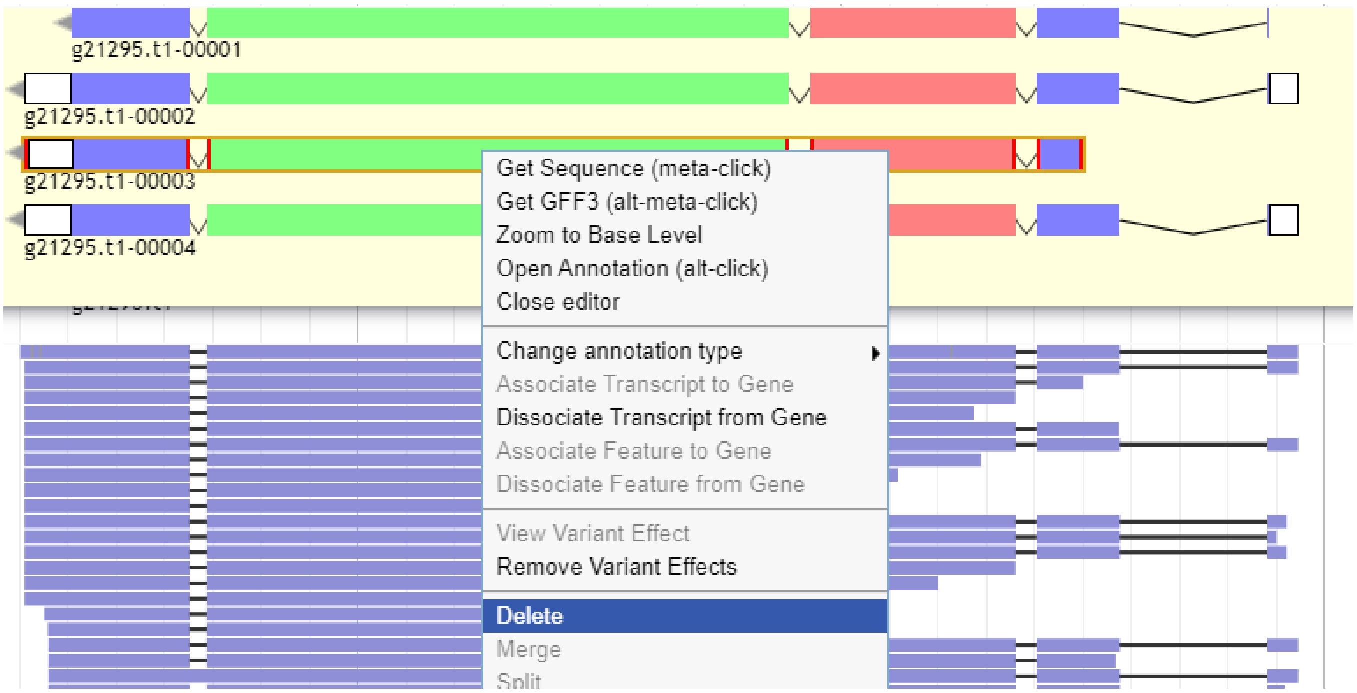


##### 2.3.3. Renaming annotations

To rename an annotation, select the annotation (gene) in the ‘Annotations’ tab of the Apollo menu, and type in the alternative name in the ‘Name’ box.

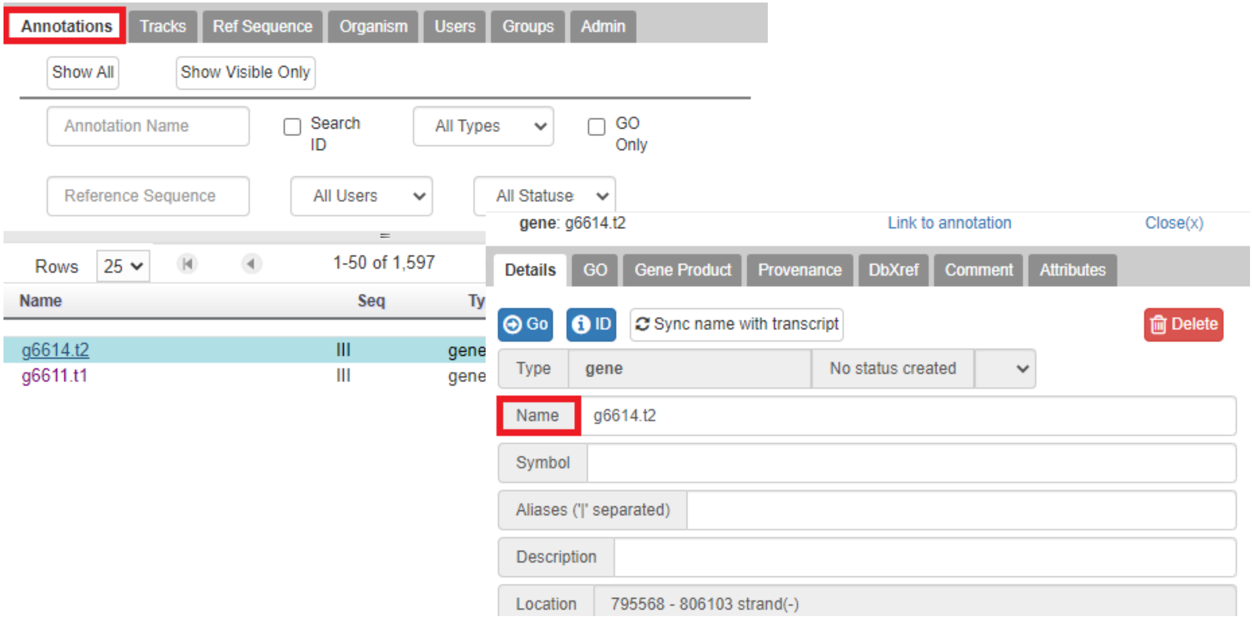

Then, click on the drop down menu on the right of the annotation entry (in the list of annotations)

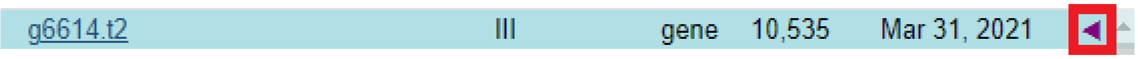


And rename each individual transcript, using the same method shown above.

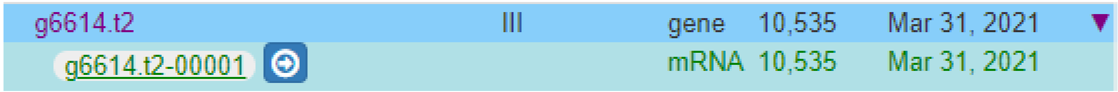


**NOTE:** If you are trying to rename a gene model that is between two genes with names that have consecutive numbers (e.g: renaming a StringTie model that is between g1454.t1 and g1455.t1), you should set the name of the intermediate model by adding an ‘.i.’ in between the gene number and transcript number of the previous model (e.g: g1454.i.t1).

##### 2.4. Gene Curation

As a general rule, three main questions need to be answered to curate a gene:

1. What evidence is available?
2. Do any conflicts exist between the sources of evidence?
3. How can I resolve these conflicts?

Several intermediate steps will help you answer these questions and are summarized in the following workflow:

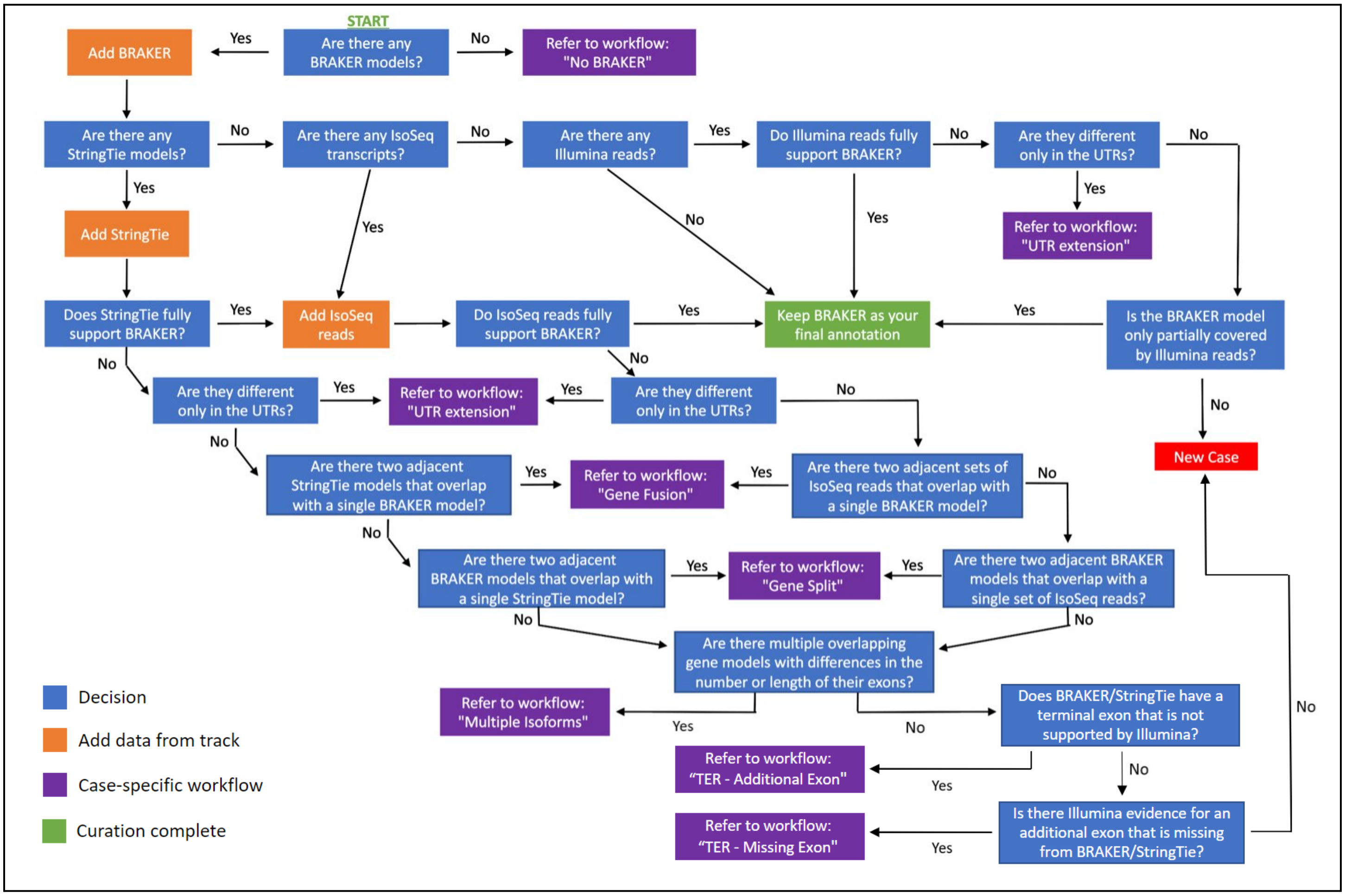


Considering the complexity of this workflow, we will break down each potential pathway by looking at specific cases.

##### 2.4.1. Only BRAKER

The simplest case would be identifying a *de novo* BRAKER prediction without any supporting evidence:

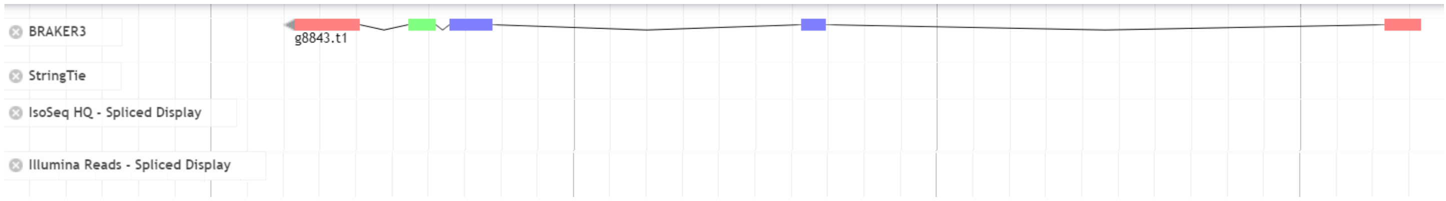


Following the General Workflow:

1. Are there any BRAKER models? (Yes)
2. Add BRAKER model
3. Are there any StringTie models? (No)
4. Are there any IsoSeq transcripts? (No)
5. Are there any Illumina reads? (No)
6. Leave the BRAKER model as your final curated model in the User-created Annotations space.

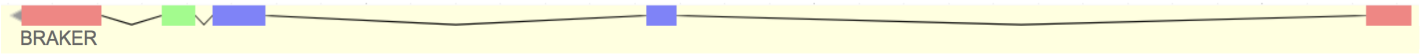


7. Fill curation metadata (Section 2.5.), and move onto the next gene.
   a. Metadata code for this case: 'Simple - B'

Note: In this and all following cases, if there are two or more BRAKER models, curate each BRAKER model independently.

##### 2.4.2. BRAKER and limited Illumina reads

A small variation to the previous case involves identifying a BRAKER prediction that has partial support from a few Illumina read alignments:

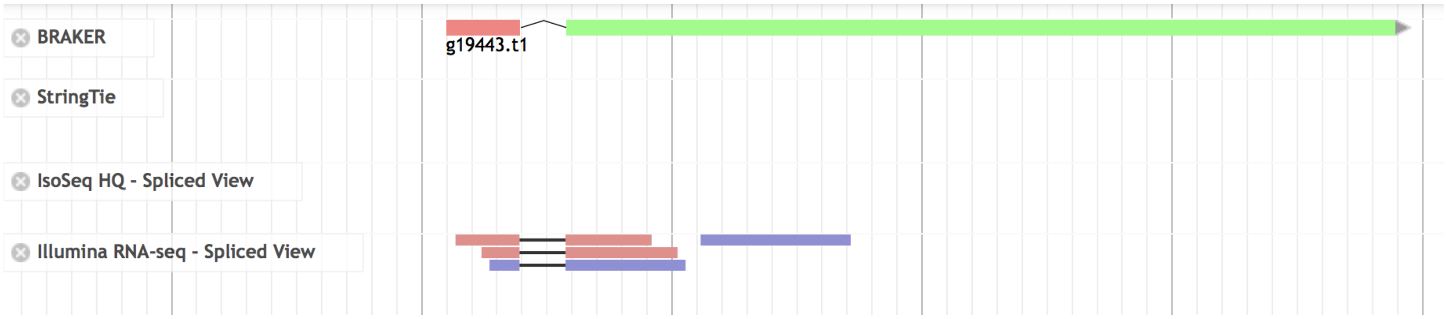


Following the General Workflow:

1. Are there any BRAKER models? (Yes)
2. Add BRAKER model
3. Are there any StringTie models? (No)
4. Are there any IsoSeq transcripts? (No)
5. Are there any Illumina reads? (Yes)
6. Is the BRAKER model only partially covered by Illumina reads? (Yes)
7. Leave the BRAKER model as your final curated model in the User-created Annotations space.

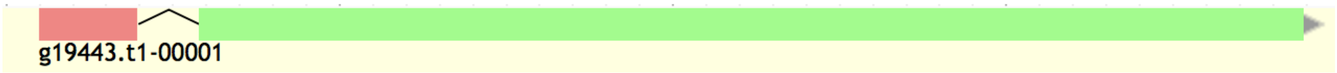


8. Fill curation metadata (Section 2.5.), and move onto the next gene.
   a. Metadata code for this case: 'Low Coverage - B'

##### 2.4.3. UTR extension

Another simple case involves identifying BRAKER and StringTie models with supporting long and short reads that only differ in the UTRs:

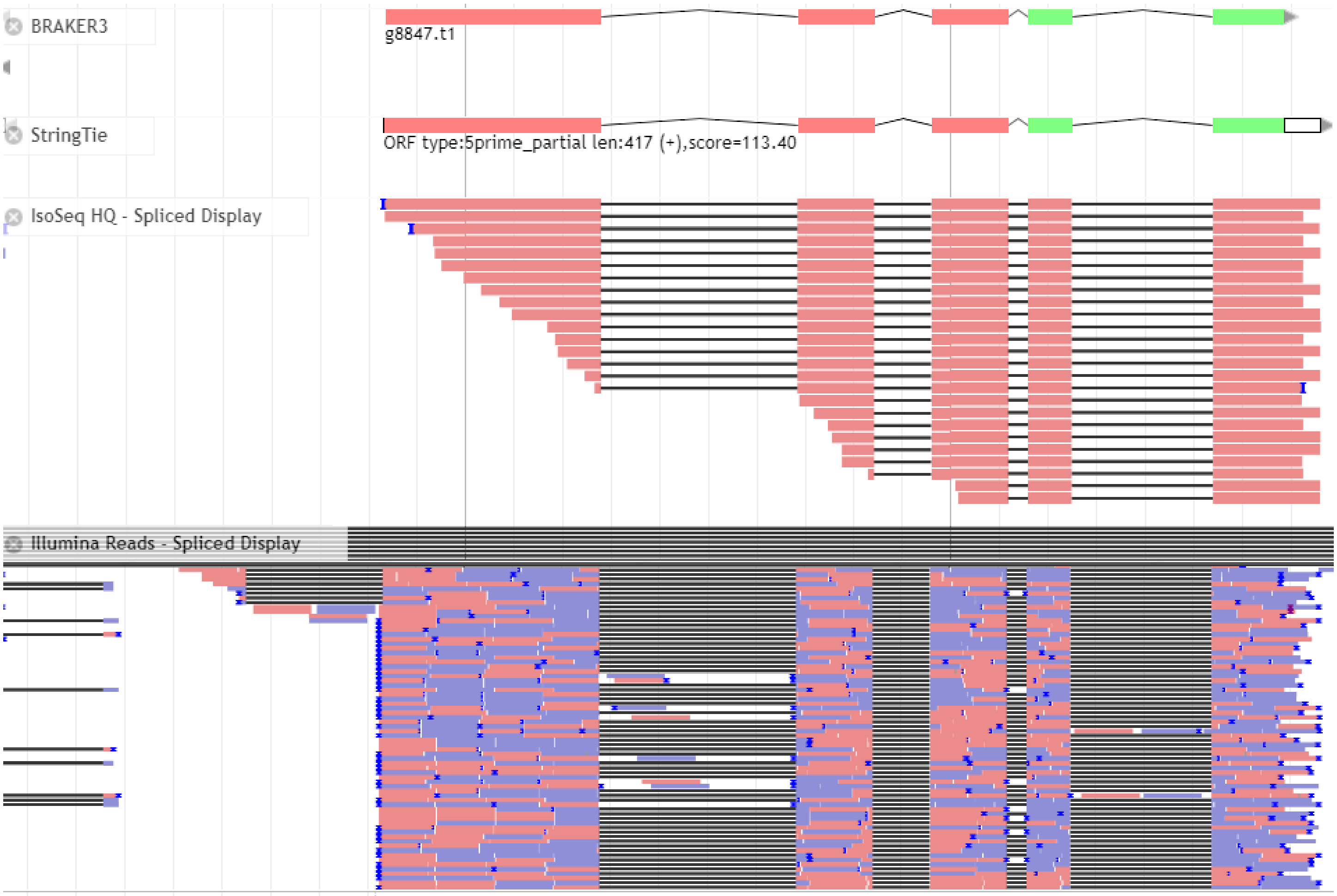


Following the General Workflow:

1. Are there any BRAKER models? (Yes)
2. Add BRAKER model
3. Are there any StringTie models? (Yes)
4. Add StringTie model
5. Does StringTie fully support BRAKER? (No)
6. Are they different only in the UTRs? (Yes)
7. Refer to workflow: ‘UTR Extension’

Following **UTR Extension** Workflow:

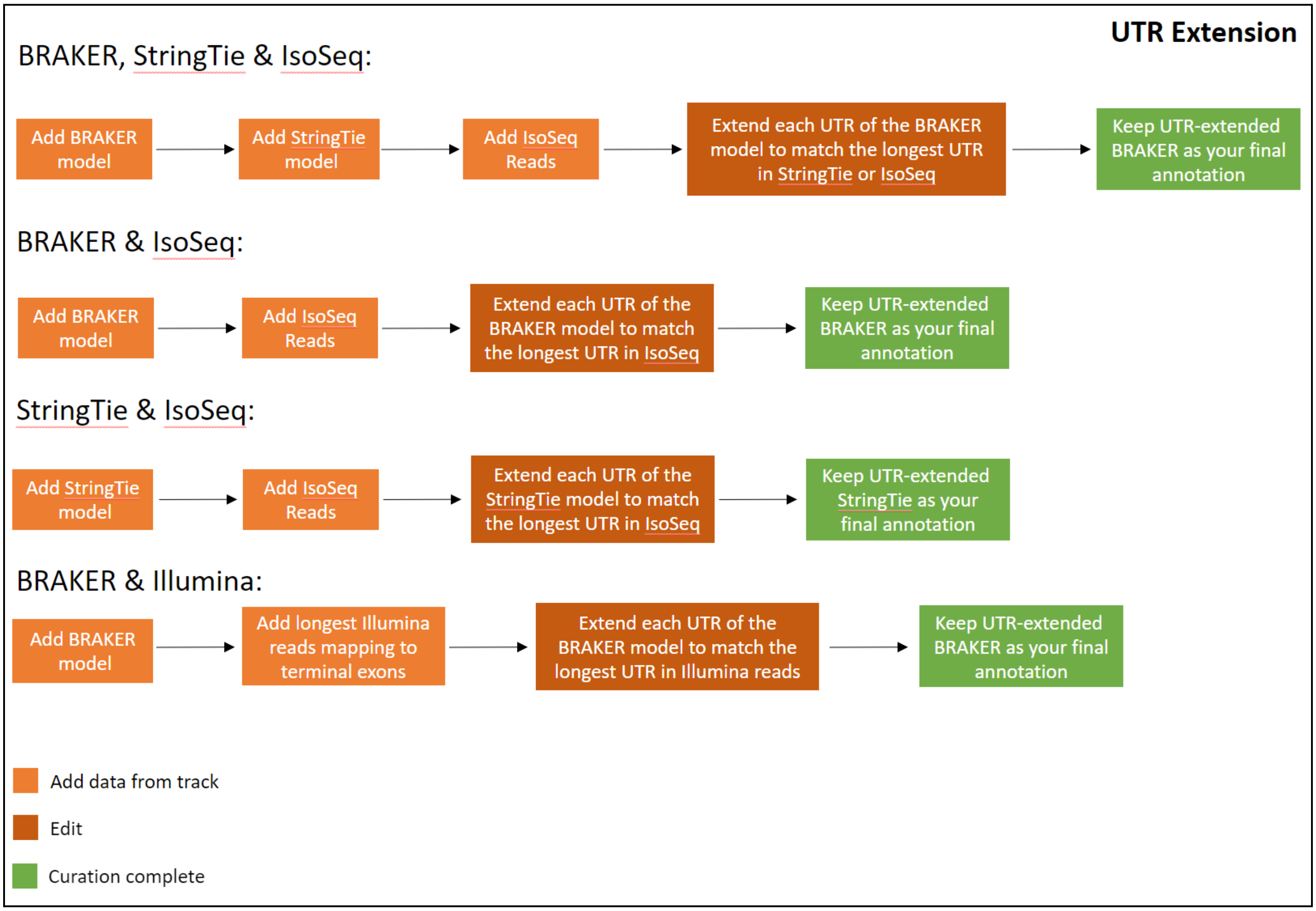


1. BRAKER and StringTie models are already added
2. Add IsoSeq reads
3. Extend the UTR of the BRAKER model to match the longest UTR in the StringTie or IsoSeq model

To extend the UTR we:

1. Select the BRAKER terminal exon (left-click the exon)

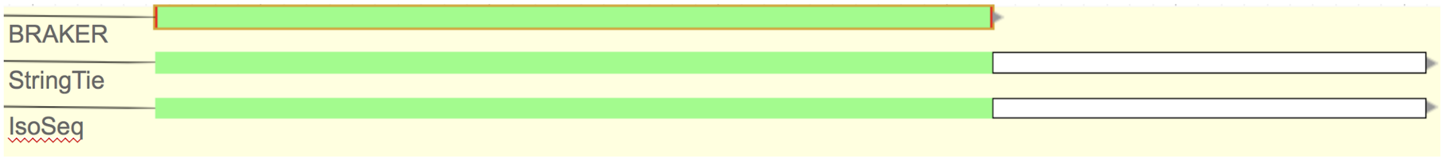


2. Place the cursor over the edge of the terminal exon that will be extended
3. Click, hold, and extend the edge until it matches the UTR of the StringTie model and/or IsoSeq transcript

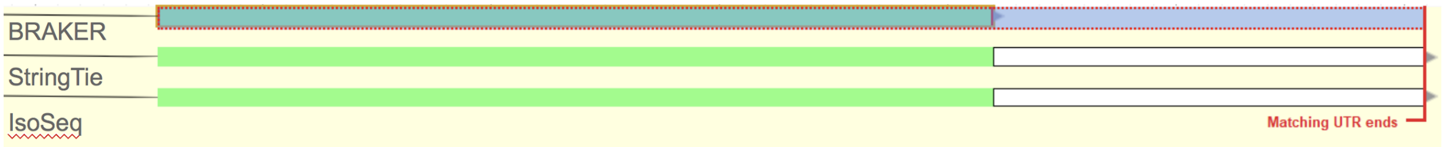

- To ensure the newly extended UTR perfectly matches the StringTie model and/or IsoSeq transcript, zoom to the nucleotide level and confirm that all models end on the same nucleotide

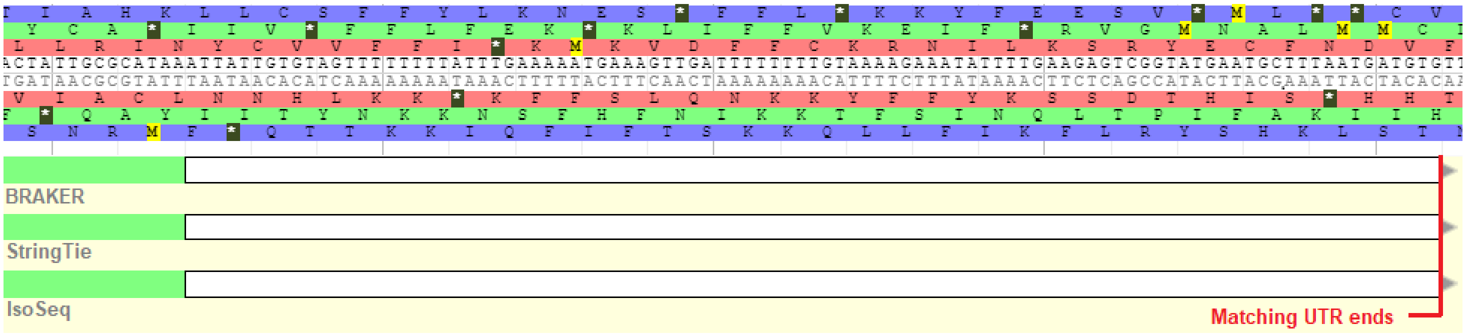


4. Leave the UTR-extended BRAKER model as your final curated gene model in the User-created Annotations space.

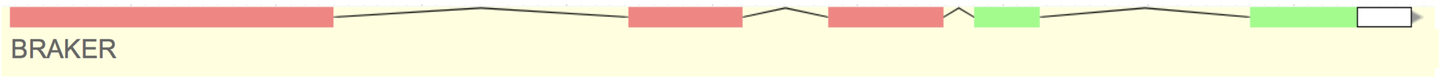


5. Fill curation metadata (Section 2.5.), and move onto the next gene.
    - Metadata code for this case: ‘UTR Extension - B:S:IS’

Alternative **UTR Extension** Workflow:

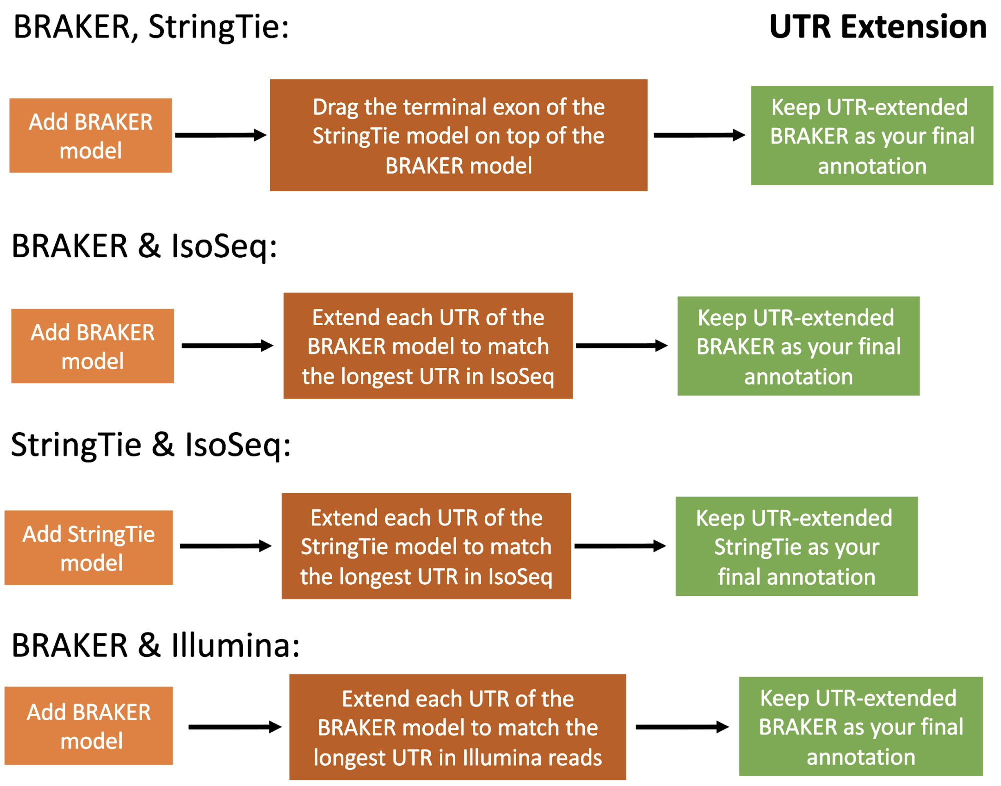


Add data from track

Edit

Curation complete

In all cases, StringTie models will always predict UTRs based on the longest IsoSeq read. If you have both BRAKER and StringTie models and there is no further extension to be done based on the Illumina data, you can extend the UTR of a BRAKER model by:

1. Add BRAKER into the User-created annotation space

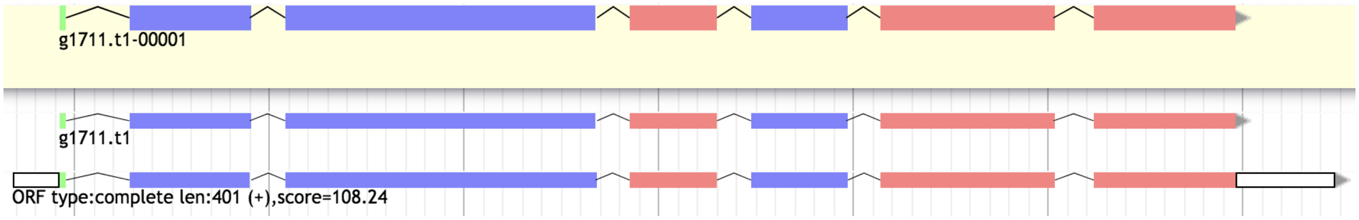


2. Select the terminal exon of the StringTie model (from the track)

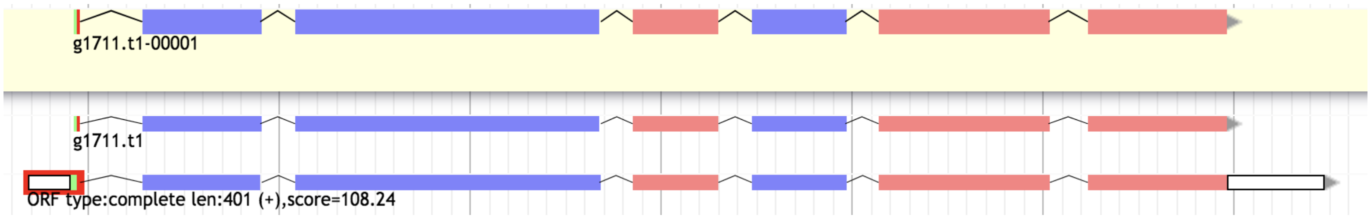


3. Click, hold and drag the selected exon on top of the BRAKER model (a green outline will appear)

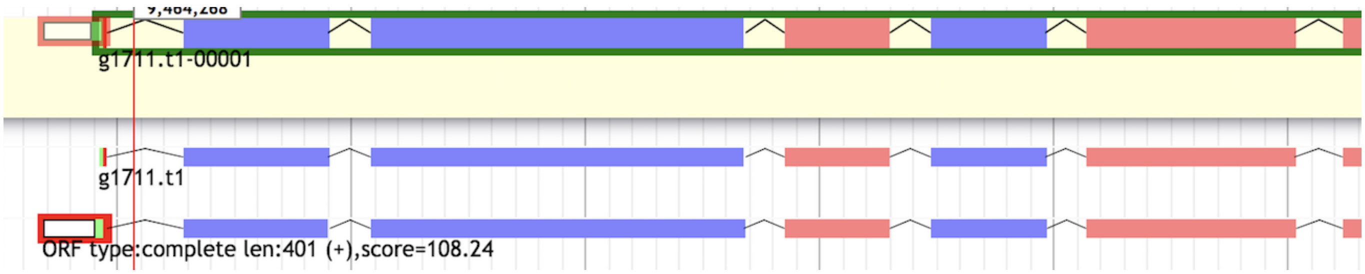


4. Release the exon, and the UTR will be automatically extended.

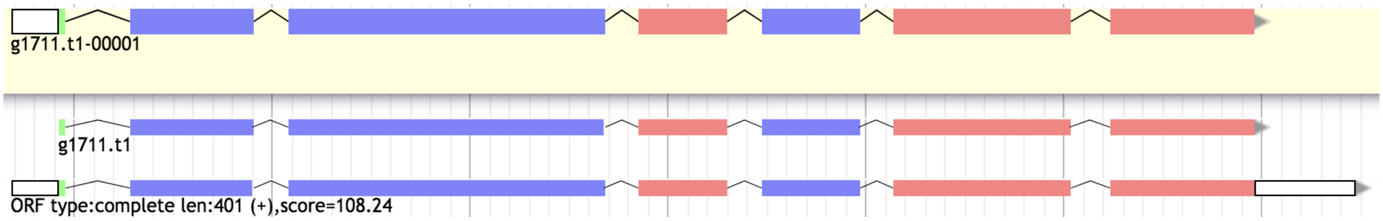

5. If you are extending a 5’ UTR, ensure that the translation start site has not been shifted
    - If the translation start site has been shifted, set it back to its original position by right clicking the ‘A’ of the ‘ATG’ in the original translation start site and then click on ‘Set translation start’

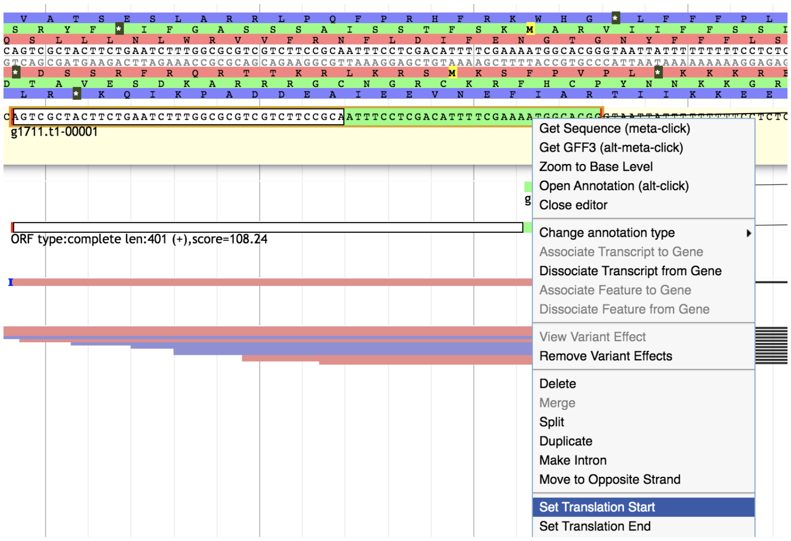


    - You can also adjust the translation start via Apollo’s coding sequence prediction tool, which involves selecting the entire transcript > right click > Set optimal ORF

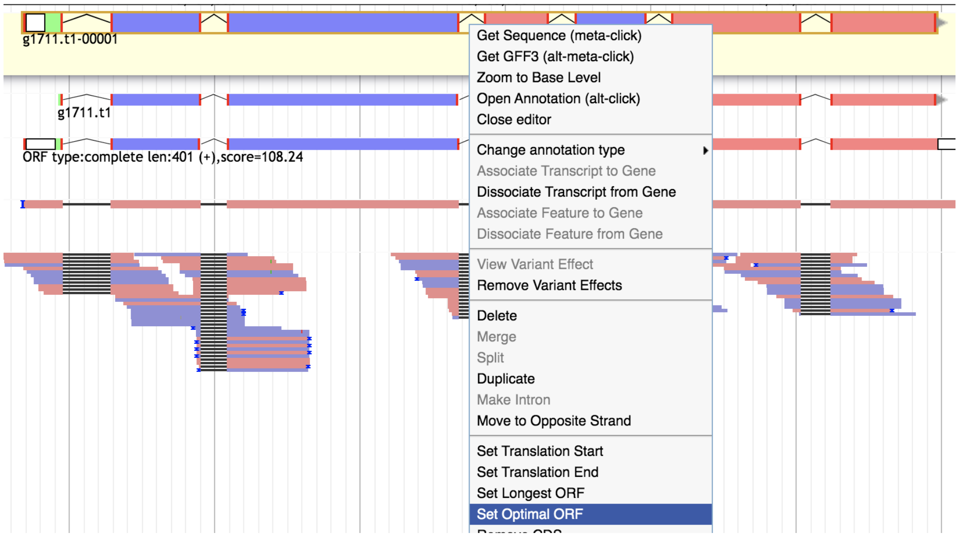

##### 2.4.4. Gene split

A more complex case involves identifying a potential gene split, where two adjacent BRAKER models overlap with a single StringTie model or IsoSeq transcript:

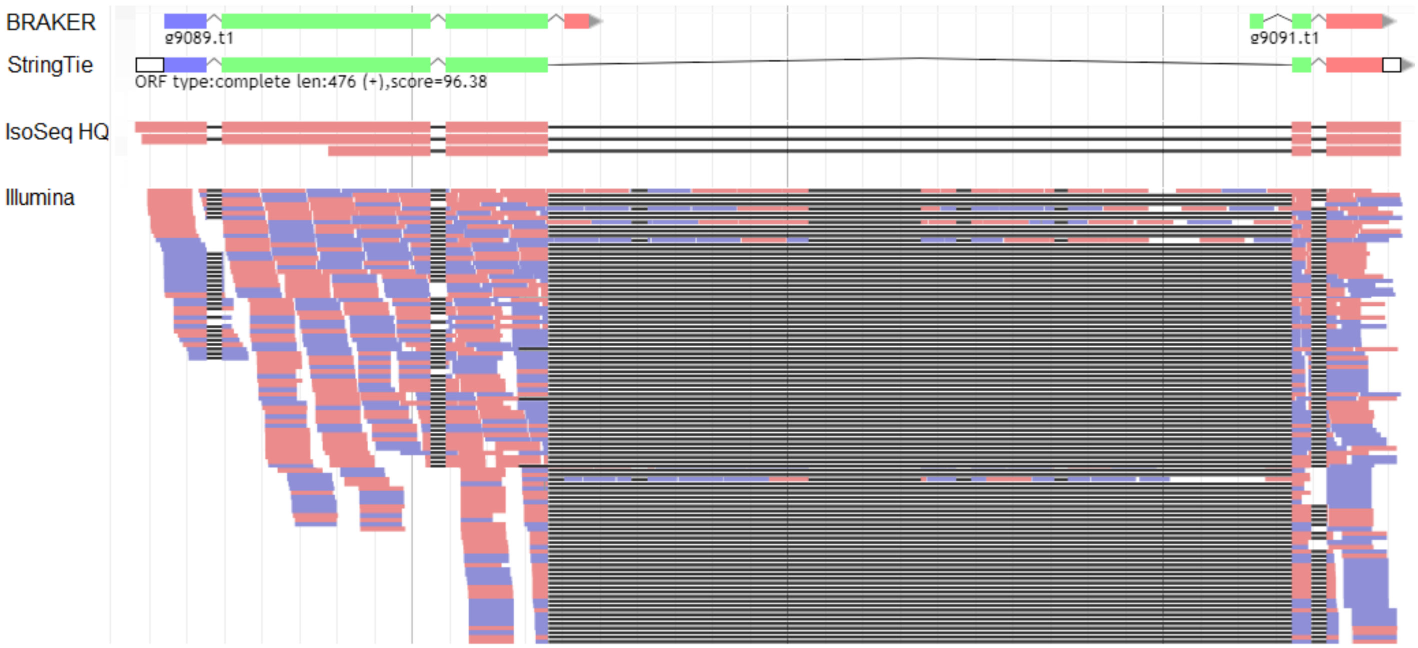


Following the General Workflow:

1. Are there any BRAKER models? (Yes)
2. Add BRAKER
3. Are there any StringTie models? (Yes)
4. Add StringTie
5. Does StringTie fully support BRAKER? (No)
6. Are they different only in the UTRs? (No)
7. Are there two adjacent StringTie models that overlap with a single BRAKER model? (No)
8. Are there two adjacent BRAKER models that overlap with a single StringTie model? (Yes)
9. Refer to workflow: “Gene Split”

Following the Gene Split workflow:

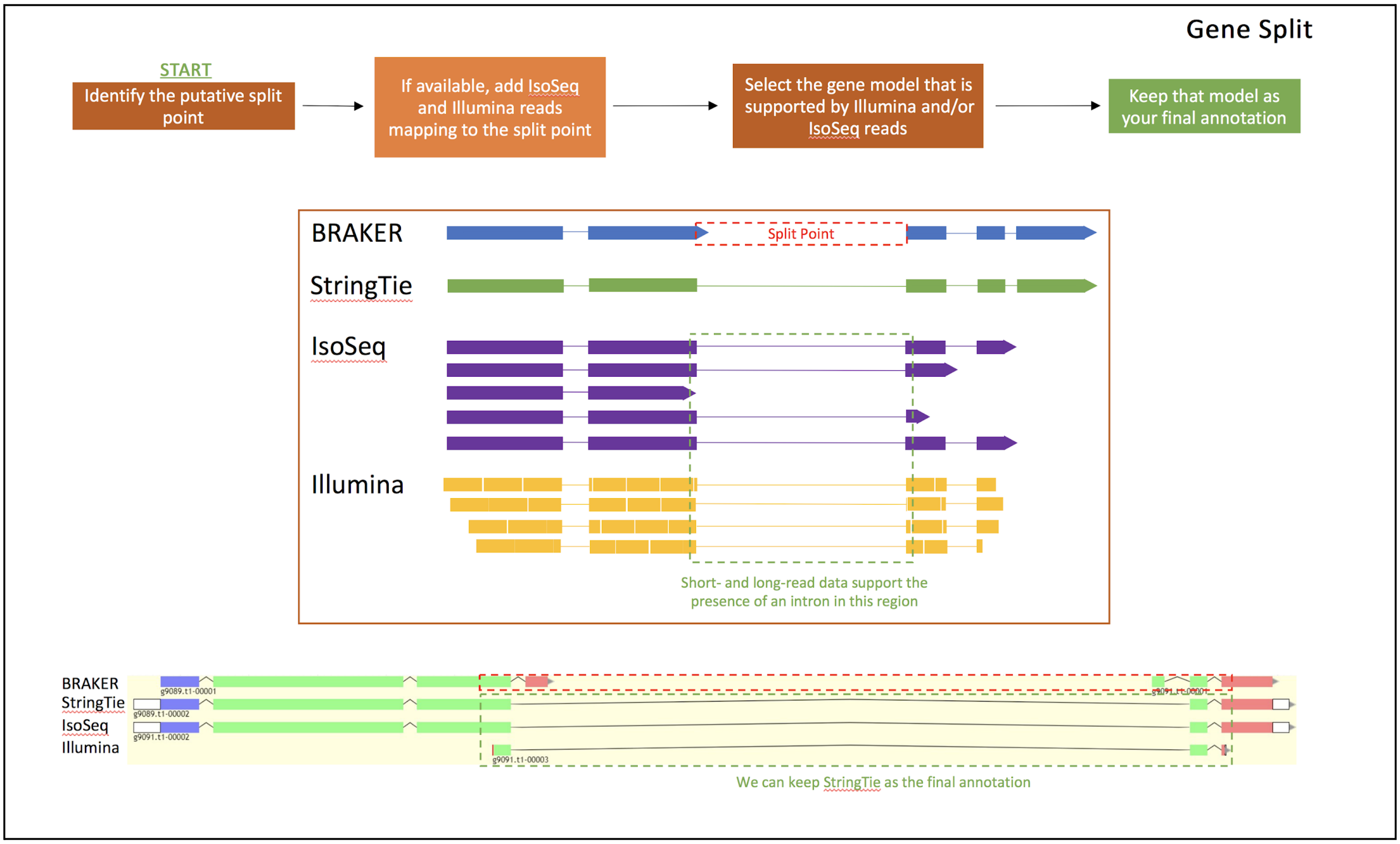


1. We added BRAKER and StringTie models previously

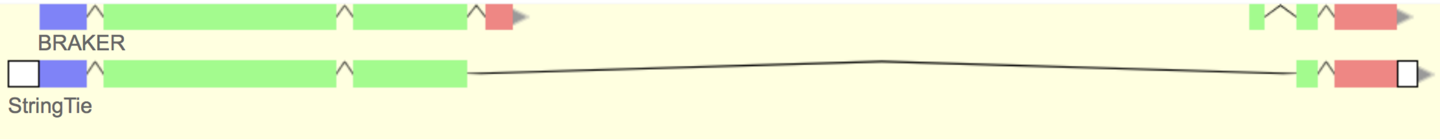


2. Identify the putative split point

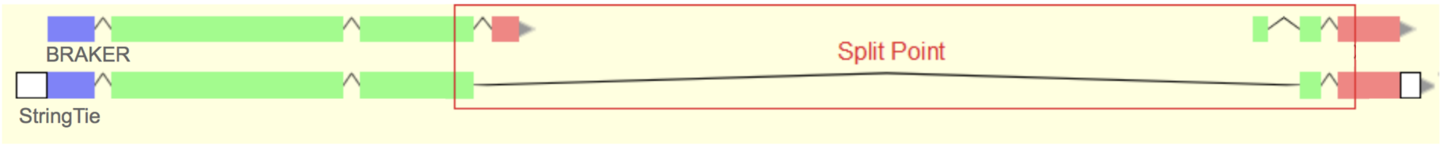


3. If available, add IsoSeq transcripts

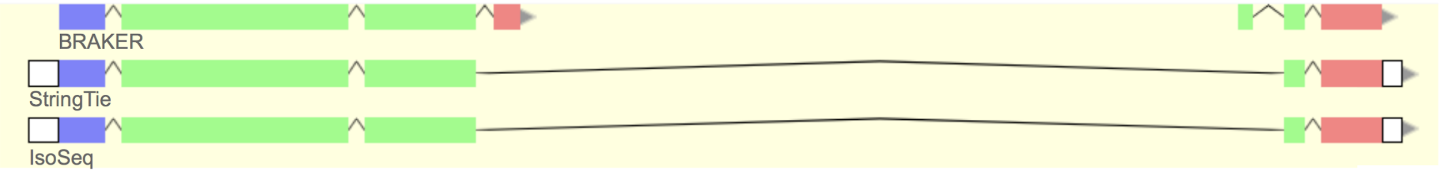

4. If available, examine Illumina reads mapping to the split point, using the genome browser grid as a reference.
   - By examining the mapped reads by eye, you will observe that the large majority (>90%) of the reads will support either the fused or split models.

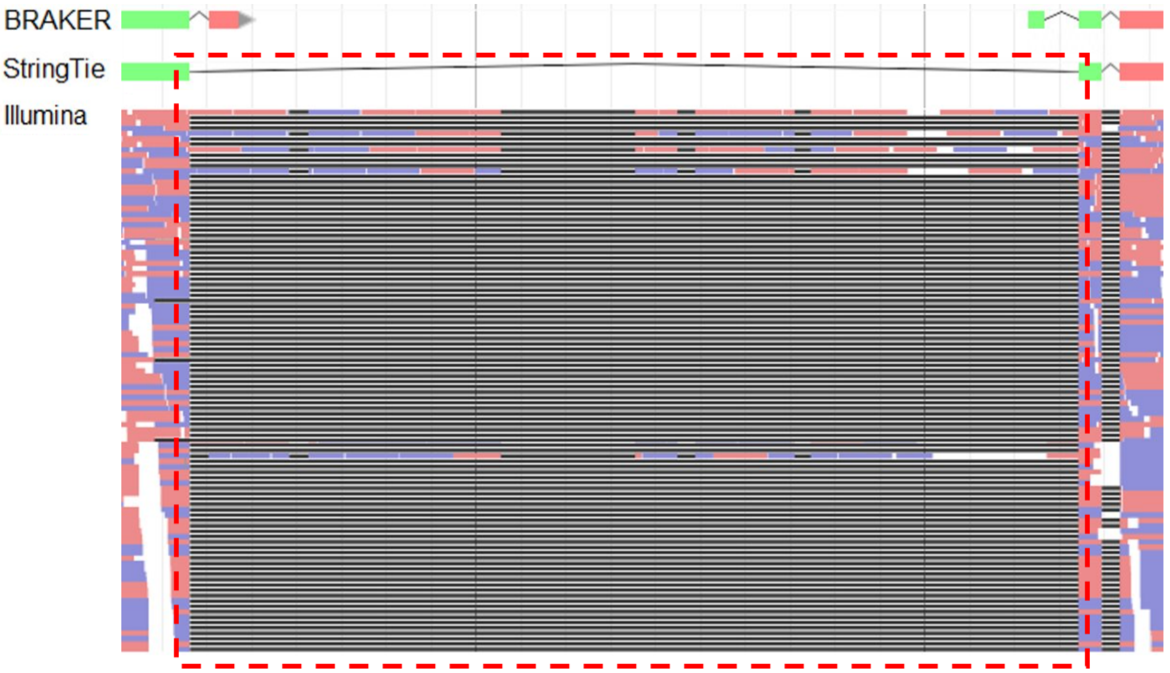


The intron of the StringTie model is strongly supported by Illumina read alignments.

5. Add a few reads from the Illumina track mapping to the split point. Ensure the boundaries covered by the reads mapping to the split point match the supported model.

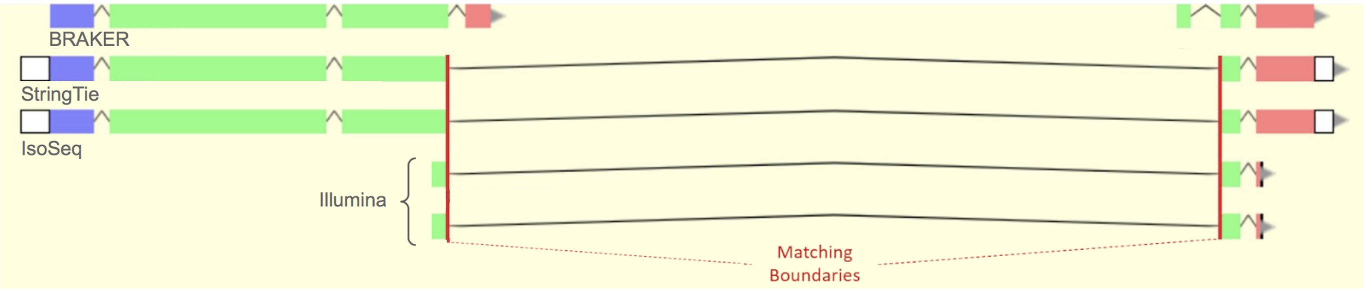


6. Select the model with that is supported by Illumina reads and/or IsoSeq transcripts, remove all others
   - In this case, both IsoSeq and Illumina reads support the StringTie model

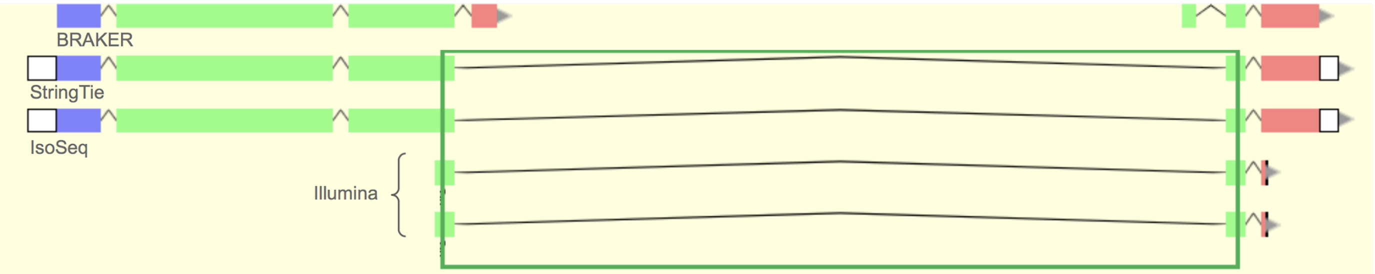


7. Leave the selected model in the User-created Annotations space and remove all other annotations

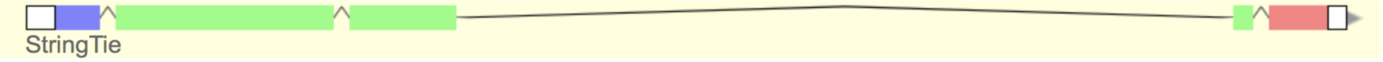


8. Fill curation metadata (Section 2.5.) and move onto the next gene.
   - Metadata code for this case: ‘Gene Split - B:S:IS:IL’

##### 2.4.5. Gene fusion

In an opposing error to gene splits, gene prediction algorithms sometimes fuse two separate genes. Here, you will often observe two adjacent StringTie models that overlap with a single BRAKER model:

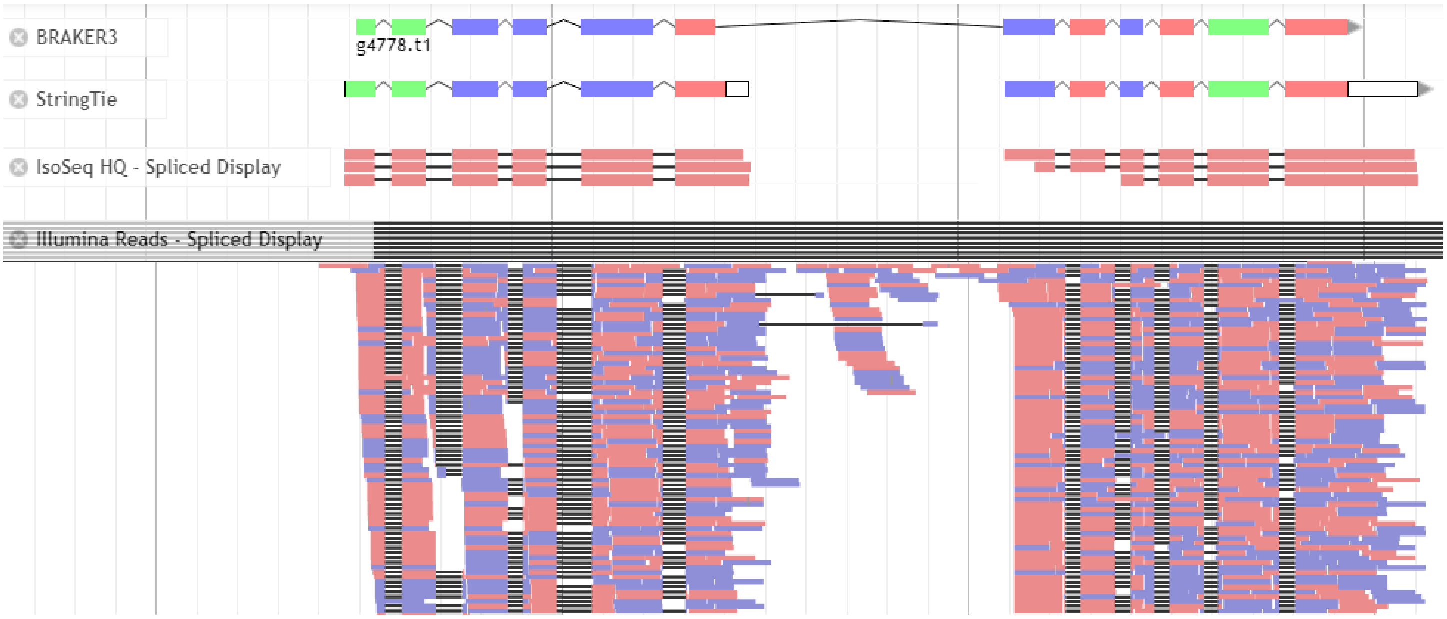


Following the General Workflow:

1. Are there any BRAKER models? (Yes)
2. Add BRAKER model
3. Are there any StringTie models? (Yes)
4. Add StringTie model
5. Does StringTie fully support BRAKER? (No)
6. Are they different only in the UTRs? (No)
7. Are there two adjacent StringTie models that overlap with a single BRAKER model? (Yes)
8. Refer to workflow: “Gene Fusion”

Following the Gene

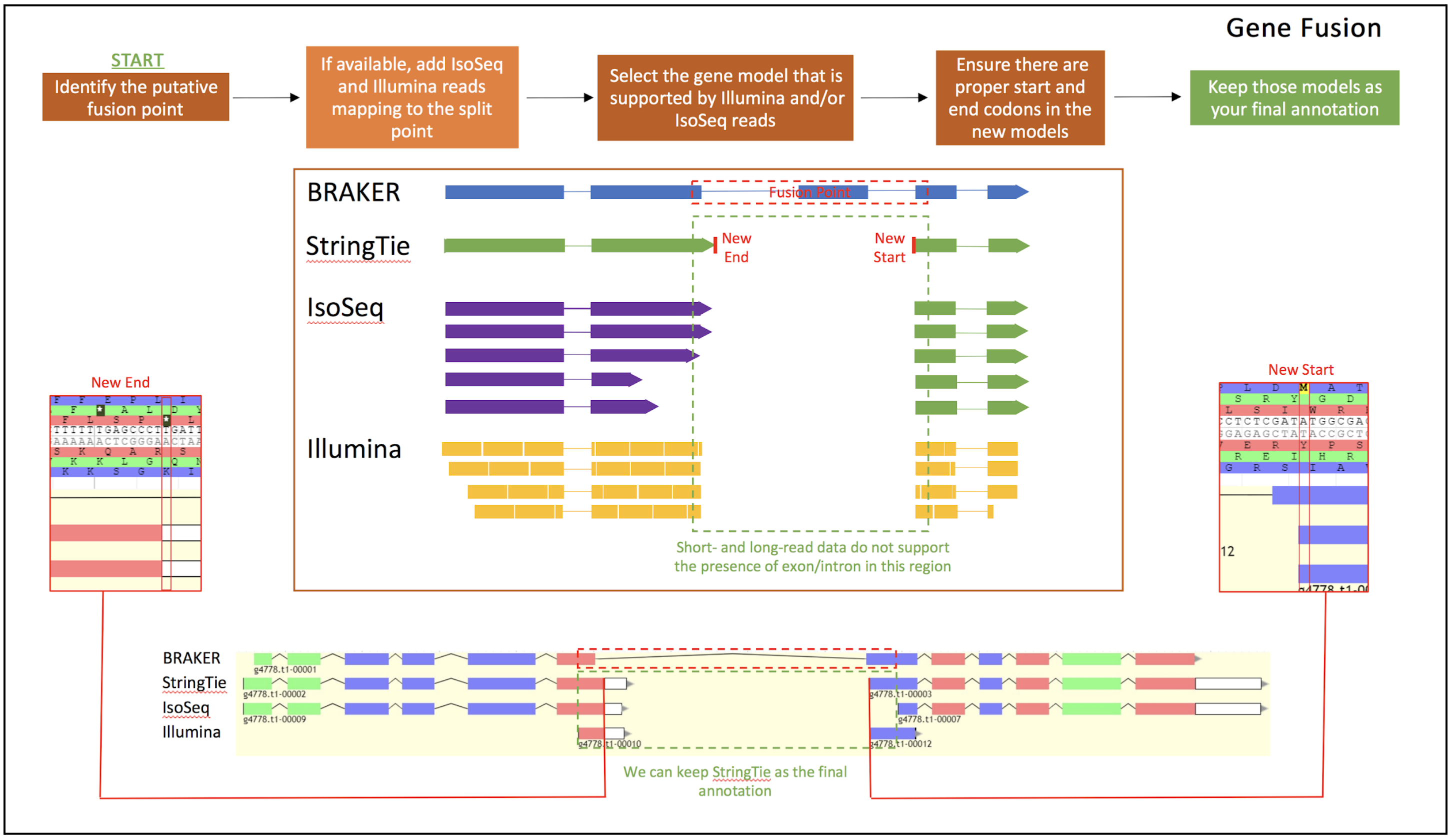


Fusion workflow:

1. We have added BRAKER and StringTie models previously.

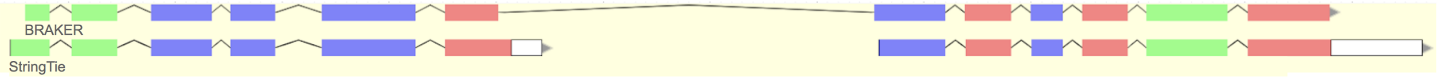


2. Identify the putative fusion point

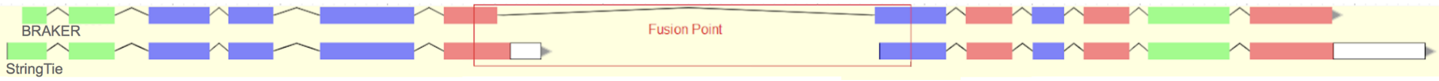


3. If available, add IsoSeq transcripts

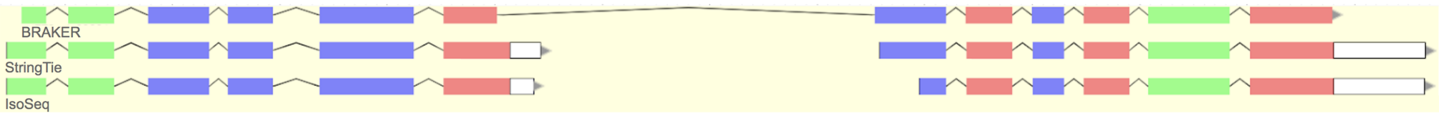

4. If available, examine Illumina reads mapping to the split point, using the genome browser grid as a reference.
    - By examining the mapped reads by eye, you will observe that the large majority (>90%) of the reads will support either the fused or split models.

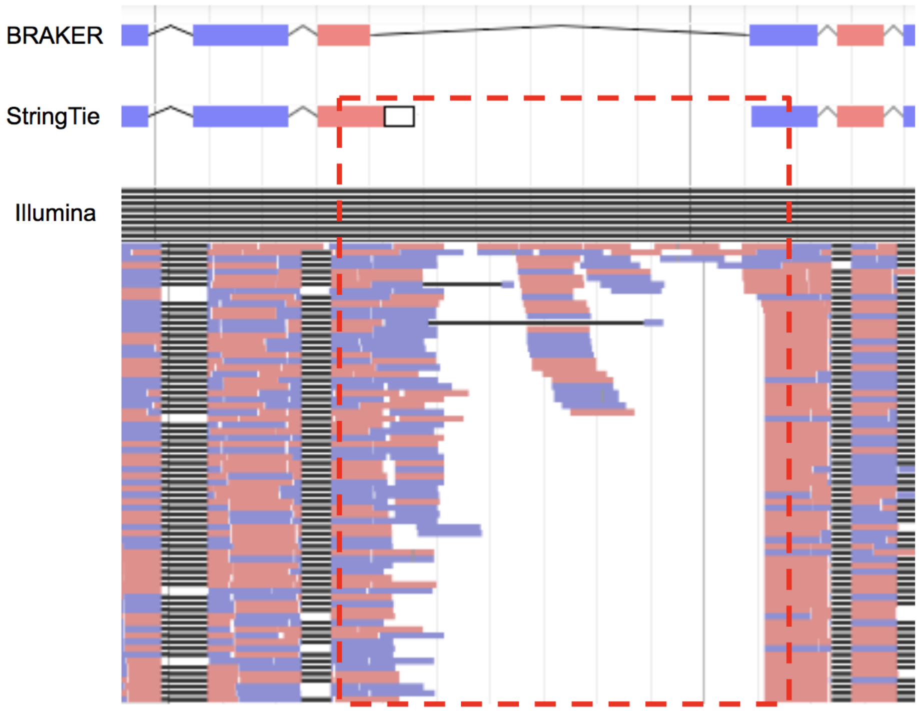


The intron of the BRAKER model is not supported by Illumina read alignments.

5. Select the model that is supported by IsoSeq transcripts and/or Illumina reads, remove all others
    - In this case, IsoSeq and Illumina reads do not support the presence of the connecting intron in the BRAKER model.

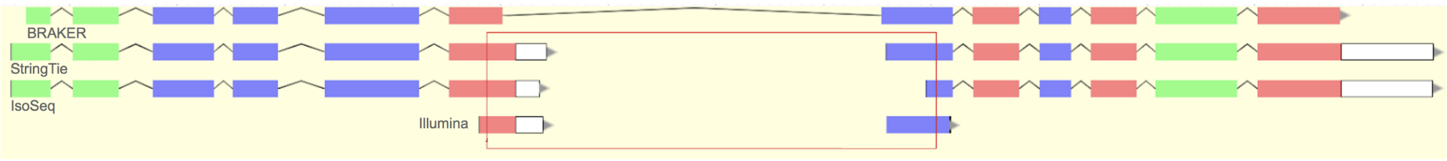


6. Ensure that you can find proper start and stop codons in the newly selected model
    - 3’ terminal exons should end on a stop codon (asterisk)
    - 5’ terminal exons should start on a start codon (M)

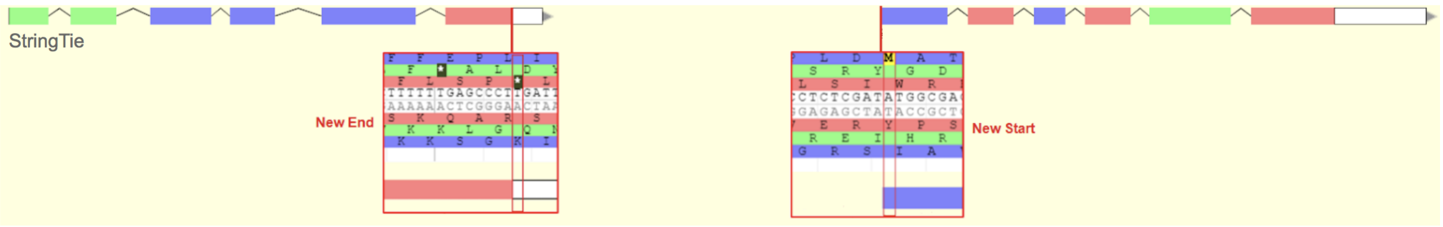

7. If a start/stop codon is not present, you can set the closest in-frame start/stop codon by right-clicking on a terminal exon and selecting ‘Set Translation Start’ or ‘Set Translation End’.

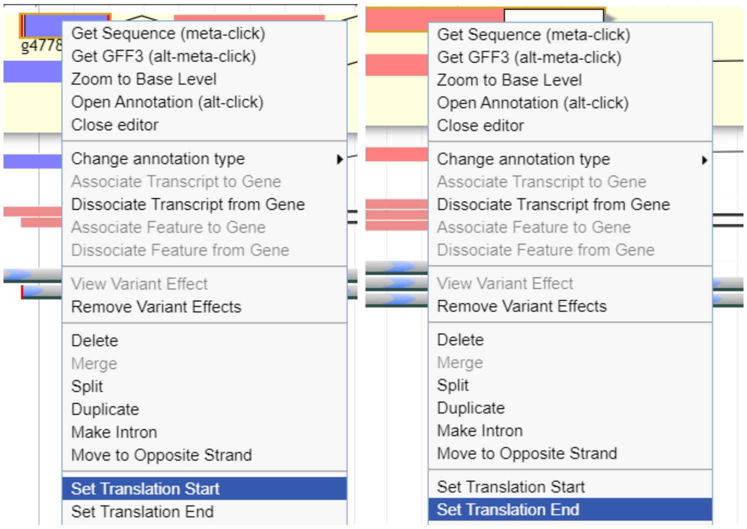


- Alternatively, you can select the entire annotation and select ‘Set Longest Orf’ to automatically reset the entire reading frame of the annotation after you have made edits to its structure.

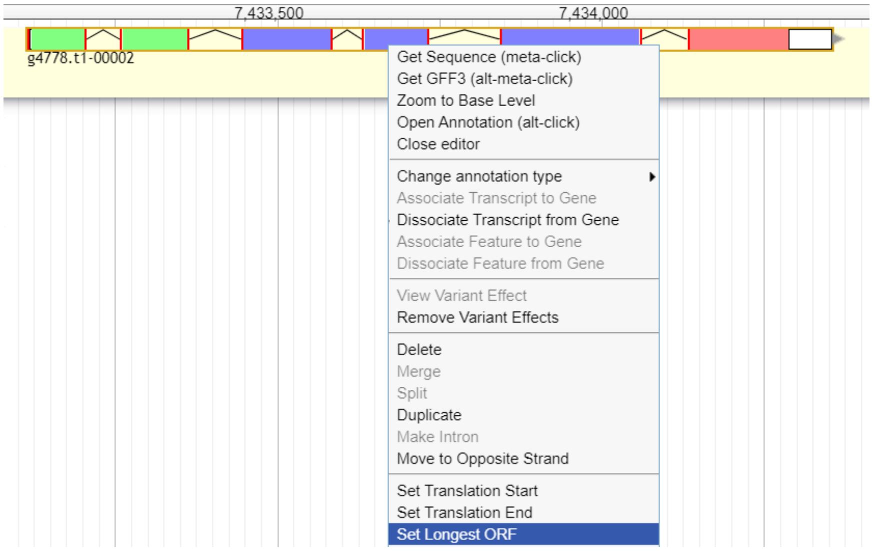


8. Leave the selected model(s) in the User-created Annotations space.

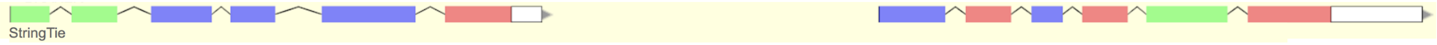


9. Fill curation metadata (Section 2.5.), and move onto the next gene.
   - Metadata code for this case: ‘Gene Fusion - B:S:IS:IL’

##### 2.4.6. Multiple isoforms

Oftentimes a single gene can be transcribed into multiple transcripts that have differences in exon length and/or number. We refer to these alternative transcripts as isoforms. To break down the Multiple Isoforms workflow, we will review a simple and complex case for this type of annotation

###### 2.4.6.1. Simple multiple isoforms case

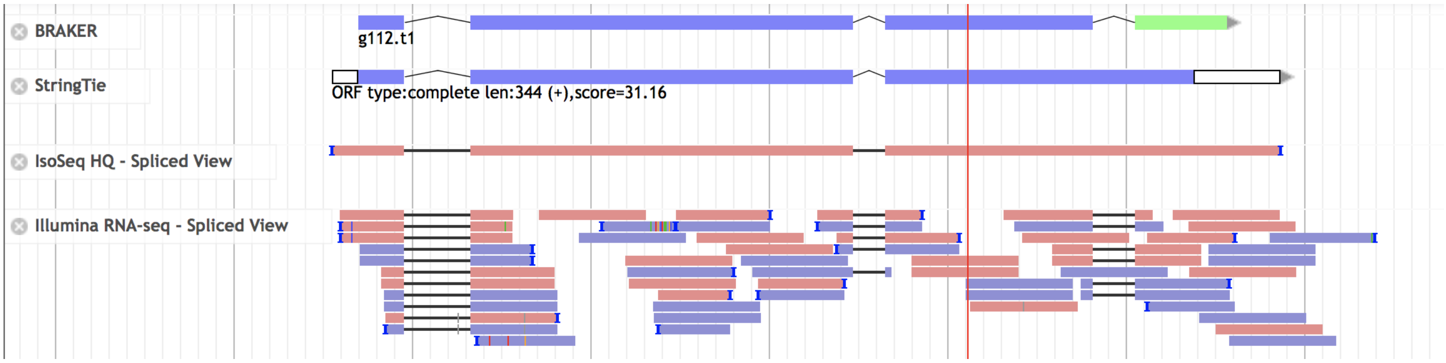


Following the General Workflow:

1. Are there any BRAKER models? (Yes)
2. Add BRAKER
3. Are there any StringTie models? (Yes)
4. Add StringTie
5. Does StringTie fully support BRAKER? (No)
6. Are they different only in the UTRs? (No)
7. Are there two adjacent StringTie models that overlap with a single BRAKER model? (No)
8. Are there two adjacent BRAKER models that overlap with a single StringTie model? (No)
9. Are there multiple overlapping gene models with differences in the number or length of their exons? (Yes)
10. Refer to workflow: ‘Multiple isoforms’

Following the Multiple Isoforms workflow:

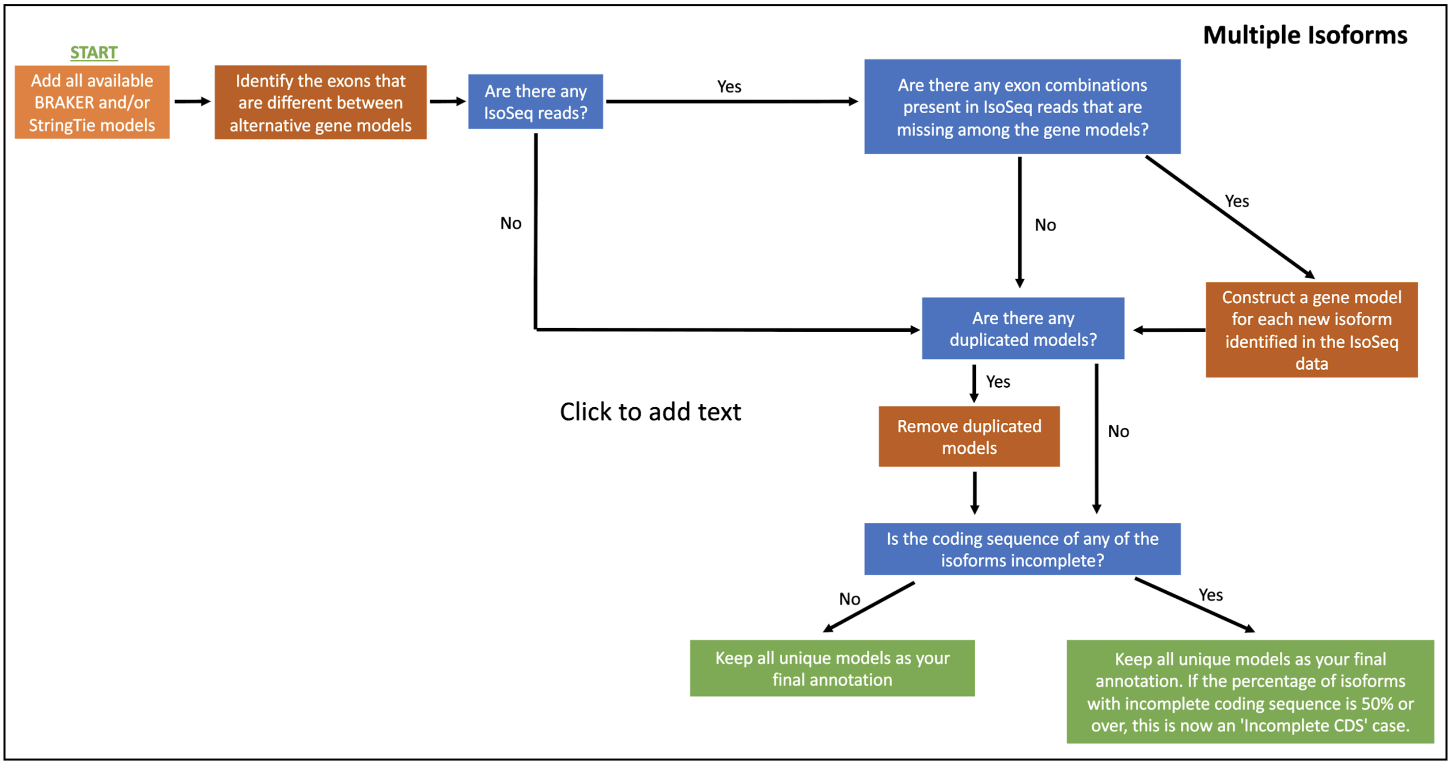


1. We have added BRAKER and StringTie models previously.

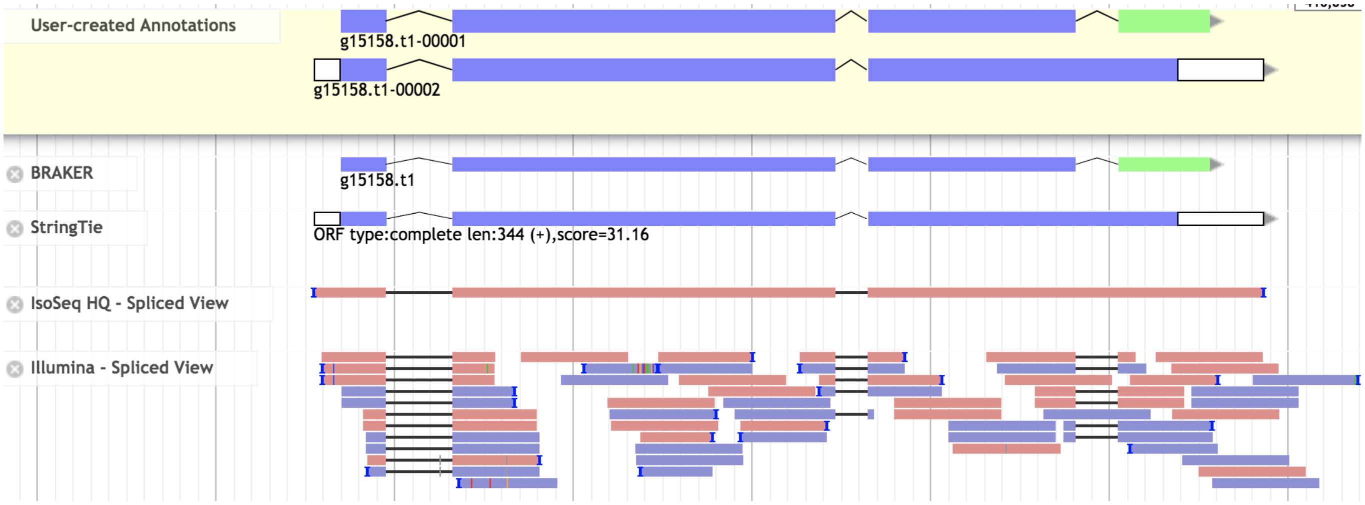


2. Identify the exons that are different between alternative gene models

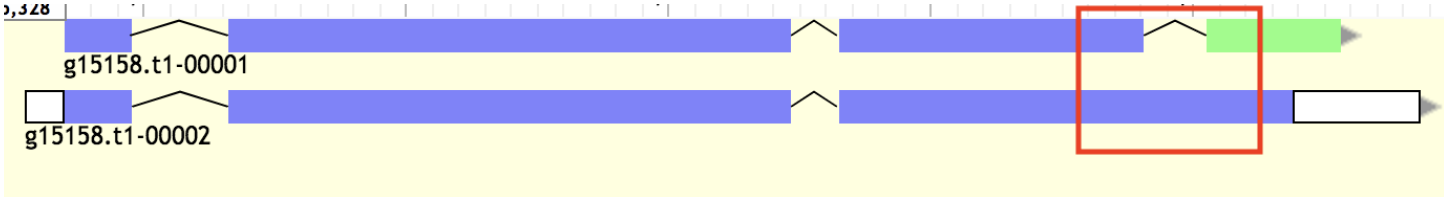


3. Are there any IsoSeq reads? (Yes)

4. Are there any exon combinations present in IsoSeq reads that are missing among the gene models? (No)

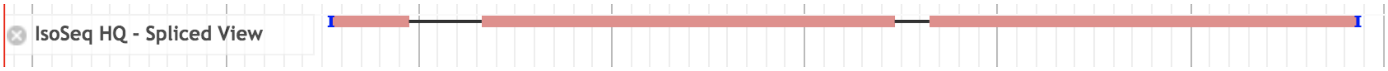


5. Are there any duplicated models? (No)

6. Is the coding sequence of any of the isoforms incomplete? (No)
   - If there were any incomplete isoforms and the percentage of incomplete isoforms exceeds 50%, this annotation would become an 'Incomplete CDS' case.

7. Keep all unique models as your final annotation.
   - Extend the UTRs based on Illumina and IsoSeq when possible.

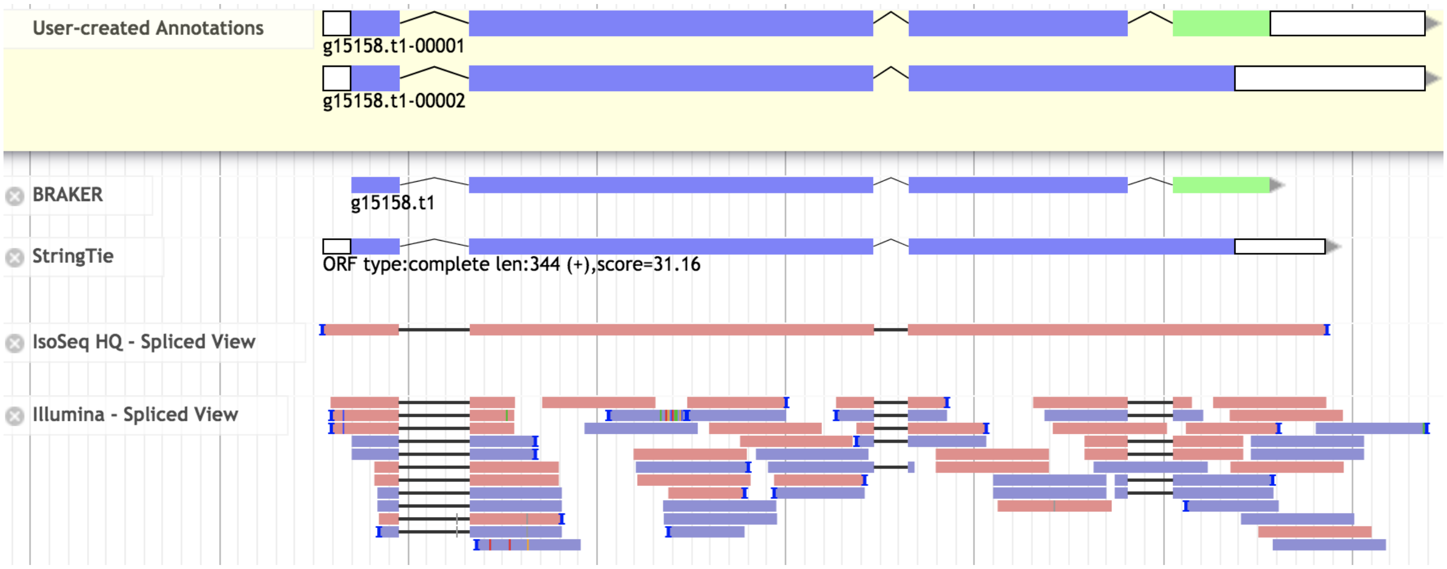


8. Fill curation metadata (Section 2.5.), and move onto the next gene.
   - Metadata code for this case: 'Multiple Isoforms - B:S:IL'

##### 2.4.6.2. Complex multiple isoforms case

The general workflow steps taken for this complex case are the same as the simple case.

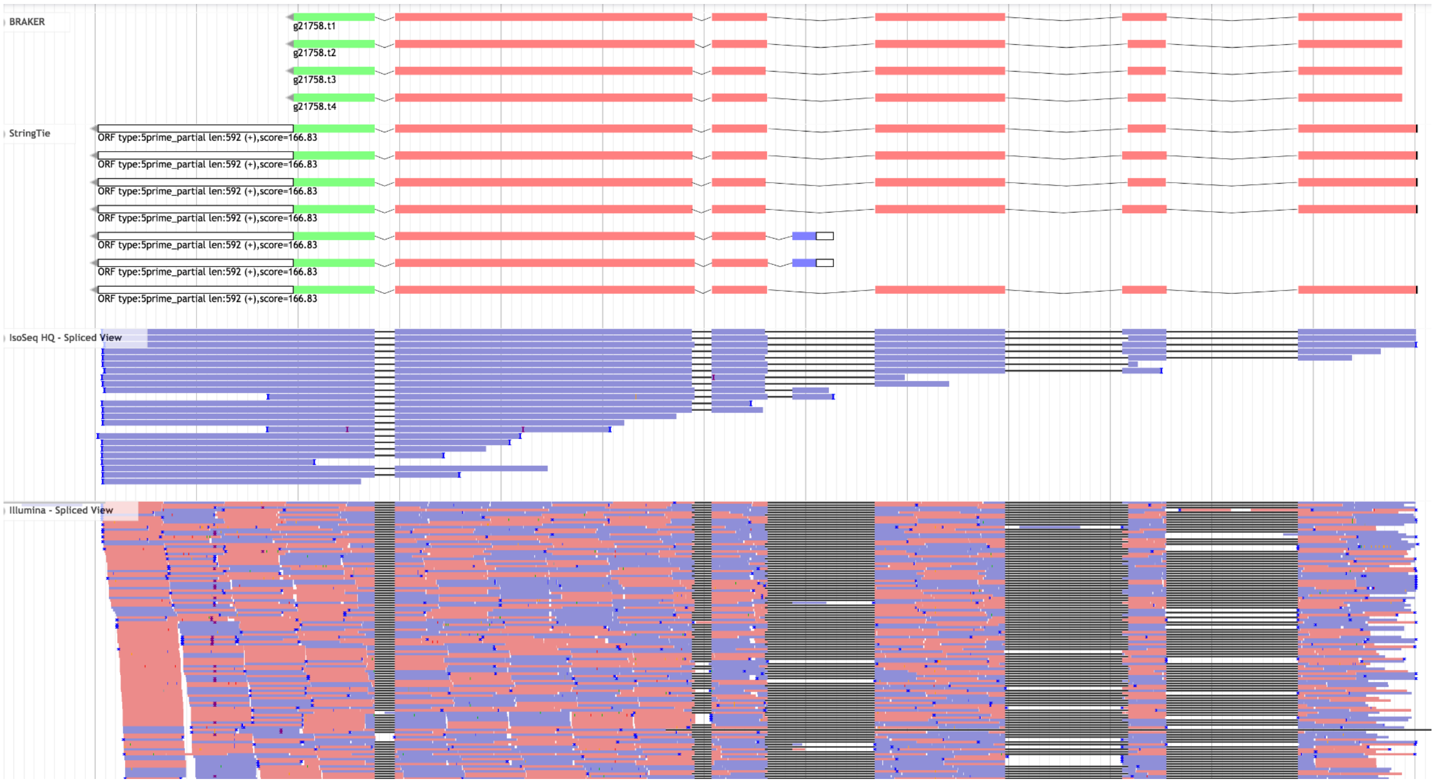


Following the Multiple Isoforms workflow:

1. We have added BRAKER and StringTie models previously.

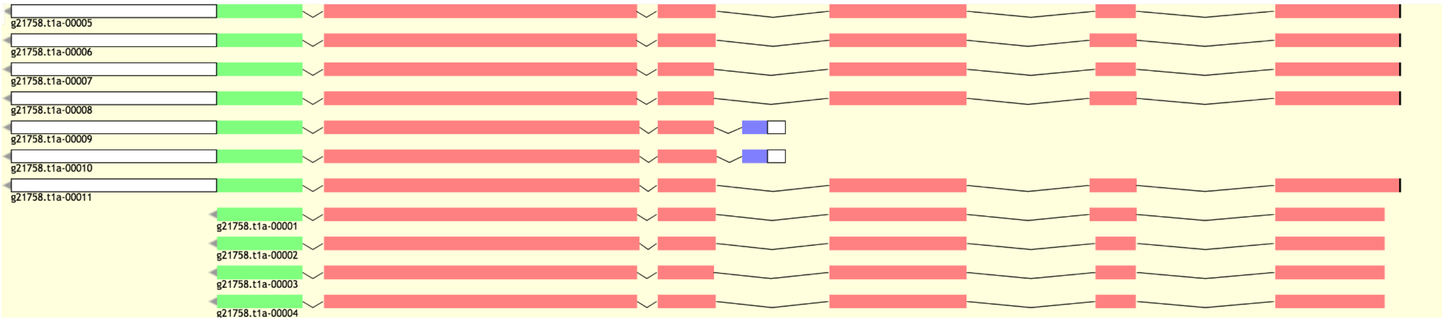

2. Identify the exons that are different between alternative gene models
    ○ The StringTie models often have errors in the prediction of the translation start (blue). This is not considered a real difference between isoforms, and must be repaired before looking for new or duplicated isoforms (Set translation start or Set optimal ORF).

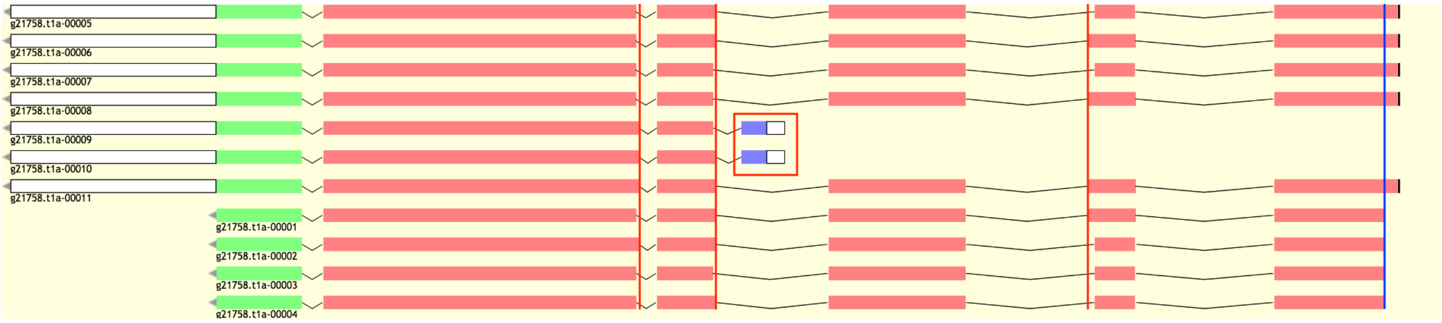


    ○ There are four differences between the models above:
        i. Variable length of Exon 2 (S and L)
        ii. Variable length of Exon 4 (S and L)
        iii. Variable length of Exon 5 (S and L)
        iv. An alternative translation start between Exon 3 and Exon 4 (ATS)

3. Are there any IsoSeq reads? (Yes)

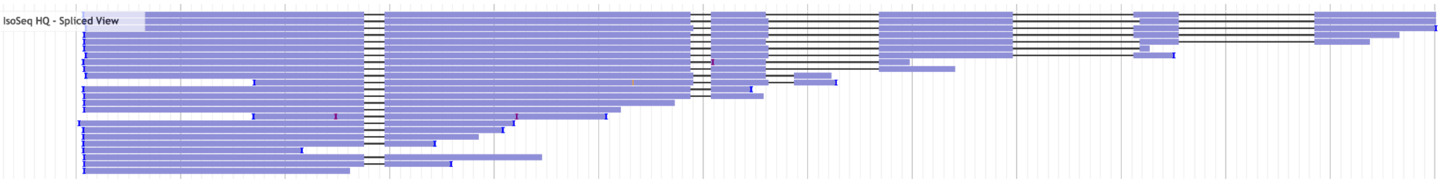


4. Are there any exon combinations present in IsoSeq reads that are missing among the gene models? (No)
    ○ In order to track which isoforms we have present in our current models, we can give each isoform an ID based on the characteristics of the exons that vary across models.
        i. For example a model that has short Exons 2, 4, and 5 can be called 'S:S:S'

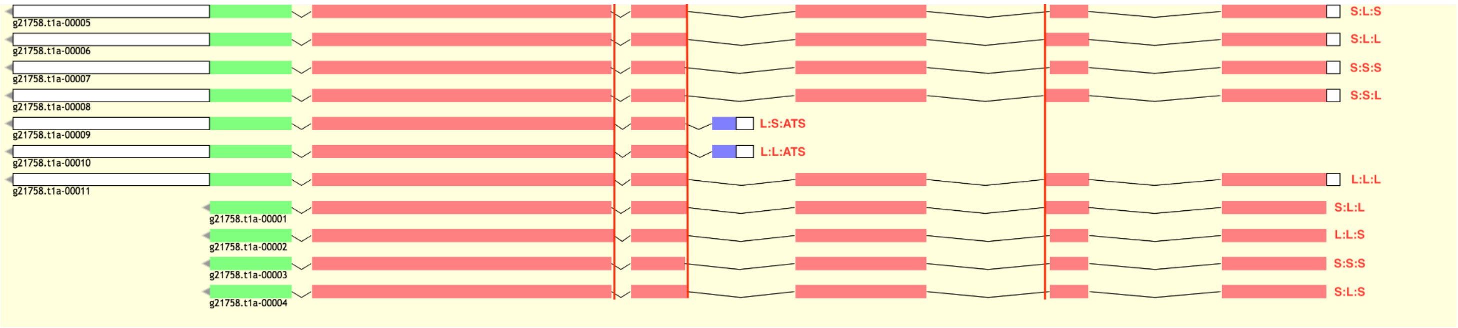

- We can apply the same label system to IsoSeq reads.

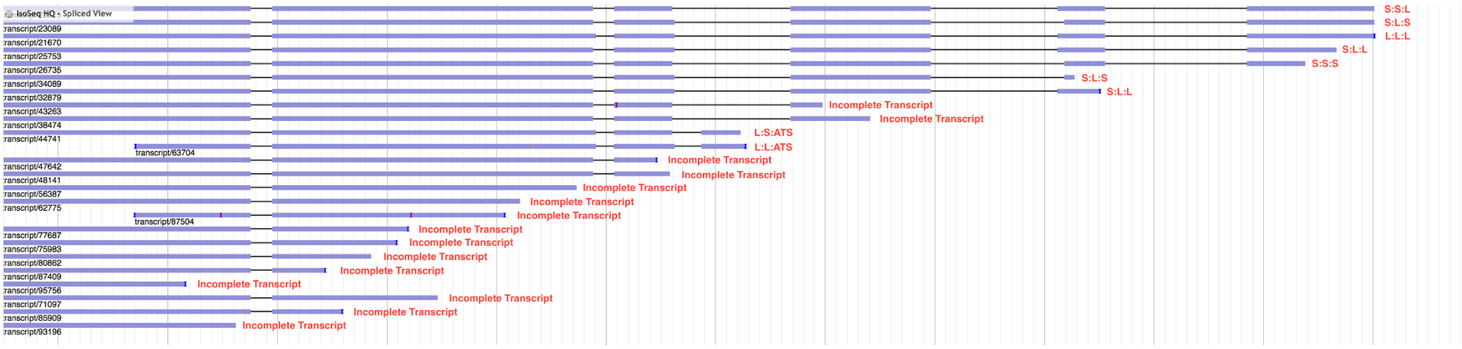


- If we had any missing isoforms, we can construct these missing models by:
    i. Duplicate a gene model that is similar in structure to a reference IsoSeq read.
    ii. Add/Remove exons that are present/missing in the IsoSeq reference
    iii. Extend/Shorten any exons that have different length in the IsoSeq reference

5. Are there any duplicated models? (Yes)
    - There are two redundant copies of S:L:L, S:L:S, and S:S:S. We must remove these.

6. Is the coding sequence of any of the isoforms incomplete? (No)
    - If the percentage of incomplete isoforms were to be 50% or over, the metadata code for this case would be: ‘Incomplete CDS - B:S’

7. Keep all unique models as your final annotation.
    - Note that the UTRs of BRAKER models were extended as well.

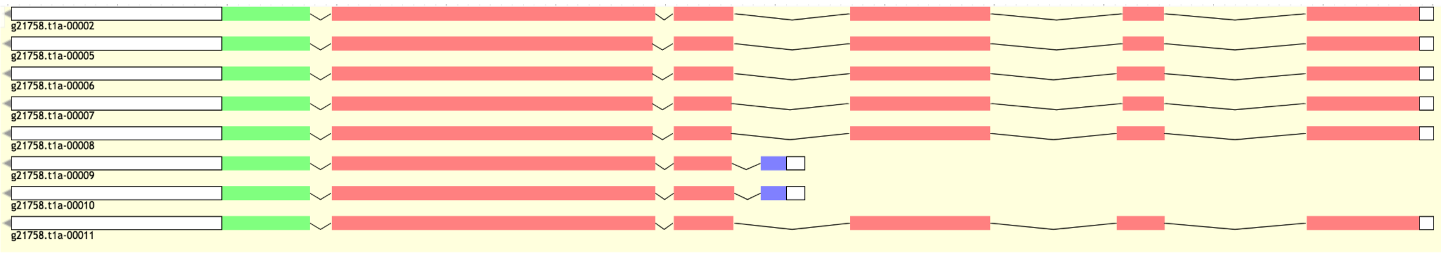


8. Fill curation metadata (Section 2.5.), and move onto the next gene.
    - Metadata code for this case: ‘Multiple Isoforms - B:S’

##### 2.4.7. Terminal Exon Repair

###### 2.4.7.1. Missing Exon

In some cases, we may observe Illumina evidence for an exon that is missing from the predicted gene models. We must repair the gene terminus by adding the missing exon.

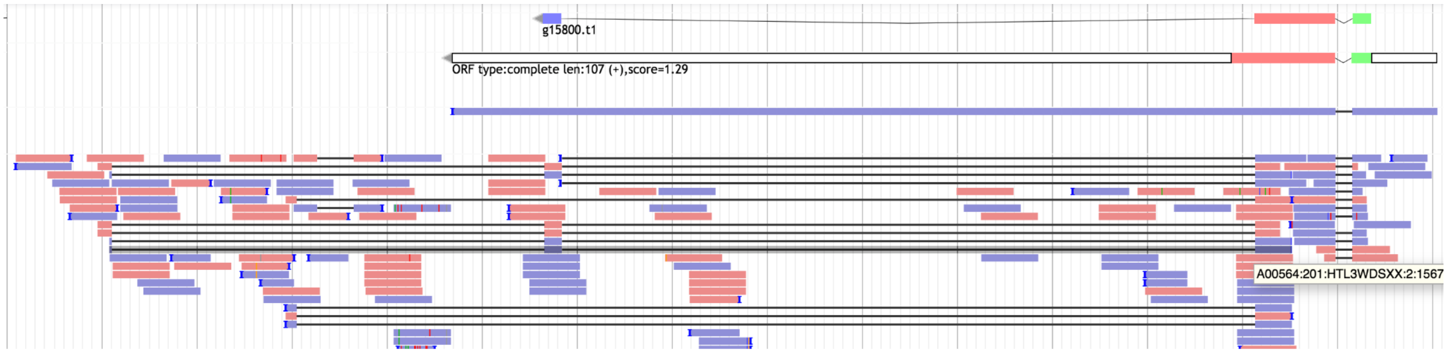


Following the General Workflow:

1. Are there any BRAKER models? (Yes)
2. Add BRAKER
3. Are there any StringTie models? (Yes)
4. Add StringTie
5. Does StringTie fully support BRAKER? (No)
6. Are they different only in the UTRs? (No)
7. Are there two adjacent StringTie models that overlap with a single BRAKER model? (No)
8. Are there two adjacent BRAKER models that overlap with a single StringTie model? (No)
9. Are there multiple overlapping gene models with differences in the number/length of their exons? (No)
10. Does BRAKER/StringTie have a missing exon that is supported by Illumina? (Yes)
11. Refer to workflow: TER - Missing exon'

Following the TER - Missing Exon workflow:

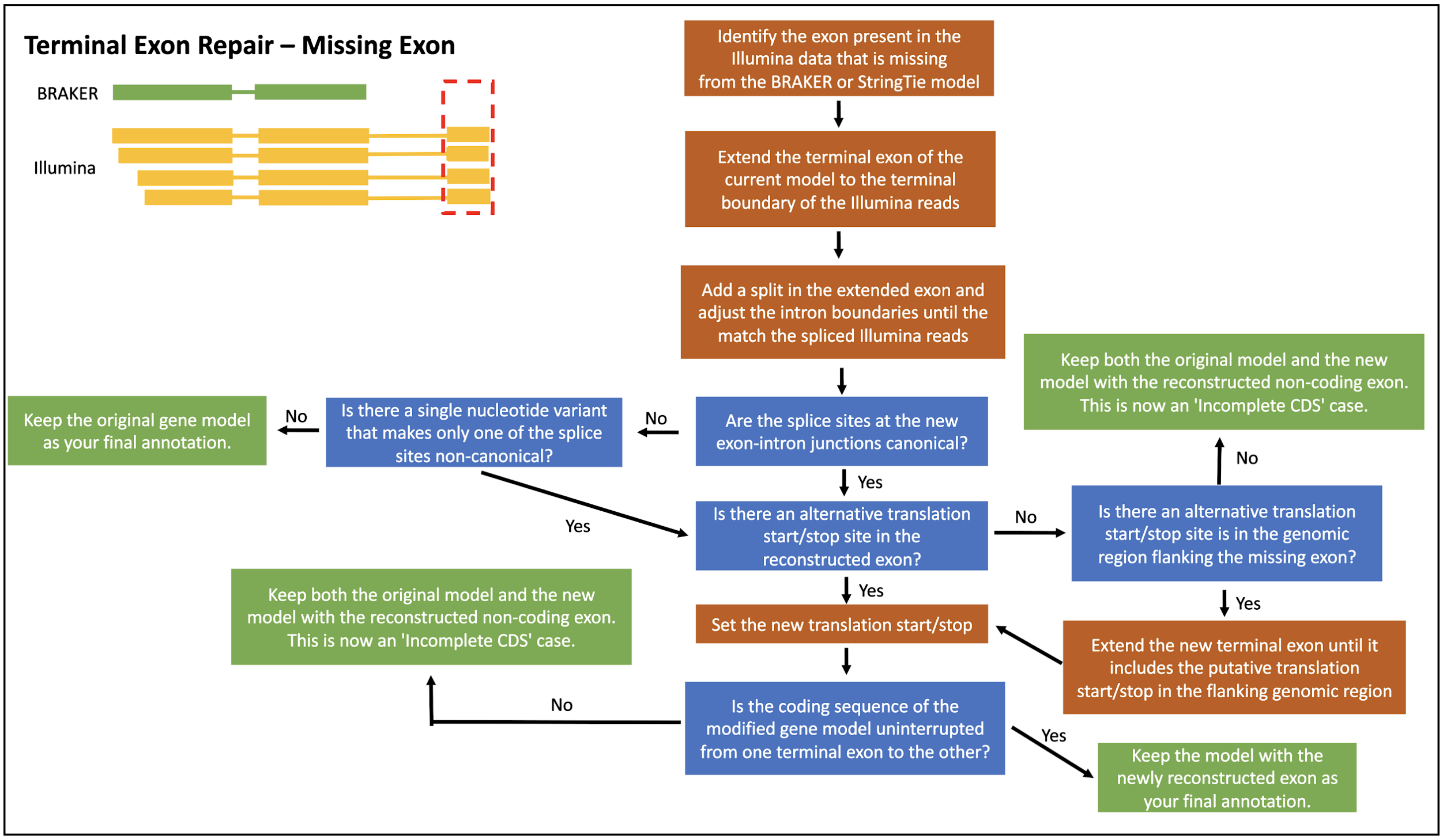


12. We have added BRAKER and StringTie models previously.

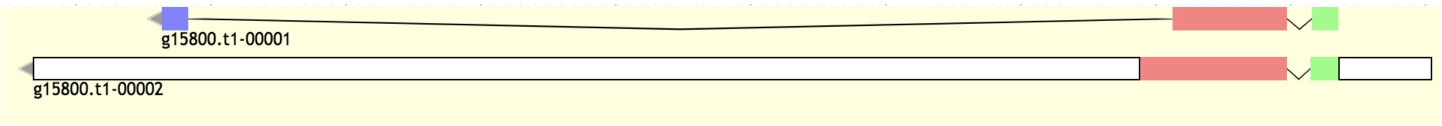


13. Identify the exon present in the Illumina data and missing from BRAKER/StringTie

a. We can observe there are two alternative isoforms that have a missing exon.

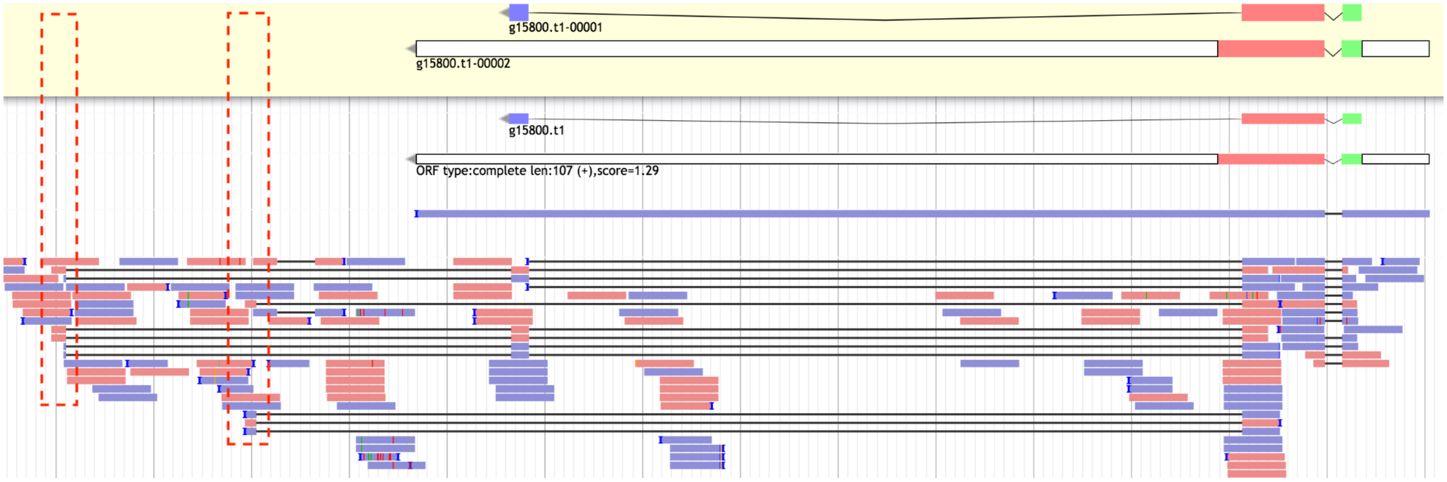

14. Extend the terminal exon of the current model to the terminal boundary of the missing exon in the Illumina data.

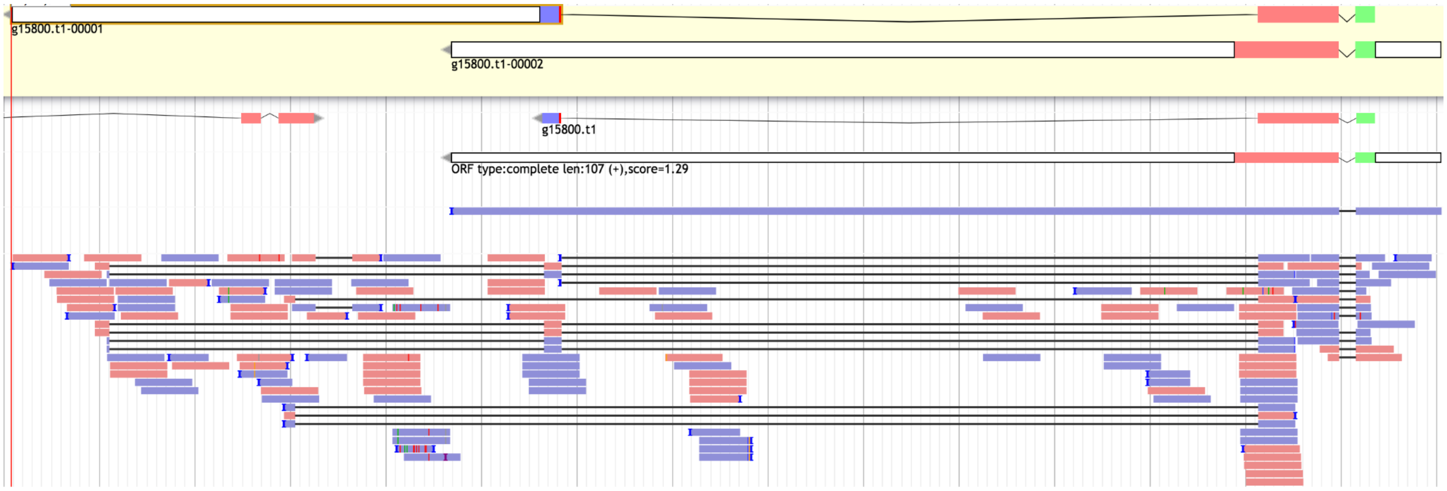


15. Add a split in the extended exon.

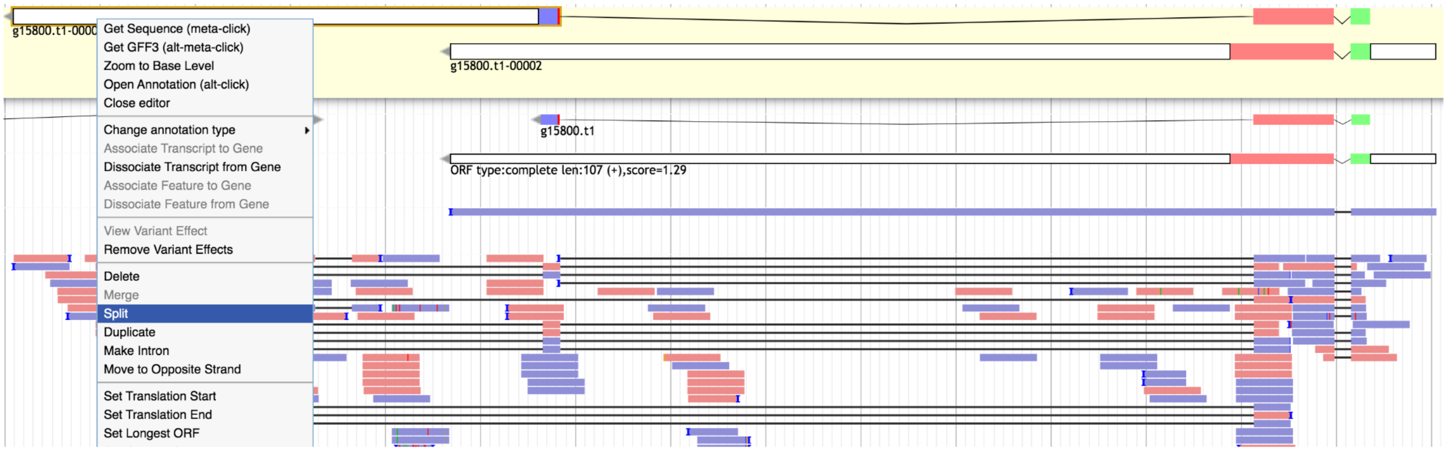


16. Adjust the intron boundaries so that they match the spliced Illumina reads.

<u>3’ boundary</u>

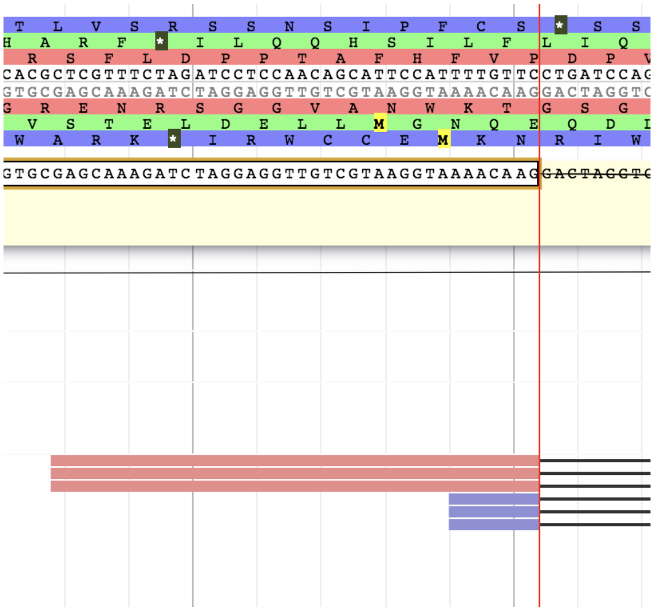

<u>5’ boundary</u>

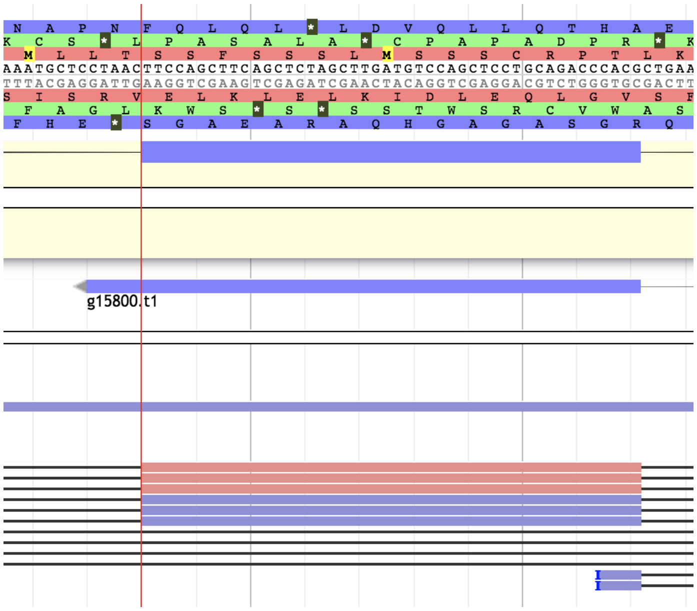

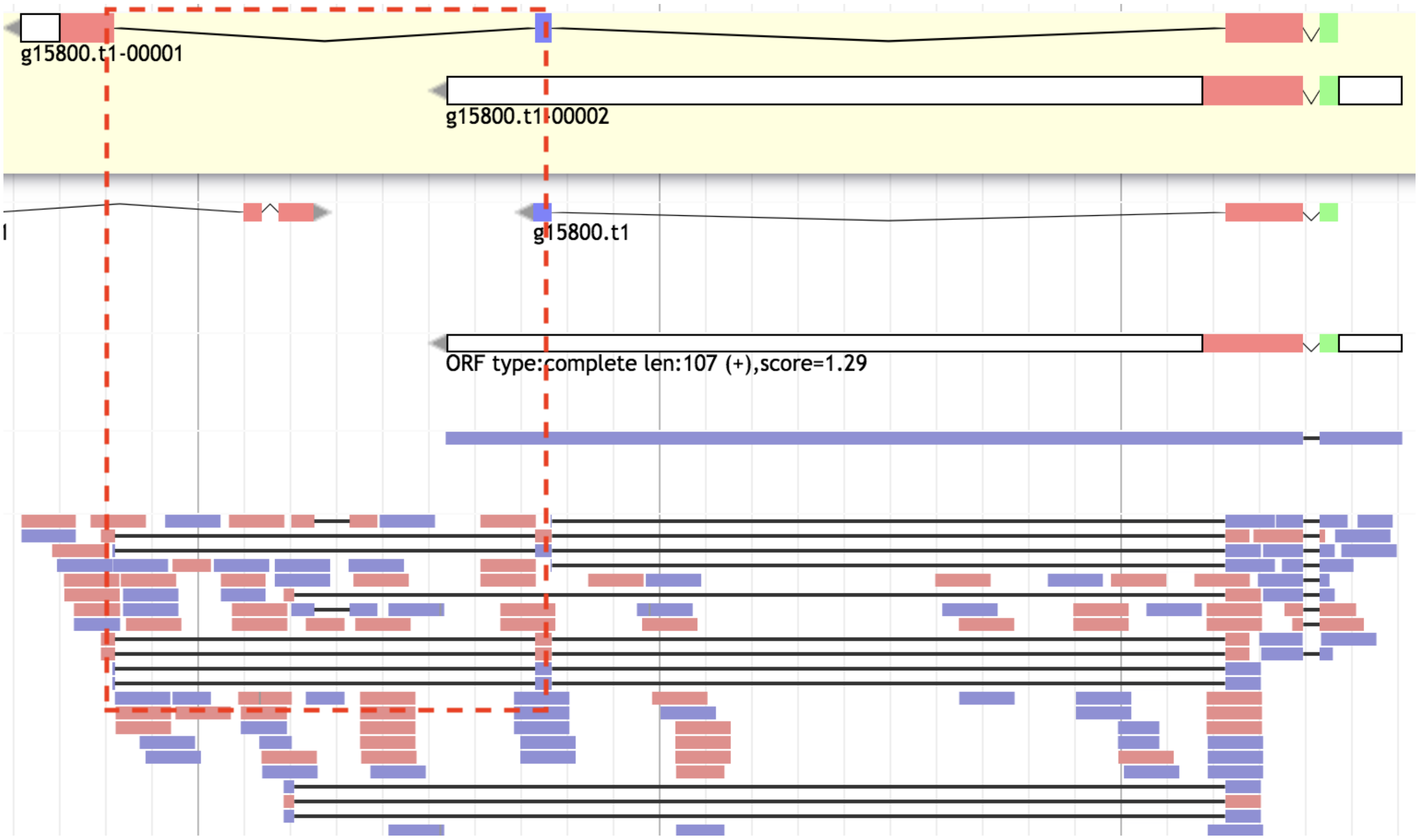


a. We can repeat the same process for the other potential terminal exon (and extend UTRs!).

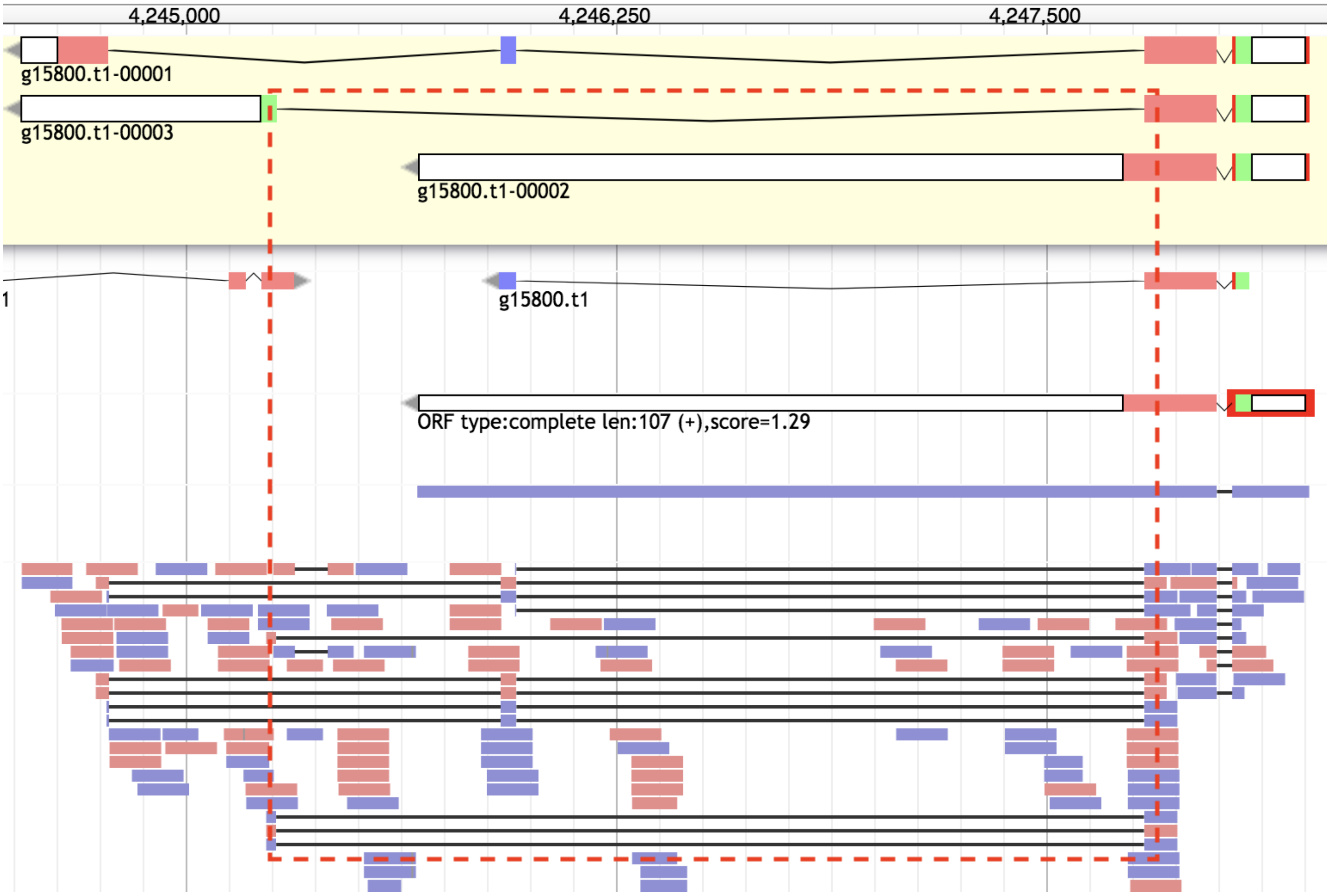

17. Are the splice sites at the new intron/exon junctions canonical? (Yes)
    a. Discard any low coverage gene models where the splice sites are non-canonical on both sides of the intron.

18. Set the new translation start/stop
    a. In this case, it was automatically set by Apollo.

19. Is the coding sequence of the modified gene model uninterrupted for one terminal exon to the other? (Yes)
    a. If adding the missing exon were to generate an incomplete model, we would keep both the modified and original model, and annotate the metadata code: 'Incomplete CDS - B:IL'

20. Keep the modified gene model(s) as your final annotation.

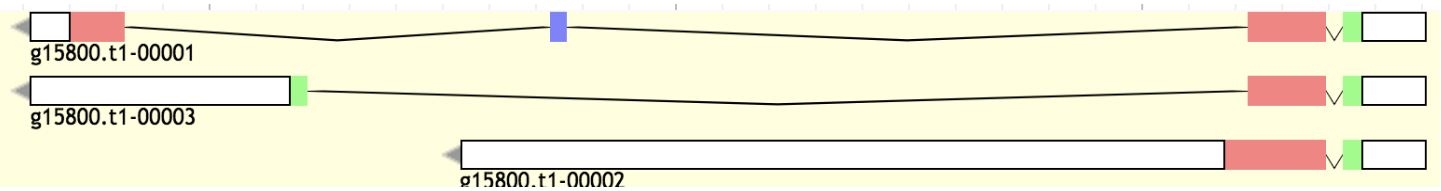


21. Fill curation metadata (Section 2.5.), and move onto the next gene.
    a. Metadata code for this case: 'Multiple Isoforms - B:S:IL'
    b. If this gene annotation consisted of a single terminal exon repair, the metadata would be 'Terminal Exon Repair - B:IL'. Since we have multiple isoforms, that takes priority.

##### 2.4.7.2. Additional Exon

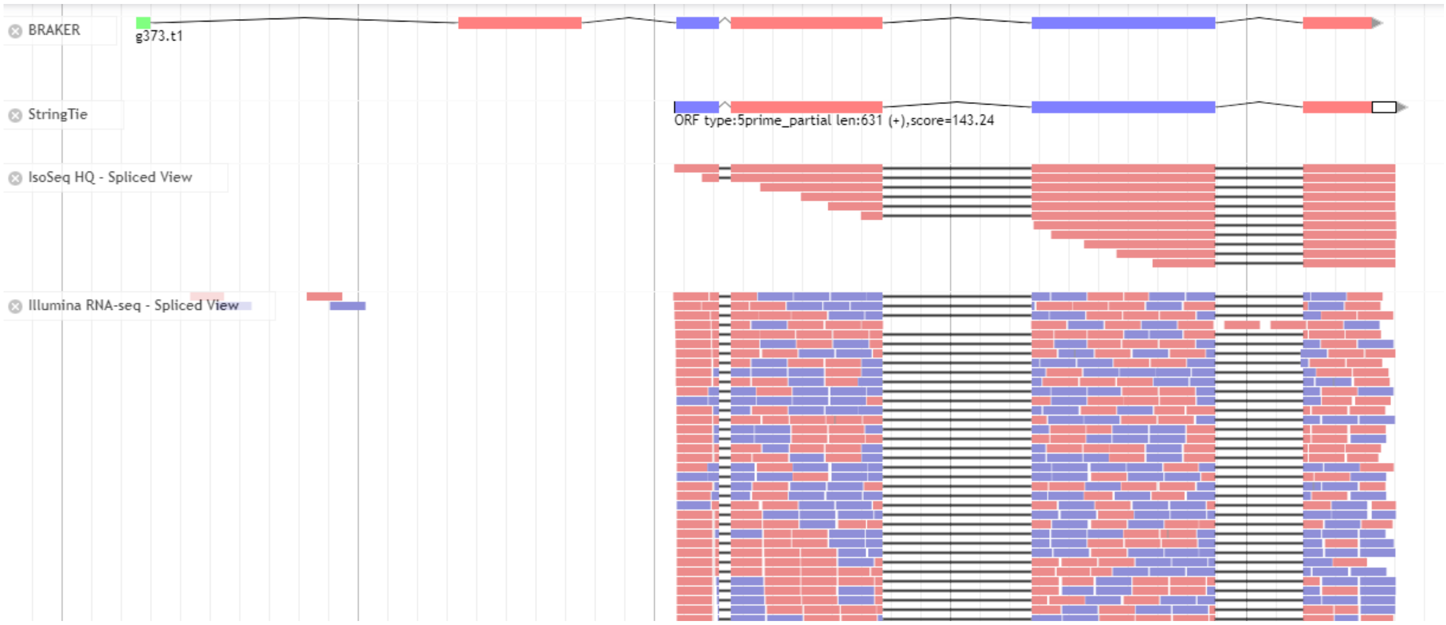


Following the General Workflow:

1. Are there any BRAKER models? (Yes)
2. Add BRAKER
3. Are there any StringTie models? (Yes)
4. Add StringTie
5. Does StringTie fully support BRAKER? (No)
6. Are they different only in the UTRs? (No)
7. Are there two adjacent StringTie models that overlap with a single BRAKER model? (No)
8. Are there two adjacent BRAKER models that overlap with a single StringTie model? (No)
9. Are there multiple overlapping gene models with differences in the number/length of their exons? (No)
10. Does BRAKER/StringTie have an additional exon that is not supported by Illumina? (Yes)
11. Refer to workflow: TER - Additional exon’

Following the TER - Additional Exon workflow:

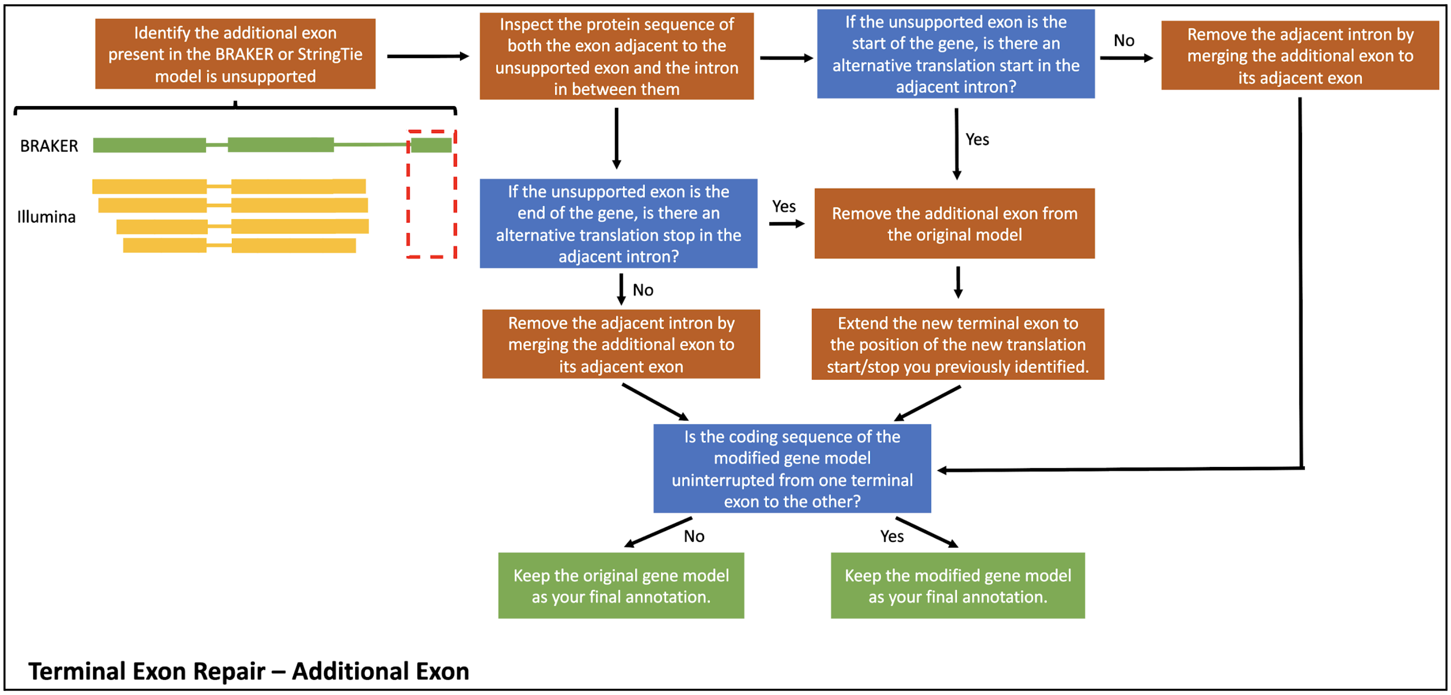


1. We have added BRAKER and StringTie models previously.

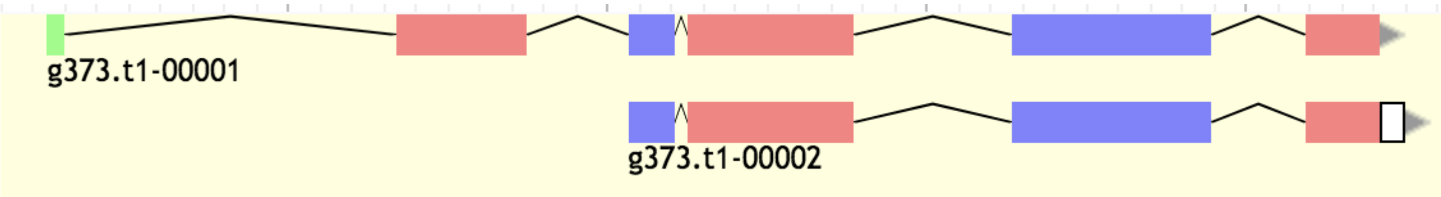


2. Identify the additional exon present in BRAKER/StringTie that is unsupported
   a. In this case, there is more than one exon that is unsupported.

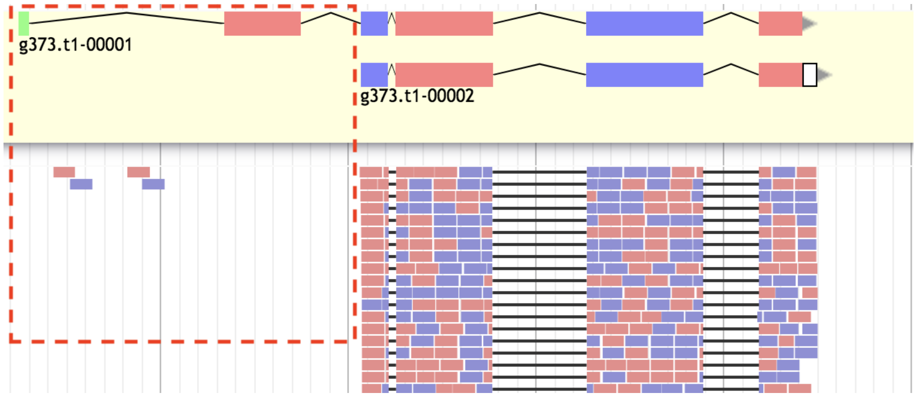

3. Inspect the protein sequence of the exon/intron that is adjacent to the unsupported exon
    a. Since there are two unsupported exons, we will focus on the first adjacent exon that is supported by Illumina.

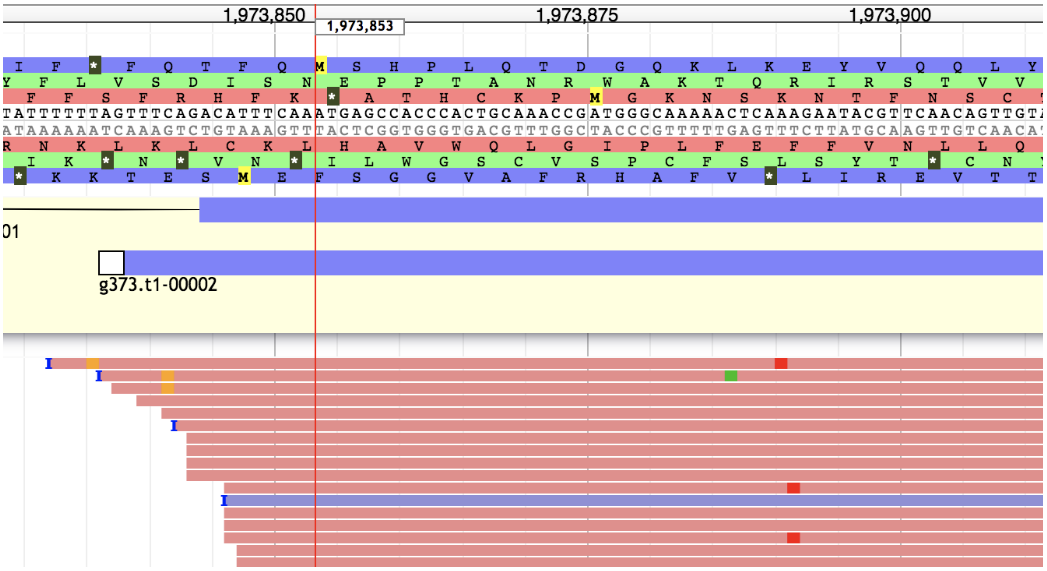


4. In this case, the unsupported exon is in the start of the gene. Is there an alternative start in the exon/intron adjacent to the unsupported exon? (Yes)

5. Remove the additional exon
    a. In this case we remove both unsupported exons

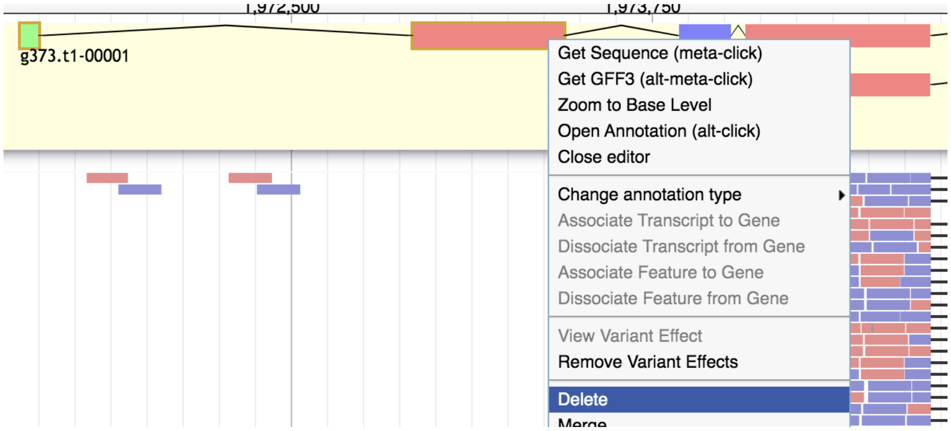


6. Extend the new terminal exon to the position of the alternative start site you previously identified.
    a. In this case, the alternative start site is already covered by the new terminal exon, so we just manually set the new translation start.

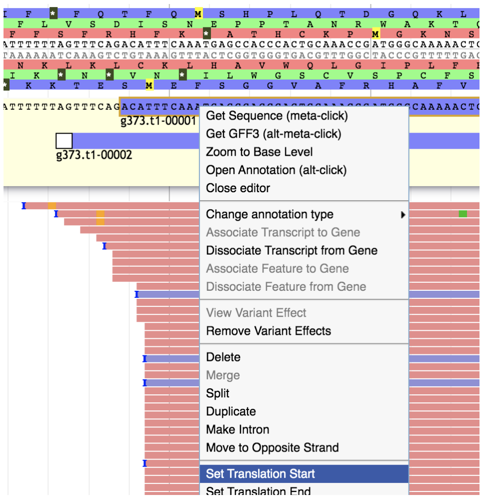

7. The modified gene model has an uninterrupted coding sequence from one terminal exon to the other.
    a. The StringTie model now becomes redundant, so we extend the UTRs of BRAKER and remove StringTie.

8. Keep the modified gene model as your final annotation.

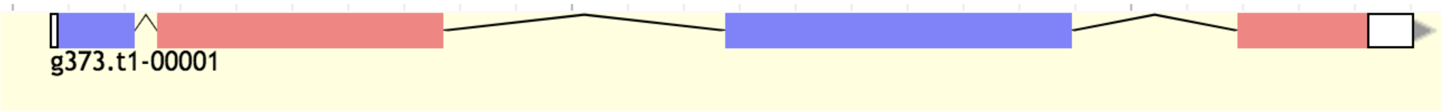


9. Fill curation metadata (Section 2.5.), and move onto the next gene.
    a. Metadata code for this case: 'Terminal Exon Repair - B:S'

##### 2.4.8. Incomplete CDS

Gene models with incomplete coding sequence may arise when resolving multiple isoforms or terminal exon repairs. An incomplete CDS is simply an outcome of other curation workflows (see example below), so there isn't a particular workflow to follow. However, we must add metadata descriptions when appropriate. Incomplete CDS takes priority over all other metadata descriptions when the percentage of incomplete isoforms is 50% or over. For example, if you perform a terminal exon repair and the resulting model has an incomplete CDS, you will keep both the incomplete model and the original model. In this case exactly 50% of the isoforms are incomplete, making that case an 'Incomplete CDS' case. The same concept applies when annotating multiple isoforms.

##### 2.4.9. New Gene

In some cases, you will encounter a large cluster of Illumina reads that resemble the structure of a gene, but lack any gene models. You can attempt to assemble a gene model from scratch following these instructions:

Following the New Gene workflow:

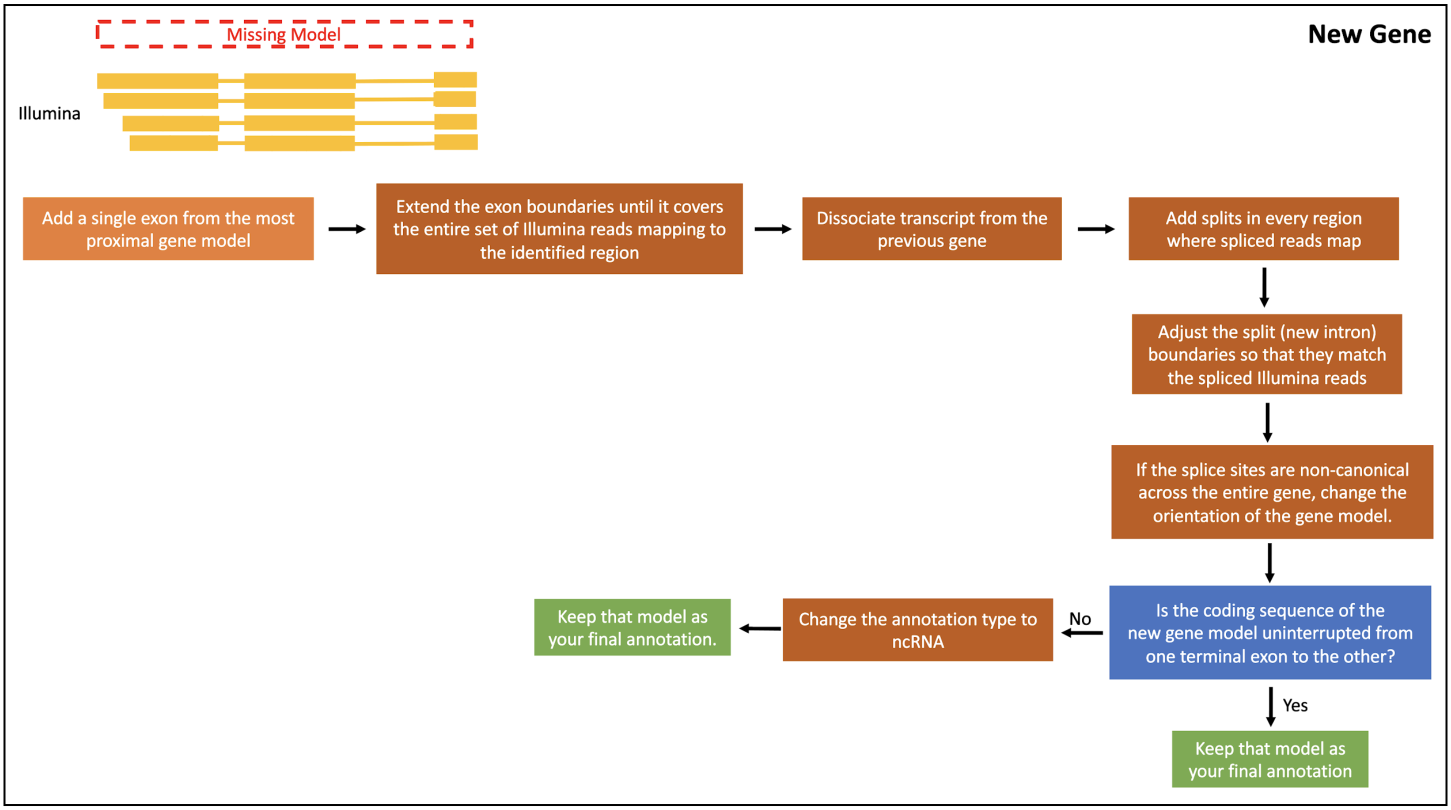

1. Add a single exon from the most proximal gene model
2. Extend the exon boundaries until it covers the entire set of Illumina reads mapping to the identified region
3. Dissociate the transcript from the previous gene
    - This will assign a unique gene ID to the new transcript
4. Add splits in every region where spliced reads map
    - These are putative exons
5. Adjust the split boundaries so that they match the splice sites of the Illumina data
6. If the splice sites are non-canonical across the entire gene, switch the orientation of the gene
    - Considering that this new gene is based on paired-end reads, we don't know the actual orientation.
    - If the non-canonical splice sites persist after changing orientation, discard the gene model and move onto the next gene
7. Is the coding sequence of the new gene model uninterrupted from one terminal exon to the other? (Yes)
    - If not, you would change the annotation type to ncRNA, add metadata, and move on.
8. Keep that model as your final annotation
9. Fill curation metadata (Section 2.5.), and move onto the next gene.
    - Metadata code for this case: 'New Gene - B:S'

#### 2.5. Curation Metadata

After each gene curation, you will record two pieces of information about the curation process you carried out:

1. What type of error was found in the gene model?
2. What sources of evidence did you use to **fix the error**?

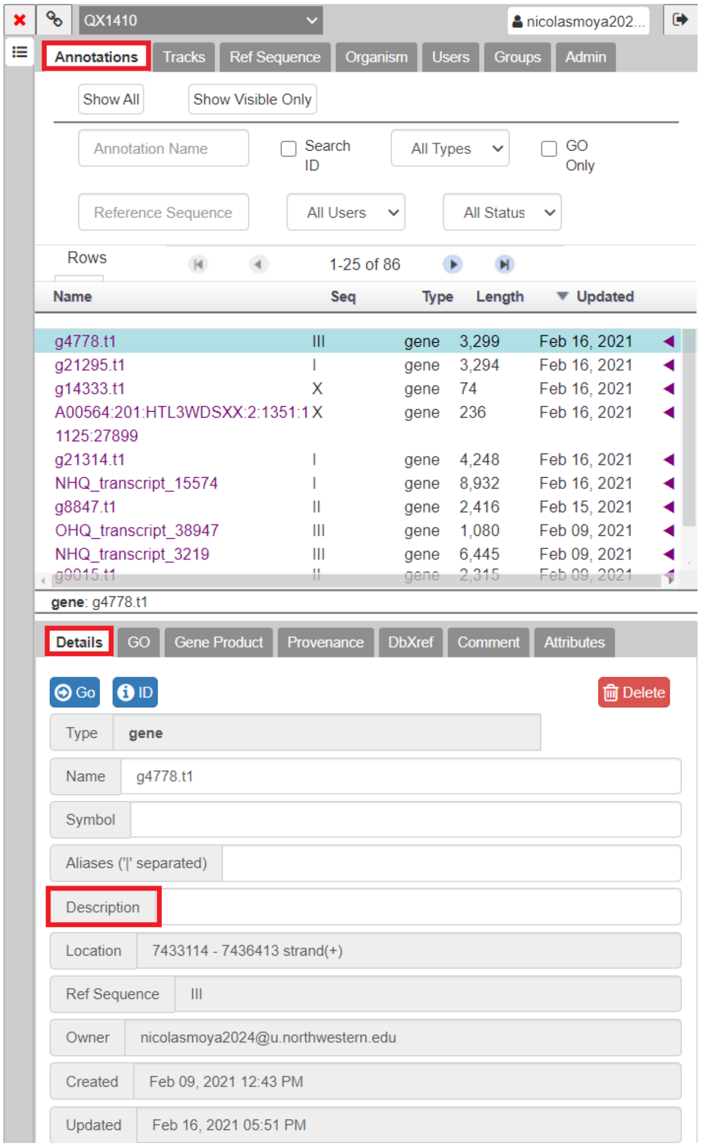


Curated models in the User-created annotations space will appear in the Annotations tab of the Annotator Panel. After selecting the curated model from the list in the Annotations tab, the details of the model will be prompted. In the model description box, you will add the type of error and sources of evidence in the format shown below.

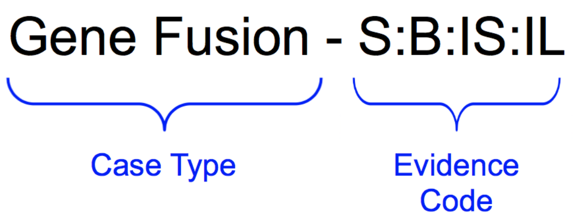


**Case type:**

- Simple
- Low Coverage
- UTR Extension
- Gene Fusion
- Gene Split

**Evidence codes:**

- B = BRAKER
- S = StringTie
- IL = Illumina
- IS = IsoSeq

After this information has been recorded, you can proceed to curate the next gene. Congratulations!

#### 2.6. New cases

As you work in the curation project, you may encounter new and unique cases that are not described in this protocol. New cases will be presented in team meetings or through Slack. We will analyze each case and create a workflow to solve these new problems. The new workflows will be added to the protocol and shared with the curation team.
