## Supplemental Information 2 for "Novel and improved *Caenorhabditis briggsae* gene models generated by community curation"

### *Caenorhabditis briggsae* gene annotations improved by community curation using short- and long-read RNA sequencing

**Supplemental Information 2**

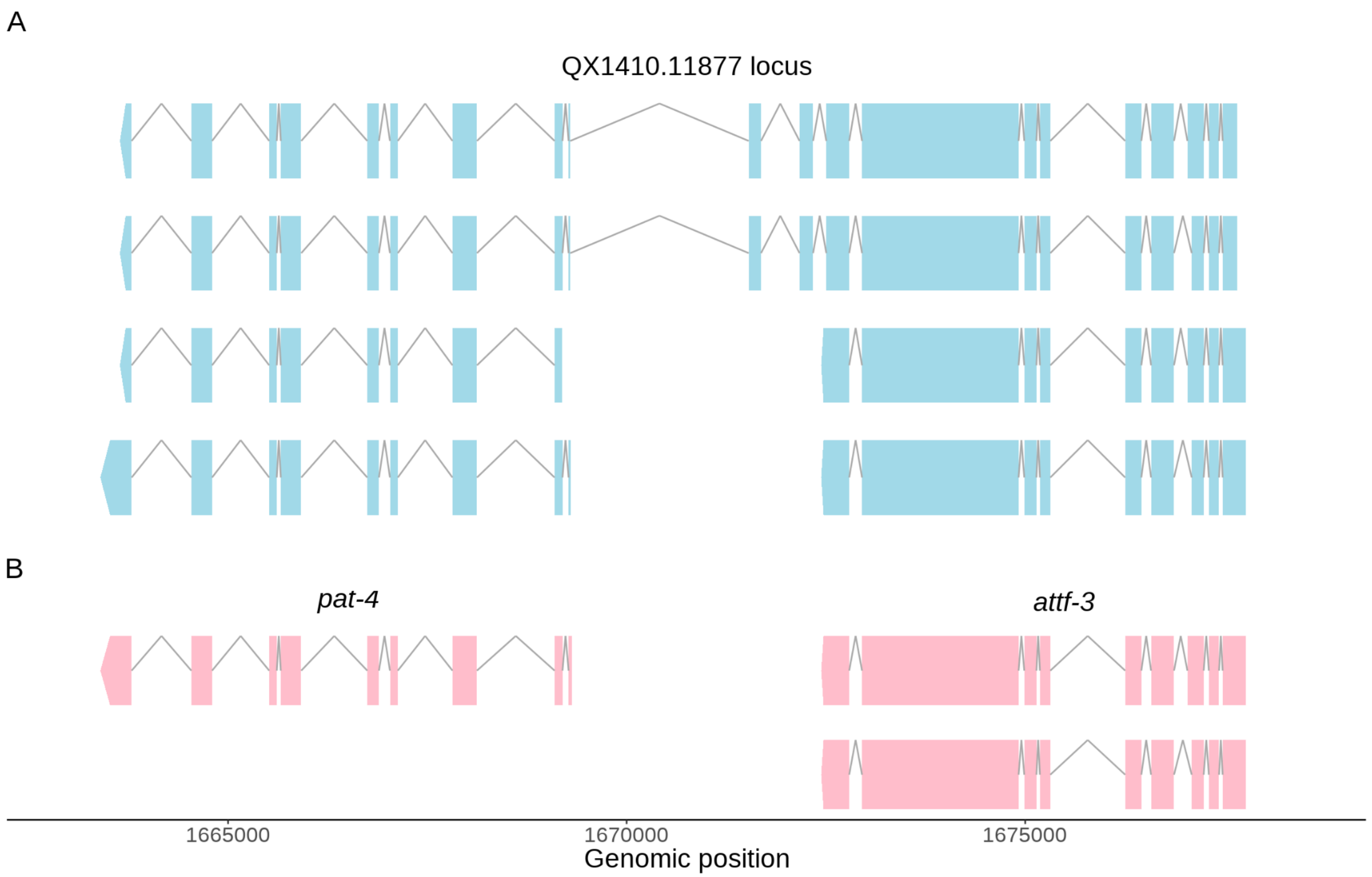


**Supplemental Figure 1: Example of BUSCO duplication reduction after manual curation**
Plot showing the gene models located on chromosome III from 1,663,398 to 1,677,797 before (A) and after (B) curation. Six software-derived models are shown in blue and are annotated as isoforms of a single gene. Three spurious isoforms were removed during the curation process, and the locus was divided into two separate genes, orthologous to the *C. elegans pat-4* and *attf-3* genes. The *pat-4* gene is part of the BUSCO database, and it is no longer duplicated. The *attf-3* gene is composed of two isoforms that have differences in the 3' termini of exon 3.

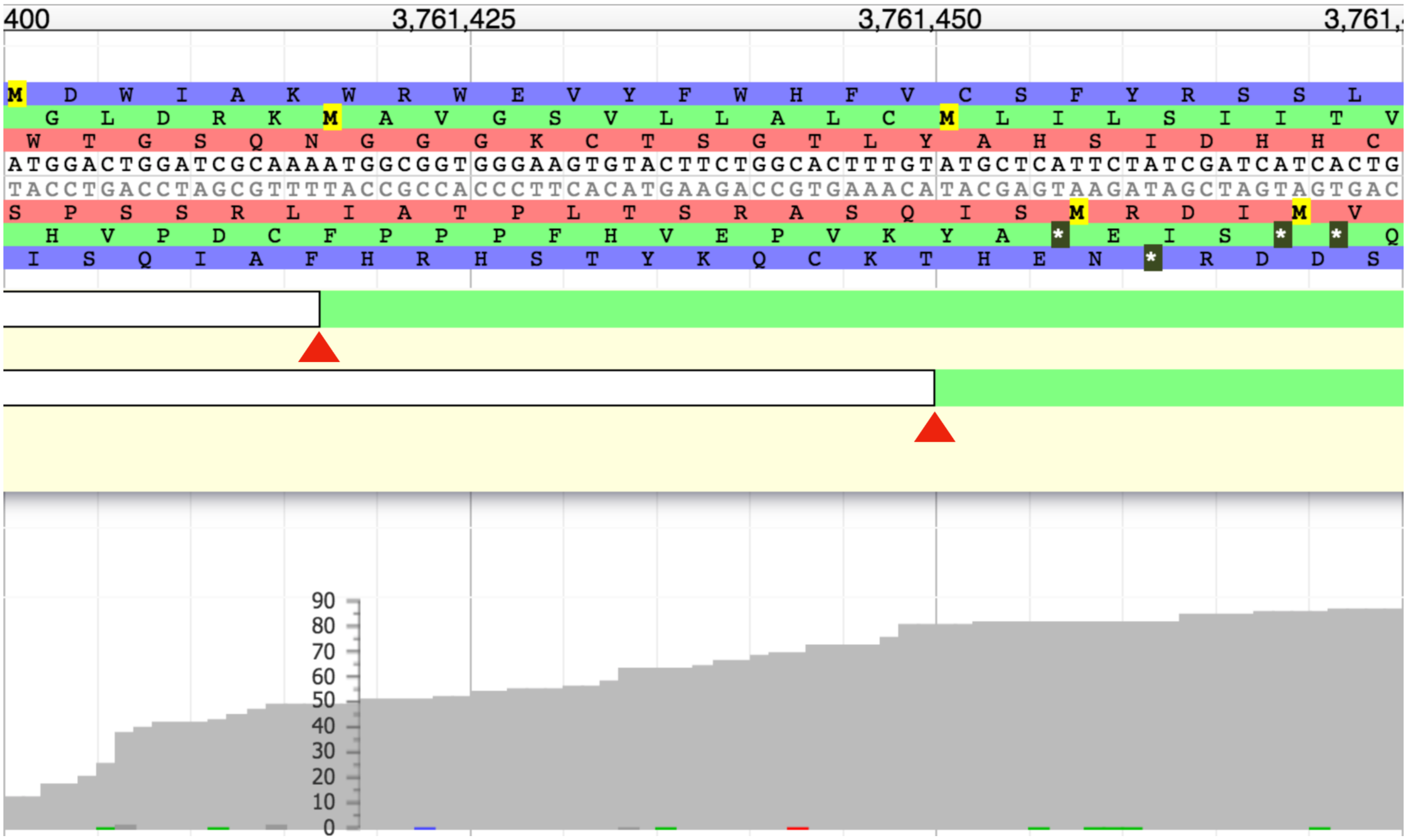


**Figure S2: Example of alternative translation initiation site**
Screen capture showing a genomic window of 75bp (located on chromosome III from 3,761,400 to 3,761,475) that covers the 5' termini of the *gar-2 C. briggsae* ortholog. Two alternative in-frame translation initiation sites are highlighted with red arrows. Both sites have similar RNA coverage (shown in gray).

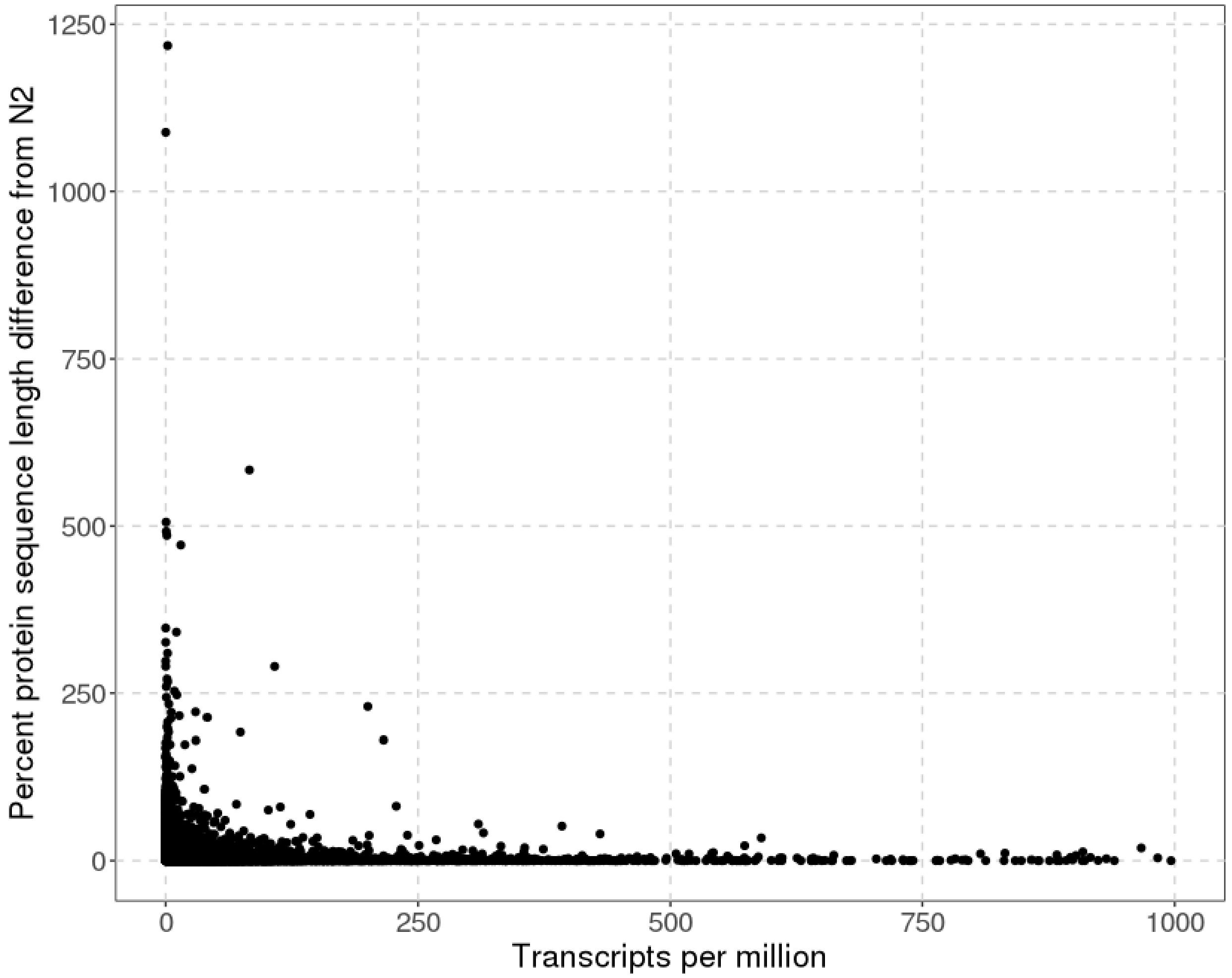


**Figure S3: Protein-length accuracy of QX1410 genes by TPM**
Plot showing the transcripts per million for each QX1410 gene against its percent protein length distance to its N2 ortholog.

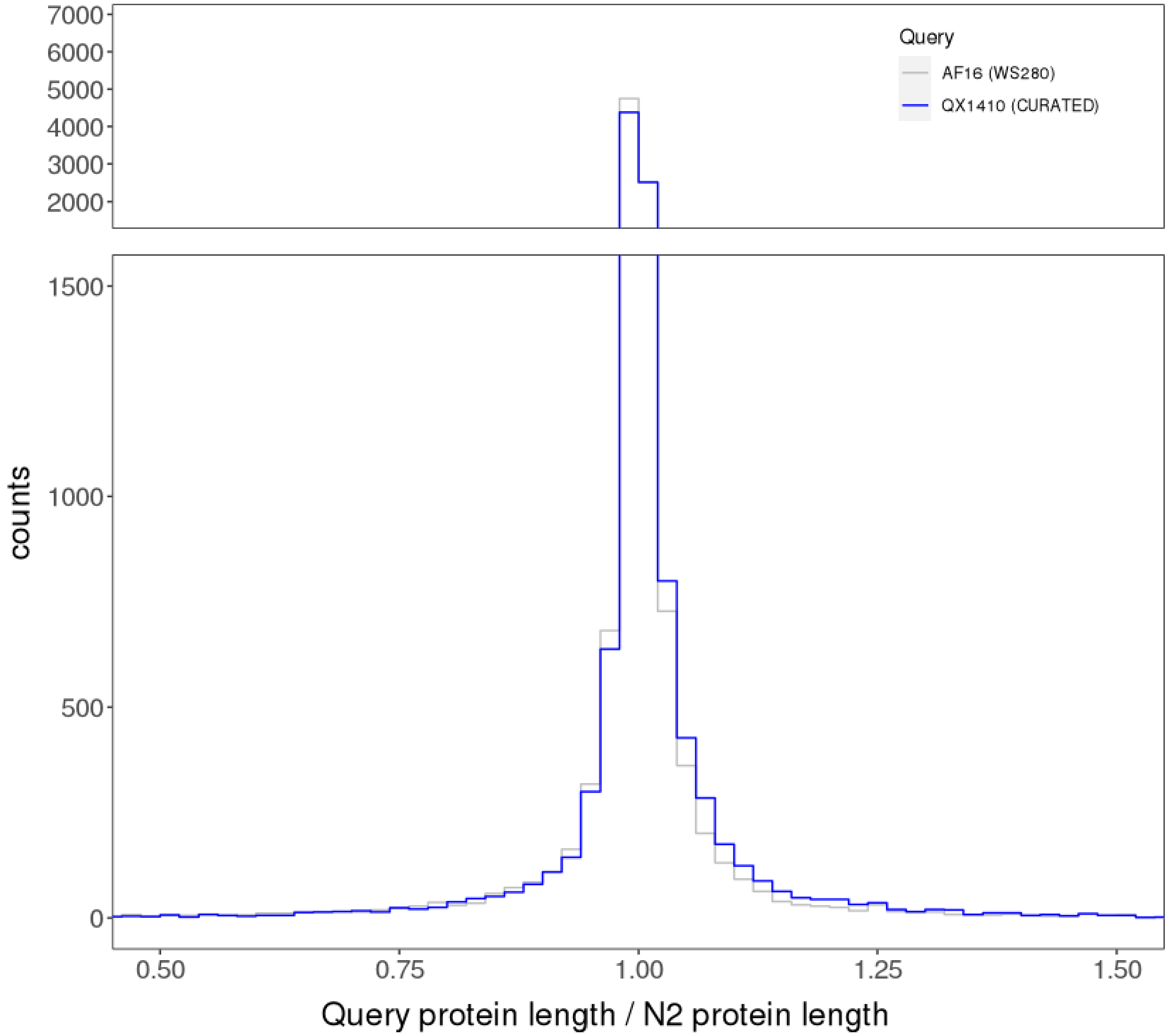


**Figure S4: Protein length accuracy of *C. briggsae* gene models relative *C. elegans* N2 inferred single-copy orthologs**

We calculated the protein length accuracy of both curated QX1410 gene models and AF16 (WS280) gene models using single-copy orthologs identified through phylogenetic orthology analysis. A ratio of one represents a *C. briggsae* gene with a protein length that perfectly matches its single-copy ortholog in N2.

**Table S1: Protein-length accuracy and BUSCO completeness before and after revision of 755 loci with deviations in protein length**

| | Before revision | After revision |
|---|---|---|
| one-to-one matches to N2 | 2601 | 2710 |
| 5% off matches to N2 | 7849 | 7940 |
| BUSCO completeness | 99.7% | 99.7% |

**Table S2: Protein-length accuracy using phylogeny-inferred orthologs between QX1410, AF16, and N2.**

| | AF16 | QX1410 |
|---|---|---|
| one-to-one matches to N2 | 2222 | 2173 |
| 5% off matches to N2 | 8762 | 8811 |

**Table S3: Intra-chromosomal duplications in the AF16 genome and affected genes**

| segment A | segment B | identity | genes in segment A | genes in segment B |
|---|---|---|---|---|
| I:12895-14211 | I:396029-397345 | 99.57 | Cbr-rps-29 | CBG11412 |
| I:175947-178569 | I:169983-172605 | 99.7 | CBG11861, CBG00361, CBG11492 | Cbr-arrd-2, Cbr-sbds-1, CBG11493 |
| I:134401-138262 | I:126814-130653 | 99.77 | CBG01029, CBG15263, Cbr-ncs-3 | Cbr-lim-6 |
| I:124226-127163 | I:131671-134750 | 99.42 | CBG30931, Cbr-pfn-3 | CBG15264 |
| I:50373-56695 | I:42341-48591 | 99.62 | CBG01049, CBG01048, CBG01046, CBG11525, CBG11524 | Cbr-glt-5, CBG00337, Cbr-fmo-4 |
| I:1171622-1178466 | I:1218277-1225147 | 96.33 | CBG16333 | CBG18556, CBG00846, CBG00845, Cbr-hpo-38, Cbr-acr-15, CBG16326 |
| I:1352610-1357683 | I:1361512-1366585 | 99.22 | CBG00754, CBG27416 | CBG15737, CBG27418 |
| I:1463157-1465588 | I:1466607-1469038 | 97.69 | CBG00785 | CBG31329, CBG00786 |
| I:1678479-1681854 | I:1700745-1704120 | 99.27 | CBG25180 | CBG30327 |
| I:1807933-1810419 | I:1811400-1813886 | 96.44 | CBG30809 | CBG22020, CBG08816 |
| I:2001351-2004701 | I:2006517-2009867 | 97.19 | CBG21555, CBG21557, CBG31291 | CBG21559 |
| I:2006686-2009893 | I:2001512-2004758 | 97.81 | CBG21559 | CBG21557, CBG31291 |
| I:3219715-3226441 | I:2424060-2430786 | 99.9 | CBG29818, CBG11638, CBG17031, CBG30501, Cbr-tmem-218 | Cbr-pqn-59.1, CBG30439, CBG26638 |
| I:2797516-2798692 | I:2802327-2803504 | 99.83 | CBG22202 | CBG22203 |
| I:3208166-3215423 | I:3275534-3282791 | 99.72 | Cbr-smo-1, CBG22302, Cbr-glna-2, CBG26833 | CBG15102, CBG15103, CBG15104, CBG13342, CBG10509, CBG10508 |
| I:3456270-3466382 | I:3456270-3466382 | 99.98 | Cbr-xrep-4, Cbr-trpp-8, CBG14903, CBG00725, CBG24735, Cbr-pes-1, CBG10454, Cbr-gnrr-3 | Cbr-xrep-4, Cbr-trpp-8, CBG14903, CBG00725, CBG24735, Cbr-pes-1, CBG10454, Cbr-gnrr-3 |
| I:3560841-3566940 | I:3567941-3574040 | 99.8 | Cbr-elo-8, Cbr-srg-3, CBG17473 | CBG30964, CBG22594, CBG10425, CBG01955 |
| I:4963727-4964839 | I:15222377-15223491 | 94.92 | CBG03918 | CBG17036 |
| I:5594552-5601685 | I:5606871-5614004 | 99.98 | CBG03762, CBG03761, CBG03760, Cbr-fip-3.1, CBG18231, Cbr-exos-1, CBG17865, CBG30898 | CBG03699, Cbr-cel-1, CBG17869, CBG17870 |
| I:8006840-8008057 | I:12076549-12077849 | 85.01 | CBG31390 | CBG00002 |
| I:5779531-5781829 | I:5830921-5833219 | 99.91 | CBG30597, Cbr-ttr-31 | CBG27553 |

| | | | | |
|---|---|---|---|---|
| I:1673321-1674689 | I:13084727-13086094 | 99.78 | CBG26161 | CBG16717 |
| I:6706546-6707749 | I:6708365-6709568 | 99.83 | Cbr-mzt-1.2 | Cbr-mzt-1.1, CBG03284 |
| I:6867179-6869525 | I:6858657-6860983 | 92.55 | Cbr-wee-1.1, CBG06567 | CBG21906, Cbr-cnc-4, Cbr-cnc-3 |
| I:6858655-6860982 | I:6867180-6869526 | 92.64 | CBG21906, Cbr-cnc-4, Cbr-cnc-3 | Cbr-wee-1.1, CBG06567 |
| I:7884100-7891151 | I:7028193-7035244 | 99.97 | CBG12104, CBG12105, Cbr-viro-2, Cbr-pot-1, CBG26353, CBG05826, Cbr-swt-7 | CBG11902, CBG11903, CBG30143, Cbr-paqr-1, CBG06610, Cbr-nhr-178, CBG06612 |
| I:7233822-7241056 | I:7245702-7252936 | 99.96 | Cbr-spe-4, Cbr-fut-1, CBG17973, CBG26323, Cbr-adk-1, CBG27367, CBG27601 | CBG11945, CBG11946, Cbr-tars-1, CBG06026, Cbr-rap-3 |
| I:7276238-7279267 | I:7892152-7895182 | 100 | CBG03133, CBG17962, CBG17136 | CBG26592, CBG27617 |
| I:7338216-7343793 | I:7344751-7350328 | 99.6 | CBG25559, CBG30033, CBG05994, CBG05993, CBG17119 | Cbr-flap-1, CBG06687 |
| I:7523170-7526884 | I:7527845-7531559 | 99.95 | CBG12018, Cbr-wip-1, CBG11828 | CBG06740 |
| I:8486251-8487568 | I:8484615-8485932 | 99.85 | CBG29853 | CBG29852 |
| I:8484614-8485932 | I:8486251-8487569 | 99.92 | CBG29852 | CBG29853 |
| I:8433926-8435390 | I:8005537-8007001 | 99.84 | CBG14192 | CBG02960 |
| I:8002165-8007001 | I:8427483-8432320 | 99.44 | CBG23931, Cbr-pdp-1.1, Cbr-rps-9, CBG02961, CBG02960, CBG05790, CBG08616 | CBG12192, Cbr-pdp-1.2, CBG23522 |
| I:8369874-8372703 | I:8364987-8367816 | 99.92 | CBG12206, CBG06038 | CBG12207 |
| I:8146914-8148994 | I:8712277-8714346 | 87.66 | CBG25949 | CBG12411 |
| I:8048439-8057284 | I:8027971-8036816 | 99.95 | CBG02255, CBG02254, CBG02253, Cbr-bigr-1, Cbr-lir-3, CBG02950, Cbr-str-48, CBG08598, CBG08597 | CBG24754, Cbr-hsp-12.1, CBG24756, CBG24757, Cbr-pqm-1, CBG08607, CBG30625, CBG08605 |
| I:8712279-8714362 | I:8146901-8148995 | 87.7 | CBG12411 | CBG25949 |
| I:9828246-9833872 | I:9835223-9840863 | 99.98 | Cbr-nuo-2, CBG30979, Cbr-dhp-2, CBG03445 | CBG29118, CBG12673, CBG26395 |
| I:10596874-10605615 | I:10606616-10615357 | 99.96 | Cbr-txt-9.1, CBG02319, Cbr-ptr-9, CBG01749, CBG01748, Cbr-set-22, Cbr-snf-10, Cbr-pap-1 | CBG02316, CBG31350, CBG02315, Cbr-nhl-1, CBG01746, Cbr-rpl-20, Cbr-cyc-2.1, Cbr-tctn-1, CBG09606 |
| I:11017779-11018528 | I:11026776-11027525 | 99.8 | Cbr-msrp-1 | Cbr-msrp-6 |
| I:11035842-11039634 | I:11078044-11081817 | 97.83 | CBG20251, CBG01632 | CBG20242, CBG30532, CBG09452 |

| | | | | |
|---|---|---|---|---|
| I:11078047-11081816 | I:11035857-11039634 | 97.81 | CBG20242, CBG30532, CBG09452 | CBG20251, CBG01632 |
| I:11080655-11081727 | I:11231511-11232585 | 89.94 | CBG20242 | CBG08500 |
| I:11399713-11403810 | I:11571154-11575259 | 99.12 | Cbr-hpo-15 | CBG13610, CBG17707, Cbr-str-27, Cbr-odd-2 |
| I:11571173-11575235 | I:11399712-11403810 | 98.95 | CBG13610, CBG17707, Cbr-odd-2 | Cbr-hpo-15 |
| I:12365185-12369064 | I:12507990-12511869 | 99.99 | CBG20401, CBG20402, CBG10926 | CBG20432, CBG20433, CBG19627 |
| I:13001070-13005877 | I:3457-8335 | 95.38 | CBG19165 | Cbr-sqv-8, CBG01066, CBG00325, Cbr-grl-17 |
| I:12722206-12726069 | I:12837614-12841445 | 99.17 | Cbr-fipr-24.2, CBG30066, Cbr-map-1 | CBG18839 |
| I:12548002-12554217 | I:13061341-13067525 | 99.24 | CBG19620, CBG08368, Cbr-odc-1, CBG30805 | CBG30069, CBG19495, CBG21194, CBG19186, CBG19187, CBG19188, CBG27815 |
| I:13812723-13815134 | I:13809420-13811856 | 99.59 | CBG18489 | CBG30593, CBG19371, CBG16882 |
| I:13809433-13811847 | I:13812715-13815148 | 99.88 | CBG19371, CBG16882 | CBG18489 |
| I:14796277-14799080 | I:14801167-14803970 | 95.19 | CBG04652 | CBG04655 |
| I:15222383-15223530 | I:4963727-4964841 | 94.57 | CBG17036 | CBG03918 |
| I:15257988-15265658 | I:15249949-15257621 | 99.79 | CBG04781 | CBG16021, Cbr-tbx-34, CBG31278, Cbr-mtl-2, CBG04780 |
| II:66677-71747 | II:72748-77818 | 99.86 | Cbr-sde-2, CBG11515, CBG08049 | CBG00342 |
| II:512064-515307 | II:515675-518922 | 93.72 | CBG13813, Cbr-egl-17 | CBG26871 |
| II:1216389-1220711 | II:1133917-1138239 | 99.06 | CBG30967, CBG00846, CBG15718, CBG16326 | CBG00857 |
| II:1121882-1129102 | II:1199474-1206680 | 98.45 | CBG18542, Cbr-sfxn-1.2, CBG30324, Cbr-oac-29, CBG16345 | CBG30969, Cbr-taco-1, CBG15715 |
| II:1488444-1491181 | II:1484260-1486997 | 99.62 | CBG00797 | CBG00795 |
| II:1595824-1596950 | II:1611602-1612728 | 99.3 | CBG26157 | CBG26158 |
| II:1592359-1596028 | II:1612521-1616190 | 99.84 | CBG16241 | CBG05101, CBG00839, CBG00840, CBG27433 |
| II:1778662-1781943 | II:1855128-1858523 | 99.76 | CBG18444, CBG29999, Cbr-josd-1, CBG00487 | CBG31002 |
| II:1989928-1991960 | II:1994663-1996695 | 95.07 | CBG29812 | CBG08875 |
| II:2314636-2315656 | II:13733152-13734152 | 94.26 | CBG31124 | CBG19348 |
| II:2426866-2430072 | II:2434507-2437713 | 97.35 | CBG30439, CBG26638 | CBG30496 |
| II:2865336-2870299 | II:2901580-2906538 | 100 | CBG25484 | CBG22230, CBG13419, CBG13418, CBG22362, |

| | | | | |
|---|---|---|---|---|
| | | | | Cbr-srj-7, Cbr-srj-11, CBG15947 |
| II:2880915-2884271 | II:3069972-3073328 | 99.94 | CBG13424, CBG22357, Cbr-srj-9 | CBG22271, CBG13380, CBG13379, CBG16983 |
| II:2885288-2892119 | II:3262407-3269238 | 99.63 | CBG13423, CBG25829, CBG22359, CBG22421, Cbr-sdz-34.4 | CBG15100, CBG13343, Cbr-nmt-1, CBG10514, CBG01894 |
| II:4158933-4168350 | II:4314672-4324083 | 99.64 | CBG04167, CBG00542, CBG00543, CBG09040, CBG09039, Cbr-lipl-5, CBG26990, CBG26989, Cbr-nlp-34.4, CBG26988 | Cbr-cep-1, Cbr-rpl-25.2, CBG04079, Cbr-wrm-1, CBG30196, CBG26982, CBG26981, CBG02102, CBG02103 |
| II:4144222-4148435 | II:4139998-4144211 | 98.55 | CBG00540 | CBG04172, Cbr-dnc-6, CBG31324, CBG00538, Cbr-erd-2.2 |
| II:4823536-4828914 | II:4834010-4839388 | 99.63 | Cbr-coa-4, CBG01159 | CBG04252, CBG31007, CBG31179, CBG21694, Cbr-srj-33, CBG30816 |
| II:4838194-4839388 | II:4839399-4840597 | 98.25 | CBG21694 | CBG04254 |
| II:4842262-4845929 | II:4830114-4833782 | 99.89 | CBG01152, Cbr-ftt-2 | CBG01158 |
| II:5304613-5308357 | II:5308458-5312191 | 99.97 | CBG12122, CBG12123 | CBG11131, Cbr-hex-3, CBG25998, CBG12124 |
| II:5819690-5824660 | II:5825661-5830631 | 99.98 | Cbr-stl-1, CBG13080, CBG13183, CBG17917, CBG27385 | CBG17919, CBG17920, Cbr-col-177 |
| II:6564597-6570699 | II:6576189-6582283 | 99.98 | CBG10822, CBG10823, CBG10824, Cbr-cki-2, CBG18137, CBG30316, CBG27086, Cbr-str-46, CBG06463 | Cbr-mat-1, CBG12945, Cbr-pqn-96, Cbr-ufbp-1, Cbr-ifc-1, Cbr-str-39 |
| II:6572050-6576178 | II:6582297-6586425 | 99.99 | CBG12946, Cbr-srw-4, Cbr-srw-3 | CBG10829, Cbr-pfkb-1.1, Cbr-rpn-3, CBG04479 |
| II:7392256-7393219 | II:9662010-9663024 | 91.8 | CBG30348 | CBG30738 |
| II:7470239-7475817 | II:7484413-7489991 | 99.64 | CBG25565, Cbr-clec-61, CBG05950, CBG05949, Cbr-cpz-2, CBG11812, CBG26804 | Cbr-unc-108, CBG03093, CBG03092, CBG25922, Cbr-nhr-8, CBG30726 |
| II:9662013-9663056 | II:7392255-7393260 | 91.6 | CBG30738 | CBG30348 |
| II:10052028-10056936 | II:10057956-10062864 | 99.96 | Cbr-mdt-26, Cbr-ift-43.1, Cbr-rpl-25.1 | CBG12716, CBG02468, Cbr-ift-43.2, Cbr-scl-1, Cbr-scl-9 |
| II:10109241-10116736 | II:10119713-10127208 | 99.95 | CBG29868, CBG12731, Cbr-cdc-14.2, Cbr-jmjd-3.2, Cbr-hum-10 | CBG12733, CBG12734, Cbr-ttc-4, Cbr-cdc-14.1, CBG25640, CBG24368, CBG24369, CBG24371, CBG09743, CBG30526, CBG27068 |

| | | | | |
|---|---|---|---|---|
| II:10185727-10189076 | II:10192435-10195784 | 99.79 | CBG02428, CBG26018, CBG06900 | CBG12747, CBG02426, CBG14554 |
| II:10189076-10191758 | II:10198301-10200983 | 99.95 | CBG02427, CBG06899 | CBG25369, CBG02425, CBG06896 |
| II:10198299-10200981 | II:10189076-10191760 | 100 | CBG25369, CBG02425, CBG06896 | CBG02427, CBG06899 |
| II:10281451-10283198 | II:10161077-10162798 | 96.56 | CBG02400 | Cbr-sec-61.G |
| II:11375377-11381801 | II:11385306-11391730 | 99.72 | CBG08467, CBG08466, CBG25403, CBG20787, CBG20788, CBG14803 | CBG08462, CBG25402, CBG20790, CBG20791, CBG09383 |
| II:12296859-12303234 | II:12303245-12309620 | 98.92 | Cbr-mtx-2, CBG00034, CBG30446, CBG18975, CBG18976, Cbr-asg-2 | Cbr-csn-4, Cbr-gon-14, CBG10907, CBG10908 |
| II:11667151-11673571 | II:11680357-11686777 | 97.8 | CBG20860, Cbr-mvb-12, CBG09304, CBG05010 | CBG26030, CBG20145, Cbr-ric-3, CBG09301, CBG05015 |
| II:11930967-11935132 | II:12407061-12411226 | 99.8 | CBG30118 | CBG20410, CBG20411, CBG31464, CBG30650 |
| II:12416274-12422208 | II:11935143-11941077 | 99.86 | CBG20414, CBG12246, Cbr-grl-8 | CBG17611, Cbr-sdz-13, CBG05074, CBG05075 |
| II:12076340-12077955 | II:11933037-11934598 | 97.26 | CBG00002 | CBG30118 |
| II:12264279-12266701 | II:12423299-12425723 | 93.86 | CBG20995 | CBG31280 |
| II:12427940-12429153 | II:12430973-12432180 | 87.31 | CBG21037 | CBG21039 |
| II:12430977-12432176 | II:12427936-12429154 | 88.91 | CBG21039 | CBG21037 |
| II:12908766-12912742 | II:12439803-12443779 | 99.94 | CBG31528 | Cbr-ufm-1 |
| II:12444663-12450908 | II:12914549-12920794 | 99.84 | Cbr-rpl-35, Cbr-ugt-59, CBG19012, Cbr-bbs-8 | CBG29933, CBG31529, CBG19532, CBG30250, CBG21224, CBG26479, CBG26478, CBG11046 |
| II:12453480-12457714 | II:12459869-12464047 | 98.02 | CBG20418, CBG19643, Cbr-srh-30, Cbr-srh-28 | CBG19641, Cbr-ugt-44, Cbr-spn-4, Cbr-tsp-16 |
| II:12459833-12464050 | II:12453480-12457672 | 96.3 | CBG19641, Cbr-ugt-44, Cbr-spn-4, Cbr-tsp-16 | CBG20418, CBG19643, Cbr-srh-30, Cbr-srh-28 |
| II:12466977-12473539 | II:12931949-12938511 | 99.71 | CBG20421, CBG19638, CBG08387, CBG19020 | CBG21221, Cbr-str-84, CBG19146, CBG11049 |
| II:13018194-13023072 | II:13011860-13016763 | 99.84 | CBG19507, CBG19171, CBG30543, CBG19173, Cbr-swm-1 | CBG18879, CBG19510, CBG26473, Cbr-nhr-30.1, CBG27177 |
| II:13101976-13105215 | II:13106002-13109244 | 96.42 | CBG19486, Cbr-sri-40, CBG10726, CBG19199, CBG19200 | CBG19483, CBG26165, CBG21186 |
| II:13106002-13109823 | II:13101394-13105215 | 99.61 | CBG19483, CBG26165, CBG21186, CBG30547, Cbr-col-174 | CBG19486, Cbr-sri-40, CBG10726, CBG30546, CBG19199, CBG19200 |
| II:13184527-13187950 | II:13188328-13191751 | 99.95 | CBG21174, CBG31168 | Cbr-atp-2 |

| | | | | |
|---|---|---|---|---|
| II:13195818-13199676 | II:13165668-13169526 | 99.94 | CBG04349 | CBG29899 |
| II:13210515-13212263 | II:13201048-13202824 | 97.81 | Cbr-nlp-61 | CBG04350 |
| II:12099401-12100628 | II:14523348-14524575 | 92.86 | CBG20948 | CBG21869 |
| II:14523348-14525105 | II:14527276-14529035 | 93.09 | CBG21869 | CBG24810 |
| II:14526707-14528432 | II:15101545-15103253 | 89.57 | CBG24810 | CBG17057 |
| II:15101738-15103253 | II:14203501-14205016 | 87.68 | CBG17057 | CBG10347 |
| II:14203502-14204860 | II:14420582-14421965 | 92.36 | CBG10347 | CBG10283 |
| II:14420582-14421793 | II:12099489-12100654 | 86.52 | CBG10283 | CBG20948 |
| II:13733152-13734152 | II:2314652-2315656 | 95.24 | CBG19348 | CBG31124 |
| II:13973009-13974032 | II:13984297-13985291 | 86.54 | Cbr-ins-32.2 | Cbr-prx-11, Cbr-ins-32.5 |
| II:14150197-14151510 | II:14143404-14144716 | 98.17 | CBG07912 | CBG07911 |
| II:14557696-14560000 | II:14568356-14570614 | 94.38 | CBG24149 | CBG16447, CBG31150, CBG04587 |
| II:14808172-14809274 | II:14871003-14872119 | 97.77 | CBG21442 | CBG21457 |
| II:15294684-15296640 | II:15284353-15286275 | 95.88 | CBG16034 | Cbr-flp-33, CBG16029 |
| II:15284353-15286278 | II:15294670-15296630 | 97.96 | Cbr-flp-33, CBG16029 | CBG16034 |
| II:16600308-16607497 | II:16607799-16614988 | 99.97 | CBG06958, Cbr-drl-1, Cbr-srxa-7 | Cbr-dhs-6, CBG30393 |
| III:251831-254547 | III:259312-262037 | 99.93 | CBG15234 | CBG11878, CBG00994 |
| III:332180-336194 | III:336342-340356 | 99.93 | CBG11438, CBG11437 | Cbr-srx-42 |
| III:528787-534740 | III:534751-540704 | 98.74 | Cbr-str-144 | Cbr-catp-3 |
| III:958113-962780 | III:1627831-1632509 | 99.96 | CBG11267 | CBG15793, CBG08765, CBG16229 |
| III:1649736-1656389 | III:1144021-1150674 | 99.92 | Cbr-gsto-3.1, CBG15798, Cbr-srt-44.1, CBG13846, CBG16223 | Cbr-gsto-3.2 |
| III:1658437-1664270 | III:1538017-1543850 | 99.94 | CBG05095, CBG31062, CBG13848, Cbr-mfsd-11 | Cbr-psf-3, CBG29807, CBG00814 |
| III:6811048-6814310 | III:1755904-1759159 | 99.72 | CBG21890, CBG30952 | CBG00479 |
| III:1880950-1883530 | III:1885400-1888002 | 97.36 | CBG22029 | Cbr-spp-23 |
| III:3406569-3412108 | III:3132111-3137650 | 98.45 | CBG11677, CBG27477 | CBG17009, Cbr-sup-10 |
| III:3177475-3184954 | III:3184965-3192444 | 99.52 | CBG22296, CBG21090, CBG31277, CBG21093, Cbr-str-164, CBG30649, CBG17019 | CBG21094, CBG31276, Cbr-str-163, CBG31037, CBG17021, Cbr-lec-9 |
| III:3936476-3944683 | III:3822463-3830670 | 99.96 | CBG04123, CBG04124, CBG29824, Cbr-tofu-4, CBG09097, CBG01426, CBG01425 | Cbr-itx-1, CBG09118, Cbr-cutl-23, Cbr-cav-1 |
| III:3940131-3944683 | III:3945684-3950236 | 99.74 | CBG29824, Cbr-tofu-4, CBG01425 | Cbr-mage-1, CBG09096 |
| III:3983939-3985307 | III:4118576-4119916 | 90.88 | CBG09091 | CBG09051, CBG30336 |

| | | | | |
|---|---|---|---|---|
| III:4529767-4531348 | III:4496830-4498393 | 84.98 | CBG05212 | CBG08958 |
| III:4557040-4562953 | III:4517123-4523036 | 99.91 | Cbr-kbp-3, CBG26141, CBG01239 | Cbr-ceh-12, Cbr-col-54, Cbr-nlp-62, CBG30197, CBG21752, CBG21751, CBG01253, Cbr-sav-1 |
| III:4629492-4631486 | III:4624163-4626157 | 99.94 | CBG05182 | CBG01217 |
| III:4792753-4797708 | III:4797809-4802764 | 99.93 | CBG04244, CBG23800, CBG29116, CBG01169 | Cbr-ept-1, CBG25513, CBG24620, CBG21708, CBG01168 |
| III:5008803-5010747 | III:5130826-5132779 | 99.28 | CBG18376 | CBG18345 |
| III:5130808-5132791 | III:5008794-5010746 | 99.23 | CBG18345 | CBG18376 |
| III:5195994-5198512 | III:4989036-4991546 | 100 | CBG18325 | Cbr-mrpl-12, CBG18379, CBG01113 |
| III:5570162-5575549 | III:6044340-6049727 | 99.29 | Cbr-ikb-1, CBG18240 | Cbr-edg-1, CBG13233, Cbr-srbc-58 |
| III:5573187-5575346 | III:6049734-6051898 | 97.56 | CBG18240 | CBG13234 |
| III:6049737-6051898 | III:5573185-5575346 | 99.78 | CBG13234 | CBG18240 |
| III:5793019-5797080 | III:5798081-5802142 | 99.97 | Cbr-npr-31, CBG30598 | CBG30025, Cbr-fust-1 |
| III:5883416-5890760 | III:5891471-5898815 | 99.97 | Cbr-bro-1, CBG26571, CBG13099, CBG26150, CBG30027, CBG13100, CBG30843, CBG19948, Cbr-srx-22, Cbr-irld-3, CBG17543, Cbr-nlp-19 | Cbr-npp-6, CBG13103, CBG26414, CBG06281, CBG06282, CBG17546 |
| III:6351606-6359943 | III:6364385-6372722 | 99.97 | Cbr-pgam-5, CBG13314, Cbr-trap-4.1, CBG13317, CBG06415, CBG06416 | Cbr-paf-1, CBG29840, CBG13319, Cbr-ccf-1, Cbr-trap-4.2, CBG13322, Cbr-ges-1, CBG06419 |
| III:6503869-6505020 | III:6505647-6506829 | 95.88 | CBG18155, Cbr-rps-25 | CBG18154 |
| III:6505649-6506802 | III:6503867-6505047 | 95.88 | CBG18154 | CBG18155, Cbr-rps-25 |
| III:6809503-6814263 | III:1754325-1759159 | 98.58 | CBG18081, CBG30952, Cbr-srv-24 | CBG30277, CBG30278, CBG00479 |
| III:7871256-7875249 | III:7875575-7879568 | 99.97 | CBG31459, CBG08655 | CBG16596, CBG08654, CBG08653, CBG08652 |
| III:9037020-9037720 | III:9034754-9035454 | 99.88 | CBG09918 | CBG09917 |
| III:9202714-9206259 | III:9207273-9210818 | 99.67 | CBG02683, CBG09964, CBG14340 | CBG14341 |
| III:9245530-9251347 | III:9306132-9311949 | 99.91 | Cbr-slc-25A42, CBG12502, Cbr-lmd-1, Cbr-rsp-5, Cbr-snr-3, Cbr-vps-53, CBG09979, CBG09980, CBG30353, Cbr-ppm-1.H, Cbr-rpl-34 | CBG02655, CBG10002, CBG10004, CBG14364, CBG14365 |
| III:9551641-9553240 | III:9554139-9555738 | 99.64 | CBG14427 | CBG23216 |

| | | | | |
|---|---|---|---|---|
| III:9757716-9763607 | III:9764252-9770143 | 99.96 | CBG12647, CBG02546, CBG10121 | CBG02545, CBG02544, CBG25631, CBG30848, CBG10123, CBG03424 |
| III:10642341-10648145 | III:10635536-10641340 | 99.96 | CBG30842, CBG06793, Cbr-aqp-5 | CBG12848, CBG02308, CBG06795, Cbr-bus-4, CBG09600, Cbr-vps-41 |
| III:10613619-10620004 | III:10606233-10612618 | 99.84 | Cbr-ptp-5.3, CBG02313, Cbr-timm-17B.1, Cbr-hoe-1, Cbr-chw-1, CBG14647, CBG14648 | CBG02316, Cbr-nhl-1, CBG01746, Cbr-rpl-20, Cbr-cyc-2.1, CBG30616, CBG09606 |
| III:10573860-10576681 | III:9960308-9963129 | 99.94 | CBG02329, CBG27075 | CBG29867, CBG23095 |
| III:10157359-10162673 | III:10163074-10168388 | 99.98 | CBG25368, CBG06907, CBG24380, CBG09731, CBG27070, Cbr-sec-61.G | CBG02435, CBG06906, CBG24382, CBG24383, CBG09729 |
| III:10234616-10235954 | III:10236549-10237922 | 91.55 | Cbr-sdz-1.2 | Cbr-sdz-1.1 |
| III:10236550-10237922 | III:10234616-10235955 | 91.55 | Cbr-sdz-1.1 | Cbr-sdz-1.2 |
| III:10480496-10483611 | III:9956160-9959307 | 97.56 | CBG27727, Cbr-dpy-8 | Cbr-col-123, CBG23097 |
| III:9950187-9954039 | III:10462958-10466810 | 99.92 | CBG23099 | CBG12811, Cbr-pho-1, CBG14615 |
| III:9956159-9959307 | III:10480499-10483647 | 99.22 | Cbr-col-123, CBG23097 | CBG27727, Cbr-dpy-8 |
| III:10864685-10868502 | III:10868977-10872770 | 99.89 | CBG20614 | CBG20616, CBG03559, CBG27741 |
| III:11685663-11689846 | III:11689947-11694129 | 99.67 | CBG20144, CBG17682, CBG05016 | CBG31295, CBG09120, CBG27755 |
| III:11758538-11763590 | III:11765319-11770361 | 99.48 | CBG13645, CBG30986 | CBG17655, CBG17654, Cbr-cest-16 |
| III:11908810-11913422 | III:11916636-11921248 | 99.96 | Cbr-rbm-3.1, Cbr-sdz-15, Cbr-nas-33 | CBG18681, CBG18680, Cbr-nos-1 |
| III:12529827-12540959 | III:12542096-12553228 | 99.41 | CBG19622, Cbr-tyr-1, CBG27775 | CBG19621, CBG19620, CBG12272, CBG08369, CBG30957, CBG08368, CBG19039, Cbr-odc-1, CBG27777, CBG30805 |
| III:12676844-12686619 | III:12687621-12697405 | 99.94 | CBG19589, Cbr-fut-4, CBG30065, Cbr-gsto-1.1, Cbr-egl-21, Cbr-bgnt-1.1, CBG19078, Cbr-nhr-286, Cbr-nhr-135 | CBG18810, CBG26173, CBG19583, Cbr-gsto-1.2, Cbr-otpl-1, CBG11000 |
| III:13666809-13676171 | III:13685043-13694405 | 99.72 | CBG15379, CBG26207, Cbr-srx-97, CBG23012, Cbr-cyp-29A4, CBG19333 | CBG15382, CBG21425, Cbr-pitr-4, Cbr-sulp-8 |
| III:14396033-14403741 | III:14404742-14412450 | 99.8 | Cbr-ztf-30.2, CBG04555, CBG04556, Cbr-srh-17 | Cbr-impt-1, CBG08010, CBG10287, Cbr-ztf-30.1, Cbr-gasr-8 |

| | | | | |
|---|---|---|---|---|
| III:14513523-14521015 | III:14246586-14254078 | 99.73 | CBG08031, CBG26568, Cbr-acbp-4.2, CBG15352, CBG15353 | CBG10336, CBG00606, CBG00607, Cbr-acbp-4.1, CBG00609, CBG30888, CBG11604, CBG00082 |
| IV:1-4892 | IV:1491435-1496326 | 99.92 | Cbr-srv-16.2 | CBG05119, Cbr-srv-16.1, CBG27424 |
| IV:578437-584361 | IV:584485-590409 | 99.65 | CBG15159, CBG08170 | CBG00936 |
| IV:1302973-1306512 | IV:11674712-11678275 | 97.68 | Cbr-mesp-1.4, CBG13754 | CBG13626, Cbr-maea-1, Cbr-mesp-1.2, Cbr-zgpa-1, CBG05012 |
| IV:1486287-1490434 | IV:1589258-1593405 | 99.97 | CBG18719, CBG13798, CBG00797 | CBG27430 |
| IV:13351979-13353578 | IV:1612007-1613606 | 99.85 | CBG25419 | CBG31306, CBG27433 |
| IV:2135188-2138867 | IV:4022649-4026444 | 87.58 | CBG21960, CBG21587, Cbr-lbp-4 | CBG04213, Cbr-atg-10, Cbr-ceh-49.1, CBG01404 |
| IV:2135429-2136848 | IV:2144091-2145584 | 88.54 | CBG21587 | Cbr-ceh-49.3 |
| IV:2141648-2144090 | IV:4019497-4021943 | 97.87 | CBG21590 | CBG30428, CBG01406 |
| IV:2144086-2145798 | IV:4016992-4018701 | 92.51 | Cbr-ceh-49.3 | Cbr-ceh-49.6 |
| IV:2147784-2150512 | IV:4014234-4016992 | 93.1 | CBG30153, CBG21592, CBG20577, CBG16117 | CBG15046, CBG01407 |
| IV:2149919-2151085 | IV:2135607-2136753 | 85.39 | Cbr-ceh-49.5 | CBG21587 |
| IV:2342084-2348827 | IV:2334602-2341345 | 99.99 | CBG30494, CBG30495 | CBG25182, CBG11716, Cbr-str-33, CBG24588 |
| IV:2906617-2913515 | IV:2914002-2920900 | 99.65 | CBG30002, Cbr-srj-32, CBG22430, CBG22431, Cbr-acdh-10, Cbr-acdh-7 | Cbr-ptps-1, CBG22233, CBG25832, CBG22432, CBG22435, CBG15943 |
| IV:2883763-2891178 | IV:2875332-2882747 | 98.44 | CBG13423, CBG25829, CBG26939, CBG22421, Cbr-sdz-34.4 | CBG22223, CBG13424, CBG22357, CBG15951 |
| IV:11132948-11134166 | IV:6677825-6679035 | 99.1 | CBG09440 | CBG17252 |
| IV:3264186-3267906 | IV:3259773-3263493 | 99.59 | CBG13343 | CBG31484 |
| IV:3440110-3446474 | IV:3462423-3468787 | 99.76 | CBG23889, CBG24733, CBG01928, CBG01929, CBG01931 | Cbr-vha-10, CBG00725, CBG30502, CBG10453 |
| IV:3485721-3491767 | IV:3536409-3542455 | 99.48 | CBG00719, CBG17492, CBG31482, CBG01942 | Cbr-srg-1, CBG22665 |
| IV:3884873-3886206 | IV:3881240-3882573 | 98.25 | CBG15020 | CBG15019 |
| IV:3881240-3882573 | IV:3884873-3886206 | 98.35 | CBG15019 | CBG15020 |
| IV:4014317-4017015 | IV:2147783-2150512 | 93.1 | CBG15046, CBG01407 | CBG30153, CBG21592, CBG20577, CBG16117 |
| IV:4019568-4021944 | IV:2141648-2144091 | 97.43 | CBG01406 | CBG21590 |
| IV:4022711-4026444 | IV:2135148-2138868 | 87.55 | CBG04213, Cbr-atg-10, Cbr-ceh-49.1, CBG01404 | CBG21960, CBG21587, Cbr-lbp-4 |

| | | | | |
|---|---|---|---|---|
| IV:4824676-4831176 | IV:4831529-4838029 | 99.81 | Cbr-knl-1, Cbr-coa-4, CBG31532 | CBG31007, CBG31179, Cbr-srj-33, CBG27518, CBG30816 |
| IV:5103394-5106501 | IV:11462602-11465725 | 96.99 | Cbr-mcm-4 | Cbr-mrpl-48, Cbr-crn-2 |
| IV:5443607-5446884 | IV:5438440-5441717 | 99.9 | Cbr-gpx-1, Cbr-gnrr-7 | Cbr-cox-7C |
| IV:5562657-5566988 | IV:5691075-5695406 | 99.56 | CBG03773, CBG11185 | CBG03730, CBG03731, CBG17501 |
| IV:6012193-6017354 | IV:6019155-6024316 | 99.93 | Cbr-vacl-14, CBG29836, Cbr-cul-4, CBG29759, Cbr-cec-6, CBG06319 | CBG08223, Cbr-cnep-1, Cbr-mrt-2, CBG17572 |
| IV:6150338-6151836 | IV:6154331-6155758 | 92.98 | CBG19895 | CBG19894 |
| IV:6154334-6155788 | IV:6150335-6151810 | 92.46 | CBG19894 | CBG19895 |
| IV:6227912-6230057 | IV:6421797-6423912 | 98.69 | CBG08267 | CBG18179 |
| IV:6421799-6423909 | IV:6227920-6230058 | 99.81 | CBG18179 | CBG08267 |
| IV:6464113-6467538 | IV:6528469-6531894 | 99.98 | CBG30211 | CBG10807, CBG12958, Cbr-skr-17, CBG12956, CBG09465 |
| IV:6848043-6853757 | IV:6794784-6800449 | 98.19 | Cbr-pfd-6, CBG18064, Cbr-ipla-6.1 | Cbr-xpa-1, CBG03266, Cbr-ipla-6.2, CBG06540, CBG06541 |
| IV:7081315-7082588 | IV:7085439-7086739 | 88.99 | CBG30873, CBG05402 | CBG05403 |
| IV:7085448-7086739 | IV:7081315-7082602 | 89.52 | CBG05403 | CBG30873, CBG05402 |
| IV:8209935-8211451 | IV:4123483-4124790 | 99.77 | CBG14145 | CBG09047 |
| IV:8181097-8182682 | IV:6408545-6409915 | 82.41 | CBG05739 | CBG19838 |
| IV:7838392-7839970 | IV:7694083-7695703 | 92.03 | CBG05839, CBG05838 | CBG05886, CBG05885 |
| IV:7836556-7839988 | IV:7831202-7834634 | 99.12 | Cbr-cyn-4, CBG05840, CBG05839, CBG05838, CBG08666, Cbr-lin-32 | CBG05843, CBG05842, CBG27616 |
| IV:7836556-7838394 | IV:7695933-7697771 | 96.92 | CBG05840, CBG08666 | Cbr-col-39, CBG05884, CBG29117 |
| IV:7694107-7695703 | IV:7833038-7834634 | 99.12 | CBG05886, CBG05885 | CBG05842 |
| IV:7830908-7833040 | IV:7695933-7698228 | 96.92 | CBG05843 | Cbr-col-39, CBG05884, CBG05883, CBG29117 |
| IV:7696959-7698219 | IV:8460130-8461347 | 95.9 | CBG05884, CBG05883 | CBG06060, CBG06061 |
| IV:7696089-7698218 | IV:7830760-7832836 | 98.32 | Cbr-col-39, CBG05884, CBG05883, CBG29117 | CBG05844, CBG05843 |
| IV:7696089-7697780 | IV:7836556-7838191 | 98.42 | Cbr-col-39, CBG05884, CBG29117 | CBG05840 |
| IV:7833037-7834645 | IV:7838392-7839999 | 90.86 | CBG05842 | CBG05839, CBG05838 |
| IV:7592978-7595000 | IV:7590592-7592614 | 93.85 | Cbr-srg-25 | CBG12042, CBG16538, Cbr-kbp-1 |
| IV:7397568-7401079 | IV:8370163-8373674 | 99.81 | CBG11987, CBG29941, CBG11789 | CBG12206, Cbr-ttr-43 |

| | | | | |
|---|---|---|---|---|
| IV:8373898-8377633 | IV:7393399-7397134 | 99.88 | CBG09753, CBG30417 | CBG16488, CBG16489, CBG06702 |
| IV:7224967-7228549 | IV:8364722-8368304 | 99.95 | CBG06658 | CBG12207, CBG02869, Cbr-best-7.2 |
| IV:8439845-8444977 | IV:8452396-8457528 | 99.88 | CBG02845, CBG06055 | CBG06057, CBG06058, Cbr-clc-3 |
| IV:8444338-8444977 | IV:7694060-7694699 | 94.01 | CBG06055 | CBG05886 |
| IV:9188433-9190210 | IV:9195134-9196911 | 99.27 | CBG06246 | CBG09962, CBG06247 |
| IV:10258416-10264727 | IV:10119515-10125826 | 99.95 | Cbr-unc-32, CBG22318, CBG22319, CBG09704, CBG14567 | CBG12733, CBG12734, Cbr-ttc-4, Cbr-cdc-14.1, CBG24368, CBG24369, CBG09743, CBG30526, CBG27068 |
| IV:10157466-10158712 | IV:10159933-10161177 | 81.26 | CBG06907 | CBG27070 |
| IV:10159935-10161177 | IV:10157460-10158708 | 81.18 | CBG27070 | CBG06907 |
| IV:10285118-10289239 | IV:10310865-10314986 | 99.52 | CBG01845, Cbr-scl-15, Cbr-irld-6, Cbr-srh-55.2 | Cbr-clec-141, CBG06864, CBG06863, CBG01837 |
| IV:10314590-10315887 | IV:10312949-10314246 | 98.2 | CBG01836 | CBG06863, CBG01837 |
| IV:10520075-10521242 | IV:10521845-10523013 | 95.93 | CBG01768 | CBG01767 |
| IV:10521846-10523014 | IV:10520073-10521241 | 96.01 | CBG01767 | CBG01768 |
| IV:10306609-10308049 | IV:10729123-10730563 | 100 | CBG25644, CBG01838 | CBG01707 |
| IV:11051208-11052589 | IV:11054362-11055742 | 92.41 | CBG29880, CBG01630 | CBG01629 |
| IV:11054362-11055823 | IV:11051208-11052589 | 92.26 | CBG01629 | CBG29880, CBG01630 |
| IV:11138532-11139817 | IV:11598259-11599460 | 96.94 | CBG03615 | CBG20840 |
| IV:11339388-11346596 | IV:11347597-11354805 | 99.99 | CBG08476, CBG08475, Cbr-kin-5.1, Cbr-nlp-10, CBG31347, CBG09397, Cbr-inx-3.1, CBG14795, CBG14796 | CBG20777, Cbr-dmsr-7, CBG09394, CBG09393, CBG14797, CBG14798 |
| IV:11462602-11465725 | IV:5103393-5106502 | 96.83 | Cbr-mrpl-48, Cbr-crn-2 | Cbr-mcm-4 |
| IV:11598257-11599461 | IV:11138552-11139755 | 96.94 | CBG20840 | CBG03615 |
| IV:13738028-13739370 | IV:13652242-13653583 | 97.18 | CBG18475 | CBG18462 |
| IV:14199703-14203180 | IV:6748366-6751865 | 98.17 | CBG10348, Cbr-irld-14, Cbr-str-124 | CBG29843 |

| | | | | |
|---|---|---|---|---|
| IV:14204214-14205410 | IV:14669012-14670261 | 94.4 | CBG03512 | CBG16471, Cbr-rpoa-12 |
| IV:14373173-14378476 | IV:14378577-14383880 | 99.87 | CBG10294, Cbr-pssy-2, CBG04548 | CBG10292, Cbr-oig-3, Cbr-unc-61 |
| IV:15073694-15080663 | IV:15065737-15072710 | 99.93 | Cbr-cht-2 | Cbr-mrps-18B |
| IV:15864423-15868944 | IV:15869987-15874508 | 99.31 | CBG07153, CBG31454, Cbr-srd-21 | CBG13950, CBG04993, CBG15449 |
| IV:16334530-16337091 | IV:16447608-16450174 | 99.69 | CBG07048 | CBG30392, CBG23981 |
| IV:16632589-16638046 | IV:16594964-16600421 | 98.27 | CBG30987, CBG27998 | CBG06960, CBG10577, Cbr-vamp-7, CBG24406 |
| IV:17221319-17225744 | IV:17212941-17217366 | 99.73 | CBG26539, CBG30908 | CBG13515 |
| V:599181-600665 | V:623616-625191 | 80.69 | CBG08173 | CBG08179 |
| V:999067-1003150 | V:1003284-1007367 | 99.9 | CBG13705, CBG16370 | Cbr-sdhb-1, CBG30300, CBG13706 |
| V:1250682-1251830 | V:1239196-1240370 | 98.89 | CBG01506, CBG01507 | CBG26907, CBG01498 |
| V:1248721-1249878 | V:1241031-1242217 | 97.48 | CBG01504, Cbr-his-63 | CBG01499, CBG01500 |
| V:1241052-1242163 | V:1248749-1249880 | 98.42 | CBG01499, CBG01500 | CBG01504, Cbr-his-63 |
| V:1239217-1240366 | V:1250659-1251832 | 98.89 | CBG26907, CBG01498 | CBG01506, CBG01507 |
| V:1615509-1617177 | V:6736547-6738217 | 96.99 | CBG27434 | CBG06519 |
| V:1793491-1798178 | V:1799932-1804619 | 99.34 | Cbr-tufm-1, CBG30329, CBG26240, Cbr-nmat-1 | Cbr-rps-18 |
| V:11475297-11476910 | V:7473277-7474891 | 98.39 | CBG26451 | CBG26804 |
| V:2002577-2006499 | V:4429198-4433114 | 98.24 | CBG21557, CBG31291, CBG20545 | Cbr-cytb-5.2, Cbr-dhod-1, CBG01282, CBG31338, CBG27508 |
| V:2591538-2598335 | V:2647224-2654021 | 99.9 | CBG13477, CBG21823 | Cbr-clec-137, CBG13463, CBG21836, CBG21837, CBG15990 |
| V:2726539-2730247 | V:2479336-2483044 | 99.91 | CBG26113, Cbr-srbc-59 | CBG31243 |
| V:3152977-3157304 | V:3148108-3152435 | 99.6 | CBG17014 | CBG27473 |
| V:3053644-3057499 | V:3049778-3053633 | 99.88 | CBG22265, CBG13383, CBG13382, CBG31198, CBG16979, CBG31269 | CBG16978 |
| V:3308773-3310341 | V:7286915-7288488 | 98.6 | CBG13335 | CBG30215, CBG17135 |
| V:3484959-3486836 | V:3490509-3492386 | 99.73 | CBG10446 | CBG10445 |
| V:3717153-3720587 | V:3551416-3554850 | 99.99 | CBG01988, CBG30710, CBG01989, CBG27487 | Cbr-wdr-12, CBG10431, CBG10430 |
| V:3576941-3578198 | V:3579150-3580410 | 95.75 | CBG10424 | CBG10423 |
| V:3579150-3580409 | V:3576941-3578199 | 95.83 | CBG10423 | CBG10424 |
| V:4256370-4257436 | V:4261518-4262553 | 84.98 | CBG00569 | CBG01340 |

| | | | | |
|---|---|---|---|---|
| V:4429253-4432344 | V:2003348-2006498 | 95.4 | Cbr-cytb-5.2, CBG01282, CBG31338, CBG27508 | CBG21557, CBG31291, CBG20545 |
| V:4534984-4539853 | V:4543559-4548439 | 97.59 | CBG30453, CBG01247, Cbr-nhr-190.1 | CBG21747, CBG01244 |
| V:4710542-4711672 | V:4732294-4733424 | 97.62 | CBG01197 | CBG01188 |
| V:4727914-4729992 | V:4714693-4716780 | 96.69 | CBG01191 | CBG01195, CBG01194 |
| V:1591124-1595255 | V:7018817-7022939 | 98.87 | CBG15784 | Cbr-mab-21, CBG26374 |
| V:4963465-4967087 | V:4748484-4752106 | 99.2 | CBG03918, CBG29829, Cbr-hsp-12.3 | CBG25242, CBG21719 |
| V:4986537-4988441 | V:4991276-4993180 | 98.53 | CBG01114 | CBG03913, CBG01112 |
| V:5443652-5448955 | V:5493285-5498587 | 99.27 | Cbr-gpx-1, CBG20035, Cbr-gnrr-7 | Cbr-nubp-1, CBG30963, CBG18258, Cbr-elo-7, CBG27003 |
| V:5553517-5555138 | V:5548397-5550061 | 96 | Cbr-nhr-247.2 | CBG03776, Cbr-nhr-247.1 |
| V:5548677-5550060 | V:5553517-5554908 | 98.64 | CBG03776, Cbr-nhr-247.1 | Cbr-nhr-247.2 |
| V:5712617-5717525 | V:5877267-5882132 | 98.66 | CBG25223, Cbr-marc-3, CBG29835, CBG17888, Cbr-lpr-5 | CBG26570, CBG13096, CBG13192, Cbr-pepm-1.1, CBG30594, CBG17941 |
| V:5873554-5874720 | V:5882233-5883412 | 99.07 | CBG17939 | CBG06277 |
| V:6097873-6101801 | V:7009895-7013828 | 99.97 | CBG06338 | Cbr-erh-2, CBG05425, CBG06602 |
| V:6102339-6106551 | V:7013985-7018197 | 99.88 | CBG13251, CBG19910, CBG06339, CBG06340 | Cbr-asb-1, Cbr-srbc-36, CBG06605 |
| V:6632625-6634060 | V:6630383-6631817 | 99.79 | CBG06483 | CBG06482 |
| V:6692003-6699160 | V:6699653-6706810 | 99.65 | CBG20490, CBG03287, CBG30480, CBG05511, CBG30902, CBG06506 | CBG20487, Cbr-best-23, CBG06508, CBG30903, CBG06509, CBG31518 |
| V:6792053-6797063 | V:17754576-17759614 | 96.77 | CBG05481, Cbr-srw-103, CBG06539, CBG17232 | CBG17824 |
| V:6798082-6807842 | V:17742713-17752578 | 96.99 | Cbr-crml-1, Cbr-dnj-23, Cbr-dyrb-1, CBG03263, Cbr-txt-9.2, Cbr-mnp-1, CBG26378, Cbr-ops-1, CBG06541, CBG06542, Cbr-gur-3 | CBG17823, CBG30845, CBG30844, Cbr-srw-101.1 |
| V:6805543-6808207 | V:17742351-17745029 | 96.69 | Cbr-ops-1 | CBG17823 |
| V:6828374-6830854 | V:17738590-17741140 | 95.12 | CBG06553 | CBG17821 |
| V:7000583-7008876 | V:7024096-7032389 | 99.86 | Cbr-col-61, CBG18035, CBG30448, CBG31384, Cbr-ddi-1, CBG05426, CBG06598, CBG06599, CBG06600, CBG06601 | CBG21944, CBG11902, CBG11903, Cbr-fan-1, Cbr-paqr-1, CBG06608, CBG06609, CBG06610, Cbr-nhr-178 |
| V:7069686-7079545 | V:7503125-7512985 | 99.69 | CBG03194, Cbr-odr-8, CBG05409, Cbr-dnj-11, | Cbr-glh-4, CBG25567, CBG03087, CBG03086, CBG16518, Cbr-tofu-7 |

| | | | | |
|---|---|---|---|---|
| | | | Cbr-lec-11, CBG05406, CBG17182 | |
| V:7286918-7288488 | V:3308773-3310341 | 98.54 | CBG30215, CBG17135 | CBG13335 |
| V:7473275-7474892 | V:5830803-5832411 | 99.38 | CBG26804 | CBG25854 |
| V:7496549-7497800 | V:12647783-12649040 | 99.44 | Cbr-tmem-17 | CBG10990 |
| V:7796605-7800344 | V:7808914-7812661 | 99.92 | CBG12082, Cbr-srab-3.2, CBG14047 | CBG12086, CBG25575, CBG25932, Cbr-srab-3.1 |
| V:7802613-7808915 | V:7812923-7819227 | 99.92 | Cbr-aars-2, CBG31425, Cbr-crn-4 | CBG12088, Cbr-mrps-6, CBG03007, CBG05848, CBG08671 |
| V:7839233-7840526 | V:3635518-3636828 | 99.01 | CBG05838 | Cbr-col-176 |
| V:8035783-8039182 | V:8323384-8326783 | 99.97 | CBG25104, CBG25105, Cbr-mutd-1, CBG08604, CBG31468 | Cbr-wdr-5.1, CBG24764, CBG24765 |
| V:8688628-8690761 | V:8696394-8698533 | 98.98 | CBG23445, CBG23444 | Cbr-col-86, CBG23437, CBG23436 |
| V:8690871-8692346 | V:8694903-8696394 | 97.39 | CBG23443, CBG23442 | CBG23438 |
| V:8694917-8696395 | V:8690860-8692345 | 98.26 | CBG23438 | CBG23443, CBG23442 |
| V:8696407-8698533 | V:8688617-8690762 | 98.79 | Cbr-col-86, CBG23437, CBG23436 | CBG23445, CBG23444 |
| V:9396376-9399738 | V:9402167-9405544 | 99.97 | Cbr-unc-120 | CBG10027 |
| V:9399708-9402130 | V:9409565-9411987 | 99.93 | Cbr-mpst-1.2 | CBG29920, Cbr-set-4, CBG03338, CBG03339, Cbr-mpst-1.1 |
| V:9549597-9550606 | V:9646412-9647413 | 99.6 | Cbr-col-157.2 | Cbr-col-157.1 |
| V:9760395-9763760 | V:8340151-8343516 | 94.88 | CBG02546 | CBG12210, CBG02877 |
| V:8340162-8342434 | V:8343618-8345890 | 95.12 | CBG02877 | CBG02875 |
| V:8353245-8358744 | V:9763771-9769270 | 99.96 | CBG05701, CBG27022, CBG31220 | CBG02545, CBG02544, CBG30848, CBG10123, CBG03424, CBG30653 |
| V:10121964-10125612 | V:10159335-10162983 | 99.87 | CBG09743, CBG30526, CBG27068 | CBG24380, Cbr-dod-24, CBG27070, Cbr-sec-61.G |
| V:10265605-10266901 | V:14102969-14104290 | 93.06 | CBG09703 | CBG11554 |
| V:11052533-11060615 | V:11060716-11068798 | 99.98 | CBG20245, CBG20667, Cbr-gst-12, CBG01629, CBG01628, CBG01627, CBG29115 | Cbr-gst-15, Cbr-gst-24, Cbr-gst-20, Cbr-csc-1, Cbr-fbxc-23, CBG03602, CBG01626, CBG01624, CBG26446 |
| V:11134458-11140118 | V:11128430-11134089 | 99.95 | CBG03615, CBG14754 | CBG08517, Cbr-max-1.2, CBG09440 |
| V:12476774-12480759 | V:12482518-12486503 | 99.52 | CBG20423, CBG19636, CBG12261, CBG19023 | CBG30245, CBG26494, Cbr-dmsr-11, Cbr-cah-5 |
| V:12600724-12604044 | V:13133560-13136877 | 99.91 | CBG26489 | CBG04336 |

| | | | | |
|---|---|---|---|---|
| V:12604039-12614128 | V:13136872-13146956 | 99.76 | CBG18795, Cbr-nhr-88, CBG12283, CBG26488, CBG30366, CBG30367, CBG19050, CBG19052, CBG19053, Cbr-alh-10 | CBG15859, CBG15858, CBG15857, CBG04338, CBG25522, Cbr-gop-2, Cbr-col-103, CBG26472, CBG19209, Cbr-srx-16.2, Cbr-srx-16.1, Cbr-pcyt-1, CBG16734 |
| V:12792386-12798751 | V:12803727-12810092 | 99.78 | Cbr-set-11, CBG19561, CBG27169, Cbr-hpo-9, Cbr-hyls-1.1, Cbr-spp-2, Cbr-spp-5, CBG27790, CBG27791 | CBG26597, CBG19559, CBG19558, CBG19557, CBG21241, CBG19115, CBG19116, Cbr-hyls-1.2 |
| V:13435507-13438945 | V:14044466-14047904 | 99.13 | CBG19274 | CBG19434 |
| V:14075721-14077736 | V:14533141-14535156 | 96.95 | CBG30297 | CBG24814 |
| V:14284794-14286909 | V:14364904-14366860 | 86.38 | CBG11614 | CBG10301, CBG04543 |
| V:14364989-14366831 | V:14284809-14286878 | 85.28 | CBG10301, CBG04543 | CBG11614 |
| V:14581636-14583534 | V:14635555-14637499 | 93.79 | CBG04591 | CBG04607, CBG04608 |
| V:14635538-14637610 | V:14581641-14583613 | 94.8 | CBG04607, CBG04608 | CBG04591, CBG04592 |
| V:15337441-15338195 | V:15335630-15336384 | 98.76 | CBG04806 | CBG04804 |
| V:14928785-14929720 | V:14904066-14905001 | 95.86 | CBG04710 | CBG04696 |
| V:14905700-14908099 | V:14902697-14905095 | 97.33 | CBG04698, CBG00220 | CBG04695, CBG04696 |
| V:14902678-14905095 | V:14905698-14908098 | 96.42 | CBG04695, CBG04696 | CBG04698, CBG00220 |
| V:15595271-15598316 | V:15521456-15524559 | 98.11 | CBG04880 | CBG04857, CBG31263 |
| V:15599123-15600191 | V:15644224-15645317 | 95.26 | CBG04882, CBG30796 | CBG04902 |
| V:15644224-15645318 | V:15519513-15520607 | 95.75 | CBG04902 | CBG04855 |
| V:15511106-15517027 | V:15602224-15608145 | 97.43 | CBG23572, CBG04852, CBG04853, CBG15530 | CBG30089, CBG04884, CBG04885 |
| V:15512562-15515972 | V:15640702-15644116 | 97 | CBG04852, CBG04853, CBG15530 | CBG04904 |
| V:15695416-15698154 | V:15704415-15707157 | 96.03 | CBG30090, CBG04923, CBG27260 | CBG04926, CBG31266, CBG27968 |
| V:15782678-15784028 | V:15784308-15785658 | 97.2 | CBG04955 | CBG04956 |
| V:15784333-15785704 | V:15782674-15784028 | 97.3 | CBG04956 | CBG04955 |
| V:16560669-16561924 | V:16567634-16568855 | 81.47 | CBG16823, CBG24668 | CBG24669 |
| V:16577531-16582375 | V:16571697-16576541 | 99.13 | Cbr-nstp-4, CBG22558, CBG24401 | CBG24357, CBG22733 |
| V:16662858-16669610 | V:16746600-16753359 | 99.87 | CBG30395, CBG22717 | CBG24530, CBG22701 |
| V:16754883-16756647 | V:16761860-16763588 | 94.75 | CBG22915 | CBG22913 |
| V:16761860-16763584 | V:16754923-16756649 | 94.81 | CBG22913 | CBG22915 |
| V:16852460-16853928 | V:16885006-16886475 | 97.89 | CBG22885 | CBG22876 |
| V:16873731-16877972 | V:16920531-16924772 | 98.54 | CBG22880, CBG22879 | CBG22500, CBG22501, CBG22867, CBG05232 |
| V:16924286-16927858 | V:16877021-16880659 | 99.31 | CBG22865, CBG05233 | CBG22878, CBG22877 |

| | | | | |
|---|---|---|---|---|
| V:16925393-16929958 | V:16930960-16935525 | 99.16 | CBG22865, CBG22864, CBG22863 | CBG26532, CBG22862, CBG22861, CBG22860, CBG05235 |
| V:16877021-16879486 | V:16930960-16933425 | 99.16 | CBG22878 | CBG22862 |
| V:16971167-16972718 | V:16972967-16974518 | 96 | CBG22843 | CBG22842 |
| V:16998066-17001663 | V:17016446-17020043 | 97.96 | CBG30398, CBG31228, CBG31227 | CBG22516, CBG22832 |
| V:17194835-17197015 | V:17217104-17219283 | 98.44 | CBG18656, CBG28021 | CBG18650 |
| V:17197171-17200946 | V:17219379-17223154 | 99.17 | CBG13511 | CBG26539, CBG18649 |
| V:17252822-17257888 | V:17257889-17262955 | 99.92 | Cbr-ham-1, Cbr-str-96.1, CBG28028, CBG05310 | CBG18638, Cbr-str-96.2, CBG28029 |
| V:17409522-17411096 | V:11475297-11476915 | 98.64 | Cbr-ced-2 | CBG26451 |
| V:17738592-17741135 | V:6828360-6830856 | 95.28 | CBG17821 | CBG06553 |
| V:17742353-17746622 | V:6803938-6808211 | 97.56 | CBG17823, CBG30845 | Cbr-clh-5, Cbr-ops-1 |
| V:6798057-6803886 | V:17746736-17752565 | 99.21 | Cbr-dnj-23, Cbr-dyrb-1, Cbr-txt-9.2, CBG26378, CBG06541, Cbr-gur-3 | CBG30844, Cbr-srw-101.1 |
| V:17754557-17759631 | V:6792052-6797090 | 96.75 | CBG17824 | CBG05481, Cbr-srw-103, CBG06539, CBG17232 |
| V:18073829-18080269 | V:18081971-18088411 | 99.88 | CBG23817, CBG23818, CBG07673 | CBG27335, CBG23820, CBG07671 |
| V:19227783-19232321 | V:19232520-19237058 | 99.92 | Cbr-snx-6 | CBG31387, CBG28079, Cbr-cdk-4 |
| V:19348264-19355264 | V:19356265-19363265 | 99.91 | Cbr-str-93 | CBG05675, Cbr-nlp-53, CBG07404 |
| X:655838-664603 | X:591827-600592 | 99.64 | CBG11342, CBG26883, Cbr-ife-2.2 | Cbr-dpl-1, CBG13652, CBG11356, CBG31495, Cbr-ife-2.1, CBG30827, CBG08173 |
| X:470451-474130 | X:1436403-1440055 | 98.03 | CBG15181, CBG11389 | Cbr-cutl-20, CBG26913, CBG00777, CBG16276 |
| X:1436403-1440049 | X:6856935-6860571 | 98.44 | Cbr-cutl-20, CBG26913, CBG00777, CBG16276 | CBG25467, CBG21906, Cbr-nex-3, Cbr-cnc-11, Cbr-cnc-4 |
| X:160068-161678 | X:5494157-5495765 | 95.77 | Cbr-arrd-4 | CBG30963 |
| X:1111674-1119550 | X:927943-935819 | 99 | CBG29802, CBG29803, CBG29804, Cbr-cus-2, Cbr-nlp-63, CBG01464, CBG31299, Cbr-acd-4, CBG16347 | Cbr-strm-1, CBG13699, Cbr-gpa-2 |
| X:1436127-1440008 | X:6856933-6860845 | 98.12 | Cbr-cutl-20, CBG00775, CBG26913, CBG16276 | CBG25467, CBG21906, Cbr-nex-3, Cbr-cnc-11, Cbr-cnc-4, Cbr-cnc-3 |
| X:1436403-1440013 | X:470479-474129 | 97.96 | Cbr-cutl-20, CBG26913, CBG16276 | CBG15181, CBG11389 |

| | | | | |
|---|---|---|---|---|
| X:1766561-1771904 | X:1771915-1777258 | 99.03 | CBG22016 | CBG18442 |
| X:2370428-2376403 | X:2376575-2382550 | 99.92 | Cbr-elc-1, CBG11706, CBG23597, CBG23599, CBG16068 | CBG23602, CBG23603, CBG23605, CBG16067 |
| X:2437112-2444649 | X:2445650-2453187 | 99.78 | Cbr-bpnt-1, Cbr-pstk-1, CBG18739, CBG26260, CBG31211 | CBG31214, CBG31215, CBG31216, CBG16054 |
| X:3047844-3049586 | X:3045681-3047423 | 91.69 | CBG16977 | CBG22262 |
| X:3443033-3445309 | X:3445816-3448095 | 99.87 | CBG01929, CBG01931 | CBG14897, CBG01932, CBG01933 |
| X:3445814-3448089 | X:3443036-3445316 | 99.78 | CBG14897, CBG01932, CBG01933 | CBG01929, CBG01931 |
| X:3714048-3716038 | X:3721080-3723070 | 99.55 | CBG30709 | CBG19690 |
| X:3721082-3723080 | X:3714049-3716038 | 99.95 | CBG19690 | CBG30709 |
| X:4595454-4598017 | X:4590125-4592679 | 96.88 | Cbr-elof-1, CBG02162 | CBG11101, CBG02161 |
| X:4740312-4741373 | X:4738999-4740060 | 97.87 | CBG26290 | CBG01185 |
| X:4899445-4905960 | X:4906961-4913476 | 98.92 | CBG30895 | CBG03932, CBG01134 |
| X:5122661-5126341 | X:5127342-5131022 | 99.49 | Cbr-tbca-1, Cbr-hpr-9 | CBG27527 |
| X:5494141-5495683 | X:160065-161678 | 95.4 | CBG30963 | Cbr-arrd-4 |
| X:5697376-5702158 | X:5664184-5668965 | 98.62 | Cbr-snpn-1, CBG30837, CBG17503 | CBG26416, CBG17881, CBG17448, CBG31519 |
| X:6289151-6292122 | X:6293261-6296232 | 99.63 | CBG04526 | CBG04525 |
| X:6618586-6620661 | X:6698713-6700787 | 99.38 | CBG26631, CBG18125, CBG31402 | CBG29924, CBG06508 |
| X:6618059-6620664 | X:6698713-6701318 | 99.61 | CBG26631, CBG18125, CBG31402, CBG17262 | CBG29924, CBG20487, CBG06508 |
| X:6856874-6860876 | X:1436127-1440050 | 98.09 | CBG25467, CBG21906, Cbr-nex-3, Cbr-cnc-5.1, Cbr-cnc-11, Cbr-cnc-4, Cbr-cnc-3 | Cbr-cutl-20, CBG00775, CBG26913, CBG00777, CBG16276 |
| X:7259860-7262600 | X:7265937-7268677 | 97.58 | Cbr-clec-156.2, CBG17143, CBG17142 | CBG11950, CBG17140, CBG17139 |
| X:7256685-7261799 | X:7262714-7267881 | 98.88 | CBG31491, Cbr-clec-156.1, Cbr-clec-156.2, CBG17143, CBG17142 | CBG11949, CBG03138, CBG17140, CBG17139 |
| X:6681205-6684273 | X:7288073-7291259 | 99.28 | CBG31260 | CBG17956, CBG17135 |
| X:7373724-7376602 | X:7284195-7287072 | 99.13 | Cbr-abf-1 | Cbr-spo-11 |
| X:7515512-7517699 | X:7440018-7442180 | 96.44 | CBG26807 | CBG03104, CBG11803 |
| X:7514141-7517487 | X:7497578-7500925 | 99.88 | CBG03083, CBG26807 | Cbr-lin-37, Cbr-flp-13, CBG11821, CBG11822 |
| X:7440202-7442180 | X:7498947-7500925 | 99.88 | CBG03104, CBG11803 | CBG11822 |

| | | | | |
|---|---|---|---|---|
| X:7449732-7454768 | X:7443695-7448731 | 99.72 | Cbr-pmlr-1, Cbr-clec-143.2, CBG05958, Cbr-cutl-1, CBG11808 | CBG29846, CBG11998, CBG29847, CBG03103, CBG06720, CBG11805 |
| X:7498950-7500925 | X:7515510-7517485 | 97.4 | CBG11822 | CBG26807 |
| X:7503651-7506695 | X:7388016-7391057 | 99.87 | CBG03087 | CBG05975 |
| X:7619401-7620961 | X:7625688-7627251 | 93.87 | CBG27610 | CBG03052 |
| X:7625690-7627251 | X:7619401-7620960 | 94.12 | CBG03052 | CBG27610 |
| X:7748515-7752456 | X:9550084-9554025 | 99.75 | Cbr-npp-19, CBG14035, CBG14036 | CBG30838, Cbr-rbmx-2, Cbr-cpb-2, CBG10073, CBG27058, CBG23217, CBG14426, CBG14427 |
| X:9646665-9655132 | X:7765137-7773604 | 99.96 | Cbr-cuti-1, CBG12616, Cbr-rbbp-5, CBG10096, CBG10097, CBG03396, CBG23184, CBG14454 | CBG31426, Cbr-srh-130, CBG27004, Cbr-srh-129, CBG14040 |
| X:8271266-8276985 | X:8282749-8288468 | 99.95 | CBG12234, CBG02903, CBG31023 | CBG12230, CBG25291, CBG02900, CBG26367, Cbr-egg-1 |
| X:8365343-8367845 | X:8356871-8359369 | 99.2 | CBG12207 | CBG31220 |
| X:8483578-8487333 | X:8685095-8688858 | 96.13 | CBG29852, CBG29853, Cbr-gspd-1 | Cbr-srh-16 |
| X:8685090-8688854 | X:8483561-8487320 | 96.49 | Cbr-srh-16 | CBG29852, CBG29853, Cbr-gspd-1 |
| X:9369751-9372074 | X:9656133-9658448 | 100 | CBG12535 | Cbr-nlp-48, CBG23182 |
| X:9487031-9490039 | X:9491391-9494392 | 99.87 | CBG25907, Cbr-vrp-1, CBG27678 | Cbr-blos-4, CBG10053, CBG14412 |
| X:9491382-9494391 | X:9487041-9490040 | 99.87 | Cbr-blos-4, CBG10053, CBG14412 | CBG25907, Cbr-vrp-1, CBG27678 |
| X:9856224-9863089 | X:9867880-9874745 | 99.96 | CBG02525, CBG14491, Cbr-amt-4.1 | CBG02522, Cbr-dhs-12, Cbr-gst-2, Cbr-gst-4, CBG03453, CBG14493 |
| X:10156562-10160143 | X:10588151-10591732 | 99.89 | CBG25368, CBG06907, CBG24379 | CBG30140, Cbr-srd-10 |
| X:10590363-10594065 | X:10603200-10606902 | 99.6 | CBG25653 | Cbr-irld-34, CBG01748 |
| X:10585966-10587150 | X:10615451-10616635 | 99.94 | CBG02324, CBG14645 | CBG14647 |
| X:10851780-10859904 | X:11347480-11355604 | 99.95 | CBG12895, Cbr-nep-12, Cbr-rpl-7A, Cbr-tald-1, Cbr-srsx-30, Cbr-srsx-29, CBG14696, CBG14697 | CBG20777, Cbr-rei-1, Cbr-dmsr-7, CBG09394, CBG09393, CBG30687, CBG14797, CBG14798 |
| X:11136277-11137206 | X:11140063-11140992 | 99.68 | CBG14754 | CBG14755 |
| X:11340033-11346458 | X:11416919-11423344 | 99.88 | CBG08476, CBG08475, Cbr-kin-5.1, Cbr-nlp-10, CBG31347, Cbr-inx-3.1, CBG14795 | CBG25399, CBG25676, CBG30239, CBG20201, CBG09375, Cbr-cox-16, Cbr-inx-3.2, CBG14812 |
| X:12756679-12761394 | X:12761613-12766328 | 99.84 | CBG30851, CBG19101 | CBG08316 |

| | | | | |
|---|---|---|---|---|
| X:13349711-13353353 | X:13353440-13357082 | 99.86 | CBG29972, CBG17758 | CBG17755 |
| X:14045022-14047859 | X:13547940-13550777 | 97.46 | CBG19434 | CBG26193, CBG19306, Cbr-nstp-10 |
| X:15027347-15032984 | X:15033985-15039622 | 99.95 | CBG21494, CBG00244 | Cbr-str-52, CBG00245 |
| X:15839420-15840722 | X:15840923-15842249 | 91.29 | CBG30555, CBG15457 | CBG15456 |
| X:15840932-15842249 | X:15839411-15840722 | 91.14 | CBG15456 | CBG30555, CBG15457 |
| X:16808152-16809399 | X:16816156-16817403 | 99.05 | CBG22684 | CBG27301, CBG22683 |
| X:17268644-17272126 | X:17274419-17277868 | 99.65 | Cbr-manf-1 | Cbr-dml-1, CBG18632, CBG05315 |
| X:17274061-17277870 | X:17268677-17272484 | 99.71 | Cbr-dml-1, CBG18632, CBG05315 | Cbr-manf-1 |
| X:18945464-18947851 | X:19079583-19081966 | 99.71 | CBG07494 | CBG05615, CBG07468 |
| X:19561527-19563294 | X:13028350-13030054 | 95.98 | CBG07374 | CBG30545, CBG16706 |
| X:20776916-20778335 | X:20568522-20569942 | 90.13 | CBG10263 | CBG10217 |
| X:20681624-20683631 | X:21078696-21080710 | 98.66 | CBG10247 | CBG08890 |
| X:20568523-20570003 | X:20776909-20778314 | 90.76 | CBG10217 | CBG10263 |
| X:20565780-20569732 | X:20557825-20561777 | 98.13 | CBG10216, CBG10217 | CBG10213 |
| X:20557825-20561793 | X:20565780-20569729 | 97.7 | CBG10213 | CBG10216, CBG10217 |
| X:21115273-21117755 | X:20882283-20884765 | 99.76 | CBG08881 | CBG08942 |
| X:21078697-21080797 | X:20681620-20683632 | 98.61 | CBG08890 | CBG10247 |
| X:21169837-21171898 | X:21537391-21539452 | 97.67 | CBG10674 | CBG10579 |
| X:21149080-21156938 | X:21189823-21197681 | 99.95 | CBG10681, CBG10680, CBG31485 | CBG10666, CBG10665, CBG10664 |
| X:21438766-21440732 | X:21444674-21446639 | 99.75 | CBG10602, CBG10601 | CBG10598, CBG10597 |
| X:21443123-21444679 | X:21440831-21442387 | 100 | CBG10599 | CBG10600 |
| X:21445506-21446609 | X:21438767-21439898 | 99.91 | CBG10597 | CBG10602 |
